# Crucial neuroprotective roles of the metabolite BH4 in dopaminergic neurons

**DOI:** 10.1101/2023.05.08.539795

**Authors:** Shane J. F. Cronin, Weonjin Yu, Ashley Hale, Simon Licht-Mayer, Mark J Crabtree, Joanna A. Korecka, Evgenii O. Tretiakov, Marco Sealey-Cardona, Mate Somlyay, Masahiro Onji, Meilin An, Jesse D. Fox, Bruna Lenfers Turnes, Carlos Gomez-Diaz, Débora da Luz Scheffer, Domagoj Cikes, Vanja Nagy, Adelheid Weidinger, Alexandra Wolf, Harald Reither, Antoine Chabloz, Anoop Kavirayani, Shuan Rao, Nick Andrews, Alban Latremoliere, Michael Costigan, Gillian Douglas, Fernando Cini Freitas, Christian Pifl, Roger Walz, Robert Konrat, Don J. Mahad, Andrey V. Koslov, Alexandra Latini, Ole Isacson, Tibor Harkany, Penelope J. Hallett, Stefan Bagby, Clifford J. Woolf, Keith M. Channon, Hyunsoo Shawn Je, Josef M. Penninger

## Abstract

Dopa-responsive dystonia (DRD) and Parkinson’s disease (PD) are movement disorders caused by the dysfunction of nigrostriatal dopaminergic neurons. Identifying druggable pathways and biomarkers for guiding therapies is crucial due to the debilitating nature of these disorders. Recent genetic studies have identified variants of GTP cyclohydrolase-1 (GCH1), the rate-limiting enzyme in tetrahydrobiopterin (BH4) synthesis, as causative for these movement disorders. Here, we show that genetic and pharmacological inhibition of BH4 synthesis in mice and human midbrain-like organoids accurately recapitulates motor, behavioral and biochemical characteristics of these human diseases, with severity of the phenotype correlating with extent of BH4 deficiency. We also show that BH4 deficiency increases sensitivities to several PD-related stressors in mice and PD human cells, resulting in worse behavioral and physiological outcomes. Conversely, genetic and pharmacological augmentation of BH4 protects mice from genetically- and chemically induced PD-related stressors. Importantly, increasing BH4 levels also protects primary cells from PD-affected individuals and human midbrain-like organoids (hMLOs) from these stressors. Mechanistically, BH4 not only serves as an essential cofactor for dopamine synthesis, but also independently regulates tyrosine hydroxylase levels, protects against ferroptosis, scavenges mitochondrial ROS, maintains neuronal excitability and promotes mitochondrial ATP production, thereby enhancing mitochondrial fitness and cellular respiration in multiple preclinical PD animal models, human dopaminergic midbrain-like organoids and primary cells from PD-affected individuals. Our findings pinpoint the BH4 pathway as a key metabolic program at the intersection of multiple protective mechanisms for the health and function of midbrain dopaminergic neurons, identifying it as a potential therapeutic target for PD.

## Introduction

The metabolite tetrahydrobiopterin (BH4) is an essential co-factor for enzymes involved in various cellular processes, including nitric oxide production, ether lipid as well as phenylalanine metabolism, and neurotransmitter biosynthesis (Werner et al., 2011). Inherited BH4 deficiencies are a rare heterogeneous group of pediatric neurometabolic disorders, characterized by a range of symptoms such as poor suckling, trunk hypotonia, hypertonia of limbs, difficulty in swallowing, microcephaly, seizures, or intellectual disabilities (Blau et al., 1996; Opladen et al., 2020). BH4 deficiency also affects dopamine synthesis in dopaminergic (DAergic) neurons and has been implicated in movement disorders such as dopa-responsive dystonia (DRD; OMIM:128230) and Parkinson’s disease (PD; OMIM 168600).

DRD is a group of rare movement disorders caused by genetic deficiencies in enzymes controlling dopamine synthesis, and it can be treated with L-Dopa (Furukawa, 1993; Van Hove et al., 2006; Segawa et al., 2013). One of the most studied forms of DRD, Segawa disease, is caused by autosomal dominant inherited mutations in the enzyme GTP cyclohydrolase (GTPCH; EC 3.5.4.16 hereafter referred to as GCH1) (Segawa et al., 1976) (**Figure S1A**), which is involved in BH4 biosynthesis. Recent studies have also linked *GCH1* mutations PD, a progressive, neurodegenerative condition characterized clinically by motor and cognitive symptoms (Mencacci et al., 2014; Nalls et al., 2019; Rudakou et al., 2019a; Xu et al., 2017; Yoshino et al., 2018). PD is predicted to affect 14 million people worldwide by 2040 (Dextera and Jenner, 2013; Dorsey and Bloem, 2018; Dorsey et al., 2007) and pathology includes the degeneration of DAergic neurons in the nigrostriatal system, leading to dopamine deficiency in the striatum. However, unlike in PD, GCH1-deficient DRD patients do not show neuronal degeneration despite significantly reduced dopamine levels (Fujitani et al., 2013; Rajput et al., 1994). Instead, GCH-1 deficient patients may present with young-onset cases of parkinsonism with movement abnormalities and non-motor psychological symptoms such as depression and anxiety in adult cases (Hahn et al., 2001; Van Hove et al., 2006; Trender-Gerhard et al., 2009), resembling typical PD symptoms (Nygaard et al., 1992; Trender-Gerhard et al., 2009). However, there has been no attempt to investigate causal link between BH4 deficiency and PD pathogenesis. In this study, we investigated the effects of BH4 deficiency or augmentation on the nigrostriatal dopaminergic system using genetic and pharmacological mouse and human-derived models. Our findings suggest that the BH4 metabolic pathway plays a critical role in the health of mitochondria and DAergic neurons, and its disruption can lead to various DRD- and PD-related pathologies, depending on the extent of BH4 deficiency. Furthermore, our results highlight the potential of targeting this pathway as a therapeutic approach for PD.

## RESULTS

### *GCH1* mutations and reduced plasma BH4 levels in PD-affected individuals

*GCH1* mutations have been linked to PD in recent human genetic association studies (Mencacci et al., 2014; Nalls et al., 2019; Rudakou et al., 2019a; Xu et al., 2017; Yoshino et al., 2018). To further investigate the role of the GCH1/BH4 pathway in PD, we recruited non-genotyped, sporadic late-onset PD-affected individuals and healthy age-matched volunteers, and measured plasma BH4 levels (**Table S1**) (Scheffer et al., 2021). We found a significant reduction in plasma BH4 levels in late-onset sporadic PD subjects, along with a decrease in plasma dopamine and serotonin levels (**Figure 1A).** We also compiled a list of published mutations in *GCH1* associated with major movement disorders caused by dopamine deficiency such as DRD and PD (**Table S2**) and mapped these missense mutations onto the three-dimensional structure of human GCH1 (**Figure 1B**). Computational modeling revealed that most of these mutations destabilize GCH1 protein function **(Table S3)**. Although historically most *GCH1* mutations have been associated with DRD, an increasing number of *GCH1* mutations are now being found in PD-affected individuals (**Table S2**). These findings indicate that *GCH1* mutations are frequently associated with both DRD and PD-affected individuals, and that plasma BH4 levels are reduced in patients with late-onset sporadic PD.

**Figure 1.**
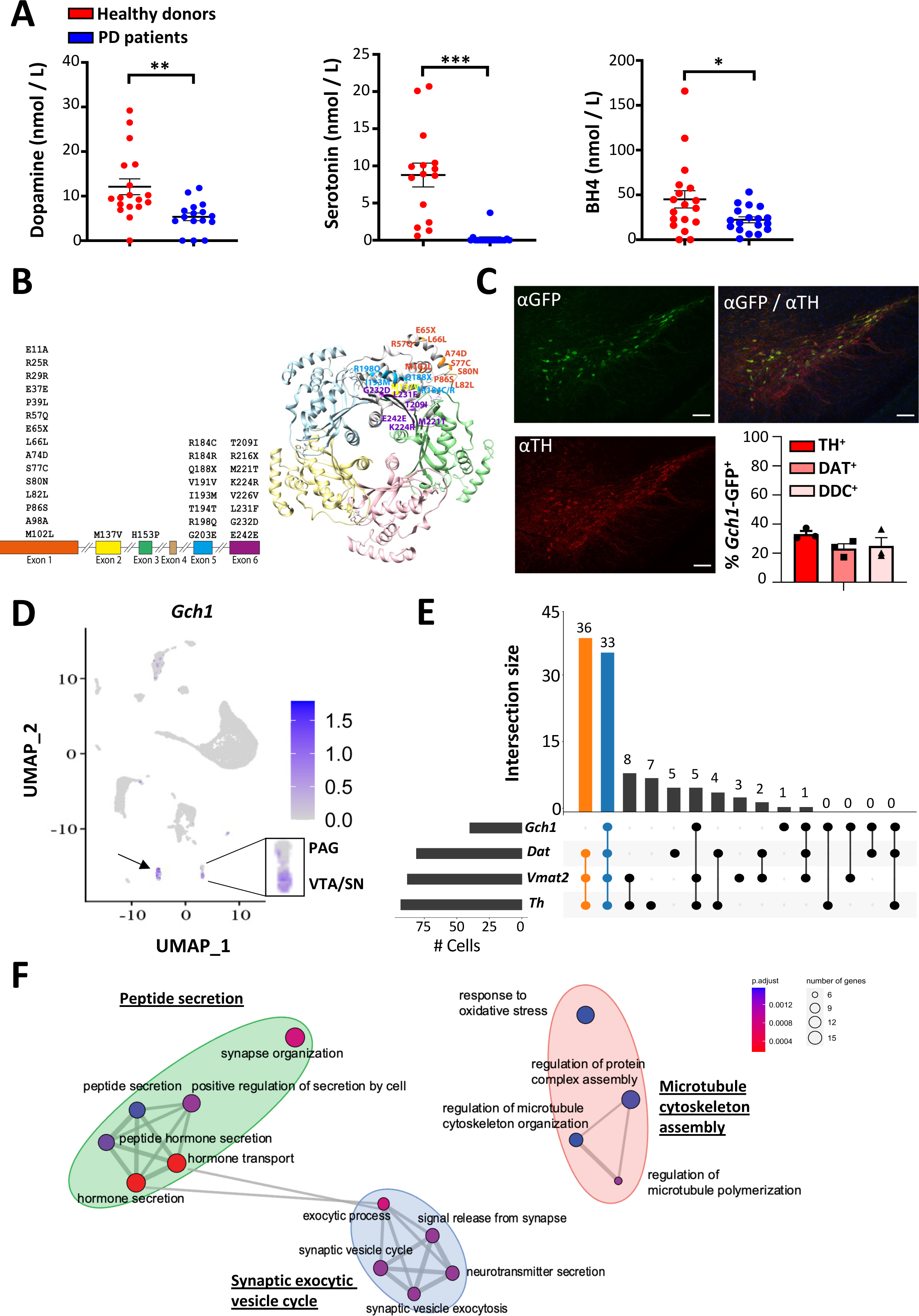
*Gch1*-expressing neurons represent a distinct midbrain DAergic population. **A,** BH4, dopamine, and serotonin levels in the plasma of sporadic PD patients. Data are shown as mean ± s.e.m. Individual PD patients and healthy donors are shown. *P < 0.05; **P < 0.01; ***P < 0.001; (Student’s t-test). **B,** Synonymous and missense *GCH1* mutations associated with Parkinson’s disease which have been reported since 2016 (see **Table S2** for details). Exons are drawn to scale according to their relative lengths (bottom). Three-dimensional configuration of the GCH1 pentamer (top). Mutations are indicated on the grey GCH1 monomer (colors correspond to exons). **C,** Representative immunofluorescence of tyrosine hydroxylase (TH)-positive dopaminergic (DAergic) neurons in the substantia nigra pars compacta (SNpc) area in the midbrain of *Gch1-GFP* reporter mice and percentages of *Gch1*-GFP^+^ neurons of total TH^+^, DAT^+^ and DDC^+^ neurons in the SNpc of reporter mice. Data are shown as mean ± s.e.m. Individual mice for each cell type are shown. Scale bar, 100µm. **D,** Density preserving UMAP (densMAP) embedding plot of ventral midbrain single-cell RNA-seq dataset reflecting expression level of *Gch1*. The rectangular box indicates DAergic neurons of the periaqueductal gray (PAG) (MBDOP1) and VTA/SN (MBDOP2) regions according to the original annotation (Zeisel et al., 2018). Arrow indicates serotoninergic neuronal population. **E,** Intersectional UpSet plot depicting number of cells identified as DAergic neurons (co-expressing *Th*, *Slc6a3* (*Dat*) and *Slc18a2* (*Vmat2*)) from the ventral midbrain, which also do or do not express *Gch1* as blue and orange columns respectively. **F,** Molecular processes enrichment analysis of those DAergic neurons identified in **(E)** which also express *Gch1* compared to those which do not express *Gch1* (see **Table S4** for differential gene list).

### *Gch1*-expressing midbrain cells represent a distinct subset of DAergic neurons

We next investigated the expression pattern of *Gch1* in the mouse brain using mRNA *in situ* data from the Allen Brain Atlas (http://mouse.brain-map.org). *Gch1* mRNA expression was primarily found in the midbrain region of adult mice (**Figure S1B**), colocalized with markers of dopamine-producing (DAergic) neurons, such as tyrosine hydroxylase (*Th*), dopamine transporter (*Dat*), and dopamine/serotonin/norepinephrine vesicle monoamine transporter-2 (*Vmat2*) (**Figure S1B**). We validated these expression patterns using a *Gch1*-GFP reporter line, in which GFP expression is driven by the *Gch1* promoter (Latremoliere et al., 2015). *Gch1*-GFP expression was mainly observed in DAergic neurons of the *substantia nigra pars compacta* (SNpc) and ventral tegmental area (VTA) (**Figure 1C**). Other brain regions where TH is strongly expressed, such as the olfactory bulb, locus coeruleus, nucleus tractus solitarius, and medial forebrain bundle, did not show *Gch1* expression (**Figure S1C**).

Immunofluorescence staining revealed that approximately 30-40 % of TH-positive (^+^), DAergic midbrain neurons expressed *Gch1*, as confirmed by co-staining with DAT and dopamine decarboxylase (DDC) (**Figure 1C; Figure S2A,B**). To further characterize the *Gch1* expression pattern, we analyzed published single-cell RNAseq data of various cell types in the midbrain region of mice (**Figure S3A; Table S4**) (Zeisel et al., 2018), and found that *Gch1* is predominantly expressed in DAergic neurons of the SN/VTA region (**Figure 1D**). Interestingly, *Gch1* is also expressed in midbrain serotonergic neurons (**Figure 1D**). Other components of the BH4 synthetic pathway *(Pts, Spr, Pcbd1/2, Qdpr, Dhfr*) showed less constrained expression compared to *Gch1,* and did not show significant association with the expression of DAergic markers, unlike *Gch1,* which was significantly correlated with DAergic neurons of the SN/VTA (**Figure S3B,C**). Moreover, single-cell RNAseq analysis of the DAergic neurons defined by the expression of *Th*, *Dat* and *Vmat2,* revealed a specific population of *Gch1*-expressing DAergic neurons in the SN/VTA region, which comprises approximately 30 % of the total DAergic neurons (**Figure 1E**). Comparative pathway enrichment analysis revealed that these *Gch1*-expressing DAergic neurons are associated with synaptic vesicle exocytosis and neurotransmitter release as well as oxidative stress responses (**Figure 1F**), which align with converged pathogenic mechanisms of other monogenetic forms of PD. These data indicate that *Gch1*-expressing DAergic neurons represent a specialized and distinct subtype of midbrain DAergic neurons.

### Ablation of *Gch1* in DAergic neurons results in movement and behavioral defects

To investigate the specific role of BH4 in DAergic neurons, we generated DAergic-specific *Gch1* knockout mice by crossing a *Dat-Cre* driver line with mice in which exons 2 and 3 of the *Gch1* gene were flanked by flox sequences (*Gch1^flox/flox^*) (Chuaiphichai et al., 2014; Zhuang et al., 2005). Cre-mediated deletion ablates the active site of the GCH1 enzyme (hereafter referred to as *Gch1^flox/flox^;Dat-Cre*) (**Figure 2A**). Cre-mediated deletion of *Gch1* led to a significant reduction in BH4 and dopamine levels in the brain, as well as dopamine metabolites DOPAC and HVA, whereas serotonin (5-HT) and noradrenaline (NA) levels were unaffected (**Figure 2B,C; Figure S4A**, **S4B**). These mice failed to thrive, lose weight (weighing ∼ 50% less than their littermates by 3 weeks), and died prematurely starting from postnatal day 16 (**Figure S4C-E**).

**Figure 2.**
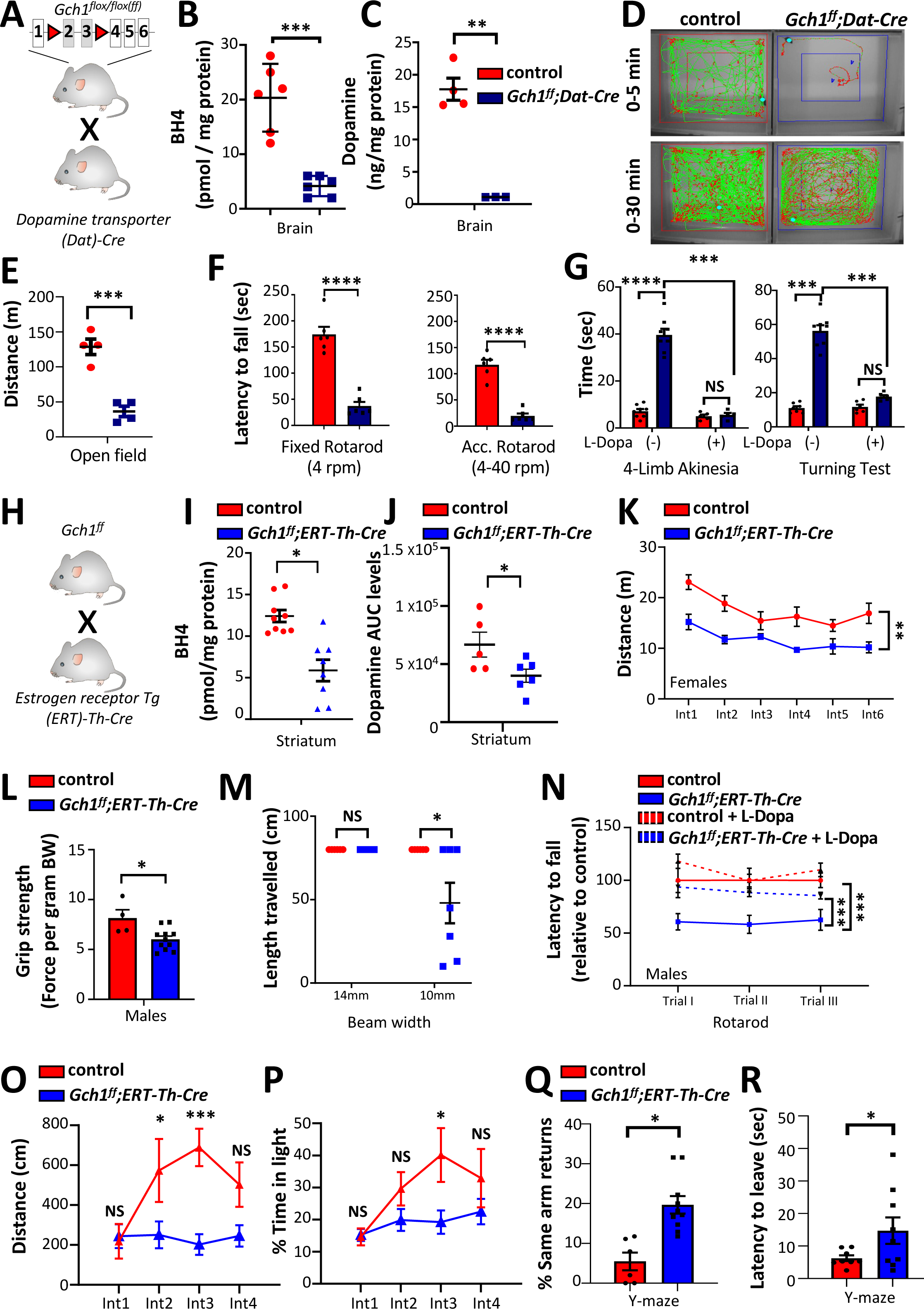
*Gch1* deficiency in DAergic neurons diminishes midbrain dopamine metabolism with accompanying progressive motor function defects and increased anxiety. **A,** Schematic depicting breeding of conditional *Gch1^flox/flox^* to *Dat*-*Cre* line, carrying deletions in exons 2 and 3 (in grey boxes) of *Gch1* in DAergic neurons. **B, C,** BH4 (**B**) and dopamine (**C**) levels in the brains of control and *Gch1^flox/flox^;Dat-Cre* mice. Data are shown as mean ± s.e.m. Individual mice for each genotype are shown. **P < 0.01; ***P < 0.001; ****P < 0.0001; NS, not significant (Student’s t-test; multiple t-test comparison). **D,** Representative video tracking snapshots of 2-week-old control and *Gch1^flox/flox^;Dat-Cre* mice in open field testing. **E,** Quantification of distance travelled by 3-week-old control and *Gch1^flox/flox^;Dat-Cre* mice in open field testing. Data are shown as mean ± s.e.m. Individual mice for each genotype are shown. ***P < 0.001 (Student’s t-test). **F,** Rotarod testing both at fixed speed (4 rotations per minute (rpm)) as well as at accelerated speed (4-40 rpm) on 3-week-old control and *Gch1^flox/flox^;Dat-Cre* mice. Data are shown as mean ± s.e.m. Individual mice for each genotype are shown. ****P < 0.0001 (Student’s t-test). **G,** Four-limb akinesia and turning tests of control and *Gch1^flox/flox^;Dat-Cre* mice before and 45 minutes after i.p. administration of L-Dopa (50mg/kg) and benserazide, (12.5mg/kg). Data are shown as mean ± s.e.m. Individual mice for each genotype are shown. ***P < 0.001; ****P < 0.0001; NS, not significant (2-Way ANOVA with Sidak’s multiple comparisons test). **H,** Schematic showing the breeding of *Gch1^flox/flox^* animals with tamoxifen-inducible *ERT-Th-Cre* line to allow genetic *Gch1* ablation in TH^+^ cells in adult mice. **I,J,** Striatal BH4 **(I)** and dopamine **(J)** levels in control and *Gch1^flox/flox^;ERT-Th-Cre* mice at 28 weeks post-tamoxifen administration. Data are shown as mean ± s.e.m. Individual mice for each genotype are shown. *P < 0.05; **P < 0.01; ***P < 0.001; NS, not significant (Student’s t-test and multiple t-test). **K**, Open-field behavioral testing during 5-minute intervals of a 30-minute observational period in control (n=9) and *Gch1^flox/flox^;ERT-Th-Cre* (n=8) female mice at 28 weeks post-tamoxifen administration for quantifying distance travelled. Data are shown as mean ± s.e.m. **P < 0.01 (2-way ANOVA). **L,** Grip strength of all paws from control and *Gch1^flox/flox^;ERT-Th-Cre* male mice at 28 weeks post-tamoxifen administration. Data are shown as mean ± s.e.m. Individual mice for each genotype are shown. *P < 0.05 (Student’s t-test). **M,** Beam walk behavioral testing of control and *Gch1^flox/flox^;ERT-Th-Cre* male mice at 28 weeks post-tamoxifen administration. Distance traversed across beams of various diameters were recorded. Data are shown as mean ± s.e.m. Individual mice for each genotype are shown. *P < 0.05; NS, not significant (Student’s t-test). **N,** Accelerated (4-40km/hr) rotarod analysis of control and *Gch1^flox/flox^;ERT-Th-Cre* male mice at 28 weeks post-tamoxifen administration, before and after administration of L-Dopa (50mg/kg) plus benserazide, (12.5mg/kg). Data are shown as mean ± s.e.m. Individual mice for each genotype are shown. *P < 0.05, ***P<0.001 (2-way ANOVA with Tukey’s multiple comparison test). **O,P**, Light/Dark box behavioral analysis of control and *Gch1^flox/flox^;ERT-Th-Cre* male mice at 28 weeks post-tamoxifen administration for measuring interval distances **(O)**, and proportion of time spent in the light compartment **(P)** Data are shown as mean ± s.e.m. *P < 0.05, ***P<0.001; NS, not significant (2-way ANOVA with Sidak’s multiple comparison test). **Q,R,** Y-maze behavioral test in which same arm returns **(Q)** and latency to initial movement **(R)** were noted. Data are shown as mean ± s.e.m. Individual mice for each genotype are shown. *P < 0.05 (Student’s t-test).

Behavioral phenotyping of *Gch1^flox/flox^; Dat-Cre* mice showed severe motor and coordination defects. At 2 weeks of age, the mutant mice displayed freezing behavior in the open field test (**Figure 2D; Figure S4F,G**), which is also observed in human PD patients (Amboni et al., 2008; Contreras and Grandas, 2012). At 3 weeks of age, *Gch1^flox/flox^; Dat-Cre* mice exhibited poor movement and coordination on the rotarod (**Figure 2E,F**), which could be improved by administering L-Dopa and benserazide, drugs commonly used in the treatment of PD (**Figure 2G)**. However, the early-onset *Gch1* deficiency in DAergic neurons of these mice was so severe that it limited the study of non-motor defects associated with *Gch1* deficiency.

To investigate the long-term effects of *GCH1* deficiency and model adult-onset parkinsonism in humans (Hahn et al., 2001; Van Hove et al., 2006; Trender-Gerhard et al., 2009), we used tamoxifen-inducible knockout mice of *Gch1* in adult DAergic neurons (*Gch1^flox/flox^;ERT-Th-Cre*) (Badea et al., 2009; Rotolo et al., 2008) (**Figure 2H**). Tamoxifen administration (2mg/kg i.p. daily for 5 consecutive days) in adult (∼ 8 weeks old) *Gch1^flox/flox^; ERT-Th-Cre* mice reduced BH4 levels and dopamine metabolism in the brain and striatum (**Figure 2I,J; Figure S5A)**. These mice did not show any significant difference in survival compared to littermate controls up to one year after tamoxifen treatment (**Figure S5B**), and their body weights did not differ from control littermates for the first 5-6 months after tamoxifen treatment. However, after 6-10 months post-tamoxifen treatment, *Gch1*-deficient mice showed reduced weight gain (**Figure S5C**). Open field analysis revealed a progressive decline in motor functions in these mice compared to controls with substantial reduction in movement, grip strength, coordination, and dexterity in various motor tests **(Figure 2K-M; Figure S5D-K)**. L-Dopa/benserazide treatment mitigated motor and coordination defects **(Figure 2N; Figure S5K)**, and these effects were observed in both male and female mice (**Figure 2N; Figure S5L**). Moreover, these motor defects seemed to stabilize from 6 months and did not worsen further up to 12 months after tamoxifen treatment (**Figure S5L**).

Anxiety often accompanies weight loss and motor deficiency as a common non-motor symptom of DRD and PD, affecting up to 40 % of PD patients (Dissanayaka et al., 2010). In addition to the reduced number of visits to the center in the open field test (**Figure S5F**), tamoxifen-treated *Gch1^flox/flox^;ERT-Th-Cre* mice also displayed a greater aversion to illuminated areas and less exploratory behavior than their littermate controls in the light/dark transition test (**Figure 2O,P**), indicative of increased anxiety in rodents (Bourin and Hascoët, 2003). Mutant mice also exhibited reduced numbers of head dips and a non-significant trend to spend less time in the open arms of the elevated plus-maze (**Figure S6A,B**), also indicative of anxiety-like behaviors in mice (Walf and Frye, 2007). Furthermore, in the Y-maze test, *Gch1^flox/flox^; ERT-Th-Cre* mice displayed a greater percentage of same arm returns (**Figure 2Q; Figure S6C,D**), indicating reduced exploratory behavior or impaired working memory. Finally, mutant mice also required a significantly longer time to initiate and leave the starting point in multiple tests (**Figure 2R; Figure S6E**). Overall, these data show that *Gch1* deficiency in DAergic neurons leads to a progressive locomotor decline which is responsive to L-Dopa therapy, as well as weight loss, freezing behavior, and anxiety.

### DAergic neuron degeneration depends on the extent of BH4 deficiency

The profound motor deficits in *Gch1^flox/flox^; Dat-Cre* mice were accompanied by a dramatic decrease in TH staining in both SNpc and VTA regions, as well as in DAergic neuronal terminals in the striatum (**Figure S7A-E)**. PD is characterized by the degeneration of DAergic neurons, starting from the axonal terminals in the striatum and eventually including the cell bodies in the SNpc. VTA DAergic neurons also degenerate, although with different kinetics (Alberico et al., 2015). On the other hand, midbrain DAergic neurons and their axonal projections to the striatum are intact in DRD patients (Snow et al., 1993). Since this is an important distinguishing factor between these two conditions, we sought to test whether the loss of TH staining corresponded to a loss of DAergic neurons.

Western blot analysis for TH protein showed that *Gch1^flox/flox^; Dat-Cre* mice were born with similar TH levels in the ventral midbrain compared to control mice, but TH levels significantly dropped in *Gch1* mutant animals by three weeks of age (**Figure S7F**). DAT and DDC protein levels were comparable between control and Gch*1^flox/flox^;Dat-Cre* mice at the three-week stage (**Figure S7F**). Additionally, immunofluorescence also depicted a selective loss of TH staining (and not DAT) in DAergic neurons in both SNpc and striatum of *Gch1^flox/flox^;Dat-Cre* mice (**Figure S7G-I**). To validate these findings, we crossed Gch*1^flox/flox^;Dat-Cre* mice onto *Gch1*-GFP reporter mice to easily identify *Gch1*-expressing neurons (**Figure 3A**). Consistent with our earlier findings, we observed a loss of TH staining in *Gch1*-expressing DAergic neurons, demonstrating that loss of *Gch1* in TH^+^ neurons does not affect neuronal viability at this stage (**Figure 3B,C**). As with the early onset model, we detected a dramatic reduction in TH staining intensity but not in the actual number of TH-positive cells in the striatum and the SNpc of *Gch1^flox/flox^; ERT-Th-Cre* mice, 6 months after tamoxifen treatment (**Figure 3D,E**). This data was further validated by Western blot analysis, which showed substantial reduction in TH expression but normal DAT levels (**Figure 3F; Figure S7J-M**), confirming that neuronal degeneration was unaffected by loss of *Gch1* at both 6- and 12-months post-tamoxifen treatment (in both males and females).

**Figure 3.**
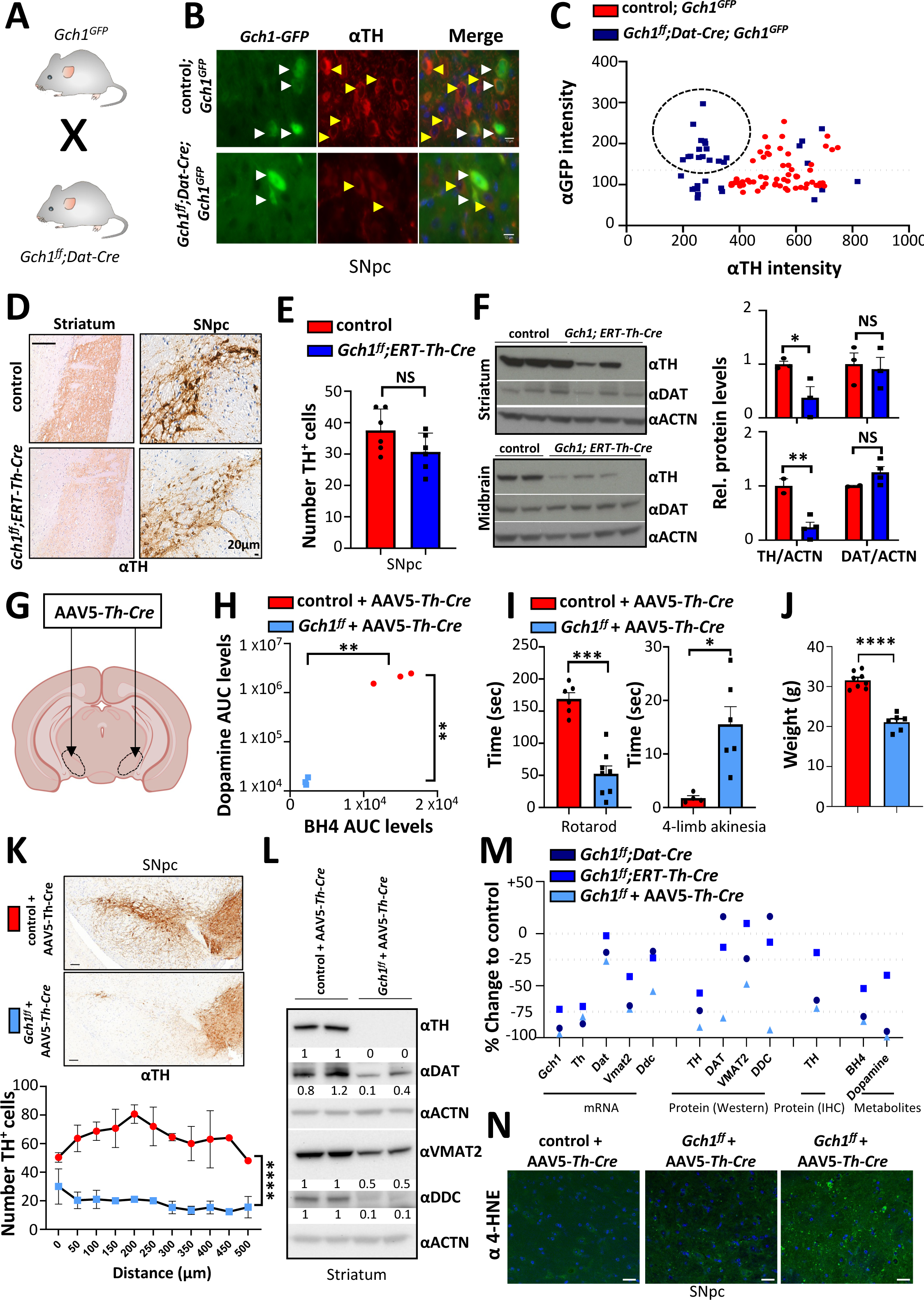
Inducible *Gch1* deletion in dopaminergic neurons results in reduced TH levels. **A**, Schematic showing the crossing of the *Gch1*-GFP reporter line with the *Gch1* deficient line, *Gch1^flox/flox^;Dat-Cre*. **B,C,** Representative immunofluorescence of TH and GFP in the SNpc region **(B)** and quantification of staining intensities **(C)** in 3-week-old control; *Gch1^GFP^* and *Gch1^flox/flox^;Dat-Cre; Gch1^GFP^* mice. Scale bar, 10µm. Data are shown as mean ± s.e.m. Individual cell intensities pooled from n=3 mice for each genotype are shown. Dotted circle indicates *Gch1*-GFP expressing neurons which have low/absent TH expression in *Gch1^flox/flox^;Dat-Cre; Gch1^GFP^*mice. **D,E,** Representative TH^+^ immunohistochemistry in the striatum and SNpc regions **(D)** and quantification of TH-positive cells (**E**) in the SNpc of control and *Gch1^flox/flox^;ERT-Th-Cre* mice at 28 weeks post-tamoxifen administration. Data are shown as mean ± s.e.m. Individual mice for each genotype are shown. NS, not significant (Student’s t-test). **F,** Western blot analysis of TH and DAT in the striatum and ventral midbrain of control and *Gch1^flox/flox^;ERT-Th-Cre* mice at 28 weeks post-tamoxifen administration. N= 2/3 (control); 4/3 (*Gch1^flox/flox^;ERT-Th-Cre*) per genotype. Actin is used as a loading control (left) and normalized TH and DAT protein levels in each tissue is quantified (right). Data are shown as mean ± s.e.m. Individual mice for each genotype are shown. *P < 0.05; **P<0.01; NS, not significant (Multiple t-test). **G,** Schematic depicting the bilateral stereotactic injection of AAV5-*Th-Cre* into the SN of wild type and *Gch1^flox^*^/flox^ mice. **H,** Striatal BH4 and dopamine levels from AAV5-*Th-Cre* injected control and *Gch1^flox/flox^* mice after 4 months. Data are shown as mean ± s.e.m. Individual mice for each genotype are shown. **P < 0.01; (Multiple t-test comparison). **I,** Rotarod (left) and four-limb akinesia (right) testing of control and *Gch1^flox/flox^* mice at 4 months after AAV5-*Th-Cre* injection. Data are shown as mean ± s.e.m. Individual mice for each genotype are shown. *P < 0.05, ***P<0.001 (Student’s t-test). **J**, Body weights of control and *Gch1^flox/flox^* mice at 4-5 months after AAV5-*Th-Cre* injection. Data are shown as mean ± s.e.m. Individual mice for each genotype are shown. ***P<0.001 (Student’s t-test). **K,** Representative TH^+^ immunohistochemistry in the SNpc region (top) and quantification of TH-positive cells (bottom) throughout the SNpc of control and *Gch1^flox/flox^* mice at 4 months after AAV5-*Th-Cre* injection. Data are shown as mean ± s.e.m. ****P<0.0001 (Two-way ANOVA). **L,** Western blot of TH, DAT, VMAT2 and DDC in striatal tissue from control and *Gch1^flox/flox^* mice at 4 months after AAV5-*Th-Cre* injection. N=2 per genotype/condition. Actin is used as a loading control. **M,** Comparison between various genetic BH4 deficiency models (*Gch1^flox/flox^;Dat-Cre*, *Gch1^flox/flox^;ERT-Th-Cre*, and *Gch1^flox/flox^;* AAV5-*Th-Cre*) and their respective controls in terms of DAergic neuronal identity markers at the mRNA, protein and metabolite levels. **N,** Representative 4-HNE/DAPI immunohistochemistry in the SNpc of control and two *Gch1^flox/flox^* mice 3 months after AAV5-*Th-Cre* injection. 4-HNE, 4-Hydroxynonenal. Scale bar, 25μm.

We next delivered AAV5-*Th*-*Cre* directly into the substantia nigra of control (*Gch1^WT/WT^*) and *Gch1^flox/flox^* animals to induce Cre-mediated excision of *Gch1* specifically in TH^+^ neurons in that region (**Figure 3G**). *Gch1^flox/flox^* animals injected with AAV5-*Th-Cre* exhibited substantially less striatal BH4 and dopamine than control animals injected with AAV5-*Th-Cre* (**Figure 3H**). BH4 and dopamine levels were reduced to a greater extent in these animals (BH4: 84.3 +/- 1.3%; Dopamine: 99.2 +/- 0.1%), compared to tamoxifen-treated *Gch1^flox/flox^;ERT-Th-Cre* mice (BH4: 52.7 +/- 29.6%; Dopamine: 40.1 +/- 20.7%) and as a result, these mice displayed stronger motor defects including bradykinesia, occasional paralysis/dragging of the hind-limbs and tremors, as well as reduced weight (**Figure 3I,J**; **Figure S8A**; **Supplementary Video S1-5**). In contrast to tamoxifen-treated *Gch1^flox/flox^; ERT-Th-Cre* mice, *Gch1^flox/flox^* animals treated with AAV5-*Th-Cre* showed a substantial reduction in the number of TH^+^ cells throughout the substantia nigra (**Figure 3K; Figure S8B**). TH staining in *Gch1^flox/flox^* animals treated with AAV5-*Th-Cre* was substantially reduced, with very few TH^+^ neurons irrespective of staining intensity (**Figure 3D** versus **Figure 3K; Figure S8B**). Western blot analysis also showed a strong reduction of TH staining in *Gch1^flox/flox^*animals delivered with AAV5-*Th-Cre* compared to *Gch1^flox/flox^; ERT-Th-Cre* mice (**Figure 3F** vs **Figure 3L**). Additionally, unlike tamoxifen-treated *Gch1^flox/flox^; ERT-Th-Cre* mice, protein levels of DAT, DDC and VMAT2 were all decreased in *Gch1^flox/flox^* animals treated with AAV5-*Th-Cre* as compared to striatal tissue from control mice (**Figure 3L**).

A comparative assessment of the mRNA, protein, and metabolite levels between the various genetically-induced BH4 deficiency models demonstrated that AAV5-*Th-Cre* administration into *Gch1^flox/flox^* animals induced the strongest depletion of BH4 and dopamine and resulted in diminished expression of several DAergic markers at both mRNA and protein levels (**Figure 3M; Figure S8E,F**), again indicative of DAergic neuron loss. Notably, BH4 deficiency specifically led to decreased expression of both *Th* and *Vmat2* transcripts, as well as their respective proteins (**Figure S8CH**). We confirmed neuronal loss in the SNpc of *Gch1^flox/flox^;* AAV5-*Th-Cre* mice by the substantially reduced numbers and sizes of cresyl violet-stained DAergic neurons when compared to AAV5-*Th-Cre-*treated control animals (**Figure S8I**). Moreover, unlike tamoxifen-treated *Gch1^flox/flox^; ERT-Th-Cre* mice, the AAV5-*Th-Cre-*injected *Gch1^flox/flox^* animals displayed a markedly reduced life span (**Figure S8J** versus **Figure S5B**).

GCH1 has been associated with an iron-dependent mode of cell death called ferroptosis (Dixon et al., 2012; Matsushita et al., 2015), which, has been implicated in PD pathogenesis given the high iron load in midbrain DAergic neurons, (Matak et al., 2016; Oakley et al., 2007). A distinctive marker of ferroptosis is an increase in lipid peroxidation, identified by 4-hydroxynonenal (4-HNE) staining. We observed an increase in 4-HNE staining specifically in the SNpc of *Gch1^flox/flox^;* AAV5-*Th-Cre* mice (**Figure 3N**), suggesting increased ferroptosis. The increase in 4-HNE staining was evident four weeks after AAV5 treatment (**Figure S8K**), progressively worsening (**Figure 3N).** Notably, a baseline 4-HNE staining is observed in the SNPc region under control conditions, indicating constitutively low levels of lipid peroxidation likely kept in check in a BH4-dependent manner (**Figure 3N; Figure S8K**). These data demonstrate that BH4 protects DAergic neurons against ferroptosis, confirming additional cofactor-independent roles for BH4 in promoting DAergic neuronal health. Thus, while inactivation of *Gch1* results in reduced dopamine levels and movement disorders in all our models, loss of DAergic neurons depends on the extent of BH4 deficiency; the greater the BH4 and dopamine depletion, the more likely this leads to neurodegeneration, severe motor defects and premature death (**Figure S9A**). Indeed in support of this hypothesis, we purified two mutants of GCH1; one associated with DRD (R198) (Kim et al., 2008) and another which has been reported to lead to DAergic degeneration and familial PD (R184) (Chenbhanich et al., 2017). Mutation of each residue caused a decrease in GCH1 activity (**Figure S9B**); though notably the mutation associated with DRD had a much milder effect on GCH1 activity than the strong decrease of activity observed with the PD-associated mutation (**Figure S9B**).

### *Gch1/*BH4 deficiency sensitizes neurons to mitochondrial PD stressors

In addition to functioning as a cofactor for TH enzymatic activity and regulating *Th* expression and protein levels in midbrain DAergic neurons, BH4 also regulates mitochondrial respiration in activated T cells (Cronin et al., 2018). Since mitochondrial dysfunction is a key pathological driver of PD (Toomey CA et, 2022), we investigated the effects of GCH1/BH4 on mitochondrial function in the human neuroblastoma SH-SY5Y cell line, which is extensively used to study cellular and biochemical aspects of PD *in vitro* (Xicoy et al., 2017). QM385 blocks BH4 production by inhibiting the activity of sepiapterin reductase (SPR), with a consequent accumulation of the metabolic intermediate sepiapterin as a sensitive and specific biomarker for SPR inhibition (**Figure 4A**) (Cronin et al., 2018). As expected, QM385 treatment reduced BH4 levels in SH-SY5Y cells with a concomitant increase in sepiapterin (**Figure 4B; Figure S9C)**. Furthermore, mitochondrial oxidative phosphorylation (OXPHOS) was compromised, as evidenced by a lower ATP/ADP ratio and a compensatory increase in lactate levels upon QM385 treatment (**Figure S9D,E)**. Based on these findings, we hypothesized that GCH1/BH4 deficiency sensitizes neurons to PD stressors.

**Figure 4.**
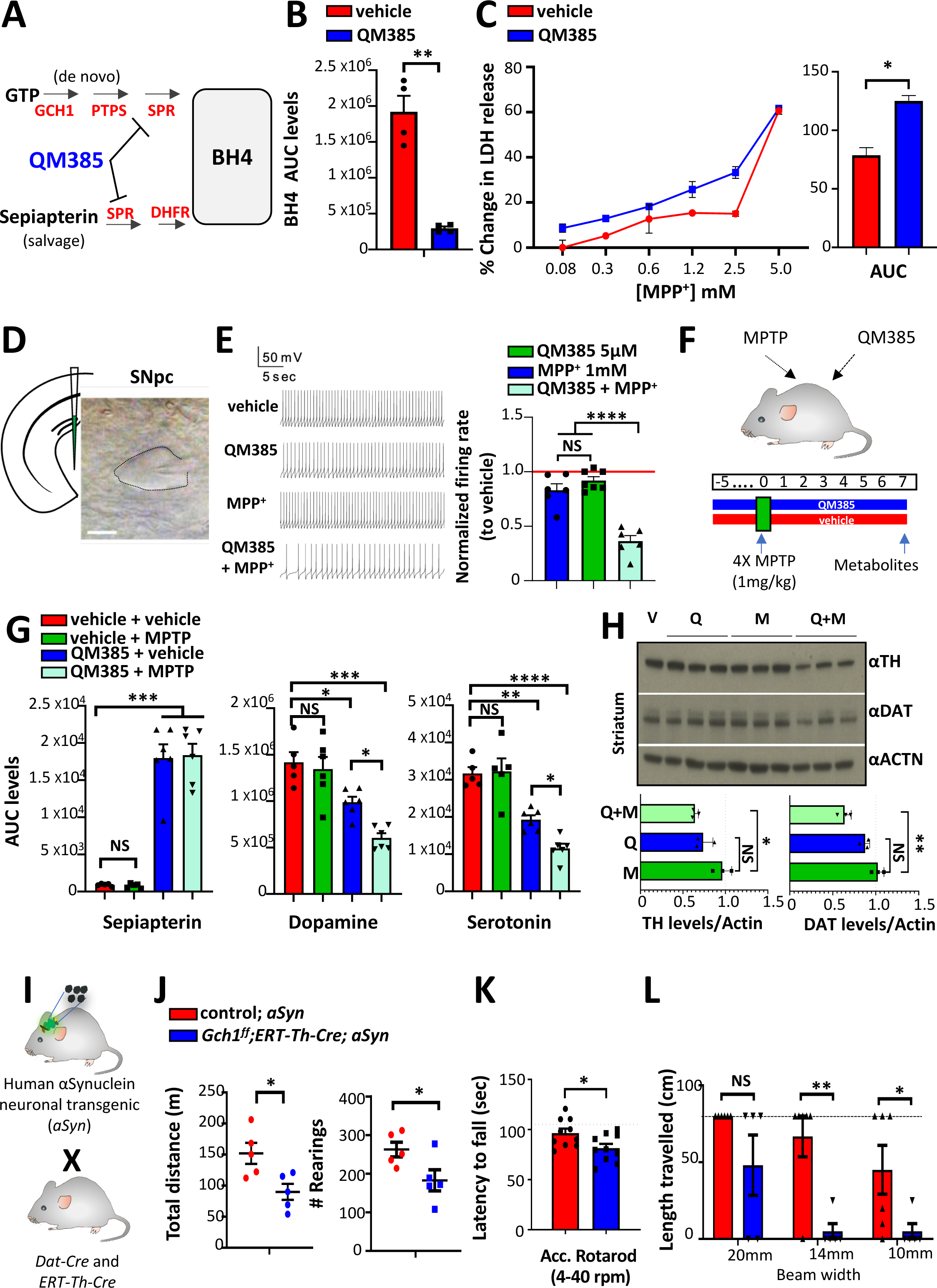
BH4 deficiency sensitizes mice to MPTP and alpha-synuclein PD stressors. **A**, Schematic of BH4 synthesis in the de novo and salvage pathways. Metabolic enzymes are denoted in red. QM385 (in blue) is a specific inhibitor of sepiapterin reductase (SPR). SPR converts the intermediate metabolite sepiapterin (bold) to BH4 in the salvage pathway. PTPS, 6-Pyruvoyltetrahydropterin synthase; DHFR, Dihydrofolate reductase. **B,** BH4 levels in SH-SY5Y cells treated with vehicle (DMSO) or SPR inhibitor (QM385, 5μM) for 16 hours. Data are shown as mean ± s.e.m. Individual samples for each genotype are shown. **P < 0.01 (Student’s t-test). **C,** Percent change (to 0 mM MPP) in lactate dehydrogenase (LDH) release in SH-SY5Y cells obtained by varying doses of MPP^+^ after 24 hours pretreated with vehicle (DMSO) or QM385 (5μM, 5 days) (left) and area under the curve (AUC) analysis. Data are shown as mean ± s.e.m *P < 0.05; **P<0.01; (Student’s t-test). **D**, Schematic of a coronal section of the midbrain containing the SNpc from wild type mouse and an image of a patched recorded neuron. Scale bars, 10 mm. **E,** left, Representative traces of spontaneous pace-making action potentials (APs) from DAergic neurons in mouse SNpc under each condition (before drug treatment (shown as control), 5 μM QM385, 5 μM QM385 with 1 μM MPP^+^); right, Quantification of the frequency of spontaneous pace-making action potentials (APs) under each condition (before treatment (shown as control), 5 μM QM385, 5 μM QM385 with 1 μM MPP^+^,). Data are shown as mean ± s.e.m. Individual brain slice recordings are shown. *P < 0.05; NS, not significant (Paired-sample Wilcoxon signed-rank test). **F,** Schematic depicting sensitization regimen with systemic SPR inhibition (QM385, 5mg/kg i.p.) *in vivo* against low dose MPTP (1mg/kg, 4 times on a single day). **G,** Sepiapterin, dopamine and serotonin levels in the striatum of wild type mice administered with the indicated treatments. Data are shown as mean ± s.e.m. Individual mice for each genotype are shown. AUC, area under the curve (see methods for quantification of metabolites). *P < 0.05; **P<0.01; ***P<0.001; ****P<0.0001; NS, not significant (One-way ANOVA with Tukey’s multiple comparison test). **H,** Western blot (upper) and quantification (lower) of TH and DAT levels in the striatum of wild type mice administered with the indicated treatments. Data are shown as mean ± s.e.m. Individual mice for each genotype are shown. *P < 0.05; NS, not significant (One-way ANOVA with Dunnett’s multiple comparison test). V, vehicle; Q, QM385; M, MPTP. **I,** Schematic depicting the breeding strategy for incorporating the human alpha-synuclein (aSyn) overexpressing transgene into *Gch1^flox/flox^;Dat-Cre* and *Gch1^flox/flox^;ERT-Th-Cre* mice. **J,** Open field behavioral testing of 4-month-old control; *aSyn* and *Gch1^flox/flox^;ERT-Th-Cre; aSyn* mice assessing total distance travelled (left), and number of rearings (right) over a 30-minute observational period. Data are shown as mean ± s.e.m. Individual mice for each genotype are shown. *P < 0.05 (One-way ANOVA analysis with Tukey’s multiple comparison test). **K**,**L,** Accelerated rotarod **(K)** and beam walk behavioral testing **(L)** of 4-month old control; *aSyn* and *Gch1^flox/flox^;ERT-Th-Cre;aSyn* mice. Length travelled along beams of different diameters was measured. Tamoxifen was administered one month before testing to all experimental mice. Dotted horizontal line represents success of control and *Gch1^flox/flox^;ERT-Th-Cre* mice at each beam diameter at this age. Data are shown as mean ± s.e.m. Individual mice for each genotype are shown. *P < 0.05; **P < 0.01; NS, not significant (Student’s t-test and multiple t-test).

Widely-used experimental models of PD employ toxins, especially 1-methyl-4-phenyl-1,2,3,6-tetrahydropyridine (MPTP) and its toxic metabolite MPP^+^ (Bezard et al., 1997; Dawson et al., 2010; Hallman et al., 1985; Meredith and Rademacher, 2011). MPP^+^ enters DAergic neurons through the DAT transporter and specifically blocks complex I of the mitochondrial respiratory chain, causing mitochondrial dysfunction and ultimately cell death (Bezard et al., 1997; Dawson et al., 2010; Hallman et al., 1985; Meredith and Rademacher, 2011). Indeed, blockage of the BH4 pathway with QM385 sensitized SH-SY5Y cells to MPP^+^-mediated toxicity *in vitro* as determined by LDH release (**Figure 4C; Figure S9F**). DAergic neurons in the SNpc exhibit autonomous pace-making action potentials in the absence of stimulatory synaptic input, which maintains extracellular dopamine levels in the striatum (Gantz et al., 2018). Various ion channels (such as ATP-sensitive potassium channels, cAMP-sensitive hyperpolarization-activated cyclic nucleotide-gated channels, and L-type Ca^2+^ channels) regulate this pace-making activity (Grace and Onn, 1989; Guzman et al., 2009; Schultz, 2007; Surmeier et al., 2005). Importantly, mitochondrial dysfunction has been linked to the aberrant firing of DAergic neurons in PD (Branch et al., 2016; Duda et al., 2016; Kann and Kovács, 2007; Liss and Roeper, 2001; Liss et al., 2005; Pacelli et al., 2015), and having established a role for BH4 in mitochondrial bioenergetics in SH-SY5Y cells, we next investigated whether BH4 could affect mitochondria-dependent pace-making activity of DAergic neurons in the SNpc of mice *in vivo* (**Figure 4D**). Indeed, BH4 deficiency induced by QM385 treatment decreased the rate of spontaneous firing in DAergic neurons from midbrain slices of wild-type mice in a dose-dependent manner (**Figure S9G,H**). Importantly, a combination treatment with low doses of QM385 and MPP^+^ synergistically and significantly reduced DAergic firing rates in SNpc slices from wild type mice, compared to treatment with either chemical alone (**Figure 4E**).

In addition to MPP^+^, MPTP can also be administered to mice to specifically target mitochondria of DAT^+^ DAergic neurons in the SNpc (Bezard et al., 1997; Dawson et al., 2010; Hallman et al., 1985; Meredith and Rademacher, 2011). Moreover, *in vivo* administration of MPTP induces the loss of DAergic neurons and dopamine depletion in the striatum and SNpc of mice (Jackson-Lewis and Przedborski, 2007). Therefore, we tested whether systemic BH4 deficiency sensitizes mice to low dose MPTP-mediated dopamine depletion *in vivo* (**Figure 4F**). The treatment of mice with QM385 alone intraperitoneally resulted in reduced dopamine and serotonin levels in the striatum as well as the prefrontal cortex (**Figure 4G; Figure S9I,J**), two brain regions associated with reduced DAergic neuronal input in PD patients that affects their motor functions and mood (Keitz et al., 2008). A dose-response in wild type mice revealed that a low dose of MPTP—1 mg/kg administered 4 times every 2 hours on day zero— did not affect striatal dopamine levels (**Figure S10A**) which is in line with previous reports following the same procedure (Jackson-Lewis and Przedborski, 2007; Meredith and Rademacher, 2011). Furthermore, we observed an increase in the levels of BH2 relative to BH4 (BH2/BH4 ratio) compared to untreated animals, suggesting oxidation of BH4 to BH2 during MPTP treatment (**Figure S10B)**. Importantly, inhibition of BH4 synthesis with QM385 synergized with low-dose MPTP treatment to further reduce both striatal dopamine and serotonin levels (**Figure 4G**), as well as significantly reduced both TH and DAT protein levels, indicating neuronal loss, compared to treatment with either chemical alone (**Figure 4H**). Thus, BH4 deficiency sensitizes DAergic neurons to MPTP toxicity.

### *Gch1*-deficiency sensitizes mice to alpha-synuclein-driven pathologies

Multiple studies indicate that alpha-synuclein (*a*Syn) aggregation plays an important pathological role in individuals with sporadic PD (Gatto et al., 2010; Masliah et al., 2000; Rockenstein et al., 2002; Spillantini et al., 1997). We therefore assessed whether *Gch1* deficiency worsens PD-related behavioral deficits, by crossing *Gch1^flox/flox^Dat-Cre* mice with mice over-expressing human wild-type *a*Syn driven by the murine Thy-1 promoter (*Thy1-aSyn* mice (Chesselet et al., 2012) **(Figure 4I).** *Gch1^flox/flox^; aSyn; Dat-Cre* mice exhibited increased weight loss and more impaired movement at 2 weeks of age compared to controls (*Gch1^flox/flox^; DAT* and *Gch1^WT/WT^; aSyn; Dat-Cre*) mice **(Figure S10D).** Since *Gch1^flox/flox^; aSyn; Dat-Cre* mice were physically too weak to perform any behavioral tests, we generated tamoxifen-inducible mutant mice (*Gch1^flox/flox^; aSyn; ERT-Th-Cre*) **(Figure 4I)**; tamoxifen was administered to 1-month-old mice and behavioral phenotyping was performed 3 months later. Control mice carried the wild-type (non-floxed) alleles of *Gch1* but had both *ERT-Th-Cre* and the *aSyn* transgene and were also administered tamoxifen. As in the case for our rapid-onset model, inducible *Gch1*-deficiency markedly worsened the motor defects of 4-month- old *aSyn* mice, as determined by distance travelled and rearing, in the open field, rotarod and beam walk tests (**Figure 4J-L**). Altogether, these data indicate that *Gch1* deficiency aggravates alpha-Synuclein-driven PD-related pathogenesis in mice.

### Increased BH4 protects DAergic neurons against the mitochondrial PD stressor MPP^+^/MPTP

Based on the above data, we hypothesized that GCH1/BH4 enhancement may protect against PD-related pathology. Indeed, a SNP within *GCH1* (rs841) associated with reduced PD risk shows enhanced GCH1 levels in humans (Rudakou et al., 2019b). Sepiapterin, an intermediatory metabolite of the BH4 pathway can be added exogenously to increase BH4 levels (**Figure 5A**). In accordance with this, while MPP^+^ treatment (similar to QM385 treatment) decreased the rate of spontaneous firing in DAergic neurons from midbrain slices of wild-type mice (**Figure 5B,C;** Chang et al., 2020),) enhancing BH4 production through sepiapterin treatment completely mitigated these altered firing rates (**Figure 5B,C**). Similarly, in current-clamp recordings, DAergic neurons exhibit a pronounced rebound depolarization mediated by the hyperpolarization-activated current (*I*_h_) in response to a series of hyperpolarizing pulses, and such potential reversal changes result in a typical “sag” (Grace and Onn, 1989). MPP^+^ treatment reduced the sag amplitude under each current stimulation; which was significantly ameliorated by increasing BH4 levels with sepiapterin treatment (**Figure S10E,F).** These data show that increased BH4 levels can rescue the inhibitory effects of MPP^+^ on both spontaneous and evoked firing activities of DAergic neurons in mouse SNpc slices *ex vivo*.

**Figure 5.**
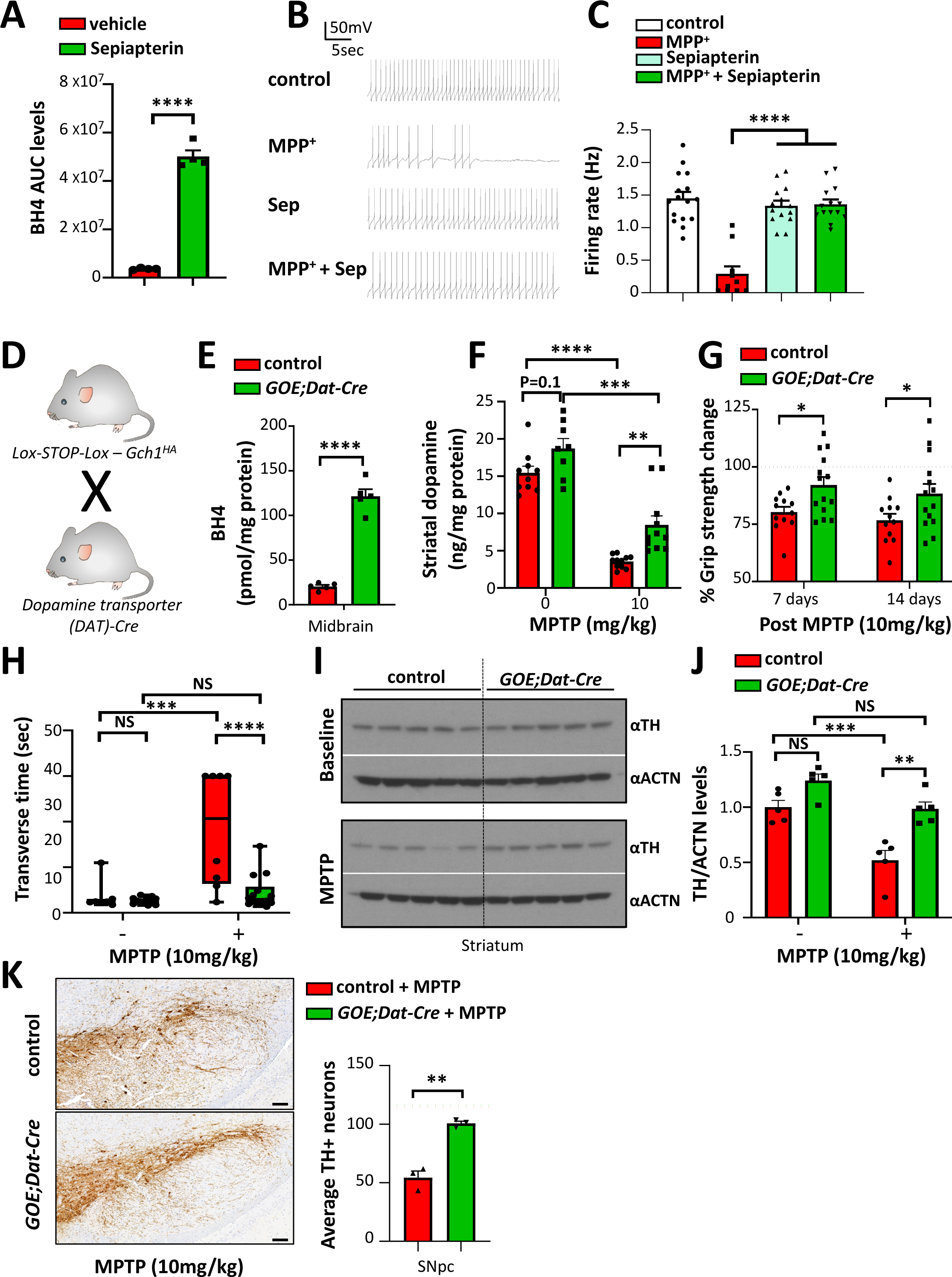
Increased BH4 ameliorates the effects of MPTP treatment as well as human alpha-synuclein overexpression in mice. **A**, BH4 levels in SH-SY5Y cells treated with vehicle (DMSO) or sepiapterin (5μM) for 16 hours. Data are shown as mean ± s.e.m. Individual samples are shown. ****P < 0.0001 (Student’s t-test). AUC, area under the curve (see methods for quantification of metabolites). **B,C,** Representative traces of spontaneous pace-making action potentials (APs) from DAergic neurons in mouse SNpc under each condition (10 mM sepiapterin, 20 mM MPP^+^, 5 mM QM385) (**B**) and quantification of the frequency of spontaneous pace-making action potentials (APs) under each condition (**C**). Data are shown as mean ± s.e.m. Individual brain slice recordings are shown. ****P < 0.0001; NS, not significant (One-way ANOVA with Dunnett’s multiple comparison test). **D,E,** Schematic of breeding to induce *Gch1* transgenic overexpression in DAT^+^ cells (**D**) and midbrain BH4 levels (**E**) in control and *GOE;Dat-Cre* mice. Data are shown as mean ± s.e.m. Individual mice for each genotype are shown. ****P < 0.0001 (Student’s t-test). **F,** Dopamine measurements in the striatum of control and *GOE;Dat-Cre* mice treated with vehicle or MPTP (10mg/kg). Data are shown as mean ± s.e.m. Individual mice for each genotype are shown. *P < 0.05; **P < 0.01; ****P < 0.0001; NS, not significant (One-way ANOVA with Tukey’s multiple comparisons. **G,** Grip strength of all paws from control and *GOE;Dat-Cre* mice, at 7 and 14 days after MPTP treatment. Data are shown as mean ± s.e.m. Individual mice for each genotype are shown. *P<0.05 (Multiple t test comparison). **H,** Beam walk behavioral testing of control and *GOE;Dat-Cre* mice at 14 days after MPTP treatment in which time taken to traverse was measured. **I,J**, Western blot (**I**) and quantification (**J**) of striatal tissue from control and *GOE;Dat-Cre* mice (untreated and at 14 days after MPTP treatment) and blotted with anti-TH. Actin was used as a loading control and for normalization. Data are shown as mean ± s.e.m. Individual mice for each genotype are shown. **P<0.01; ***P<0.001 (Two-way ANOVA with Tukey’s multiple comparison test). **K,** Representative staining of TH and quantification of TH^+^ cells in the SNpc of control and *GOE;Dat-Cre* mice at 14 days after MPTP treatment (10mg/kg). Data are shown as mean ± s.e.m. Individual mice for each genotype are shown. **P<0.01 (Student’s t test). Dashed lines indicate the average number of TH+ cells from naïve control and *GOE;Dat-Cre* mice (see Figure S11G).

To investigate whether our *in vitro* and *ex vivo* findings—on the protective role of enhanced BH4 against MPP^+^ toxicity— translate to an *in vivo* setting, we genetically overexpressed *Gch1* specifically in DAergic neurons using *Dat-Cre* and a conditional *Gch1* over-expressing mouse line (hereafter referred to as *GOE;Dat-Cre*) (Cronin et al., 2018; Latremoliere et al., 2015) (**Figure 5D**). These mice exhibited substantially increased BH4 levels in the ventral midbrain compared to control animals (**Figure 5E**). Behaviorally, both male and female *GOE; Dat-Cre* mice showed no apparent differences in the open field assay, rotarod activity, or grip strength, compared to their littermates (**Figure S11A-F)**. Moreover, *Gch1* overexpression did not affect the number of midbrain DAergic neurons (**Figure S11G).**

We next tested whether *Gch1* overexpression in DAergic neurons *in vivo* protects against MPTP-induced dopamine depletion with two dosing protocols: 1) one injection daily for 5 days; and 2) 4 injections in one day, each separated by a two-hour interval (**Figure S12A-C**). In the first dosing protocol, striatal dopamine levels and its metabolites were reduced in control mice, but largely preserved in *GOE; Dat-Cre* mice (**Figure S13A-G**). Similarly, in the second, stronger dosing protocol, a significantly higher striatal dopamine was observed after MPTP treatment in *GOE; Dat-Cre* mice compared to control animals (**Figure 5F**). Importantly, MPTP treatment in control animals resulted in a significant reduction in grip strength, which was ameliorated in *GOE; Dat-Cre* mice (**Figure 5G**). Additionally, control animals spent less time running and more time stopping after MPTP treatment, which led to longer times to traverse the beam walk; *Gch1* overexpression in DAergic neurons almost completely mitigated these effects (**Figure 5H; Figure S14A-C**). As expected, MPTP administration caused a significant drop in striatal and midbrain TH levels in control animals, which was again prevented by *Gch1* overexpression (**Figure 5I-K**). These data demonstrate that increased BH4 levels in DAergic neurons can ameliorate the biochemical and behavioral defects induced by the mitochondrial PD stressor MPTP *in vivo*.

### GCH1/BH4 protects against DAergic neuron dysfunction in *a*Syn overexpressing mice

In animal models, *a*Syn aggregation, similar to MPTP effects, has been linked to mitochondrial dysfunction and is considered as yet another possible causative mechanism for PD development (Devi et al., 2008; Hsu et al., 2000; Reeve et al., 2015). Therefore, in addition to treating mice with the acute chemical PD stressor MPTP, we also investigated the effect of BH4 enhancement by overexpressing the more chronic PD stressor aSyn. Intriguingly, when *GOE;Dat-Cre* mice were crossed with the *Thy1-aSyn* line, the enhanced BH4 production in DAergic neurons significantly ameliorated the motor defects observed during rotarod, open field and beam-walk tests as compared to motor behaviours in *aSyn;Dat-Cre* control littermates (**Figure S14D-I**). Importantly, the reduction in midbrain dopamine content in 8-month-old *Thy1-aSyn* mice (Chesselet et al., 2012) was normalized by increased BH4 production in the *GOE; aSyn;Dat-Cre* mice (**Figure S14J-L**). These data show that the GCH1/BH4 pathway protects against DAergic neuron dysfunction produced by chronic genetic *a*Syn overexpression.

### BH4 promotes mitochondrial health in DAergic neurons

Numerous studies have demonstrated the importance of axonal mitochondria health of SNpc DAergic neurons in PD pathogenesis, showing that mitochondrial dysfunction represents an early hallmark of PD initiation (González-Rodríguez et al., 2021; Pickrell et al., 2011; Reeve et al., 2018; Toomey CA et, 2022). Reduction in BH4 led to impaired cellular ATP levels (**Figure S9B**). Conversely, sepiapterin treatment to enhance BH4, increased ATP levels in SH-SY5Y cells (**Figure 6A**). These BH4-induced metabolic effects occurred rapidly within several minutes of treatment (**Figure S15A-C**). Next, we monitored mitochondrial energetics in *GOE;Dat-Cre* mice. We focused on striatal tissue as this brain region contains axonal mitochondria in the projections from the SNpc DAergic cell bodies. Notably, GCH1 localized at the soma as well as at the axonal projections from midbrain DAergic neurons into the striatum (**Figure 6B**). As expected from these projections, striatal tissue from *GOE;Dat-Cre* mice displayed significantly increased BH4 and dopamine levels (**Figure 6C**).

**Figure 6.**
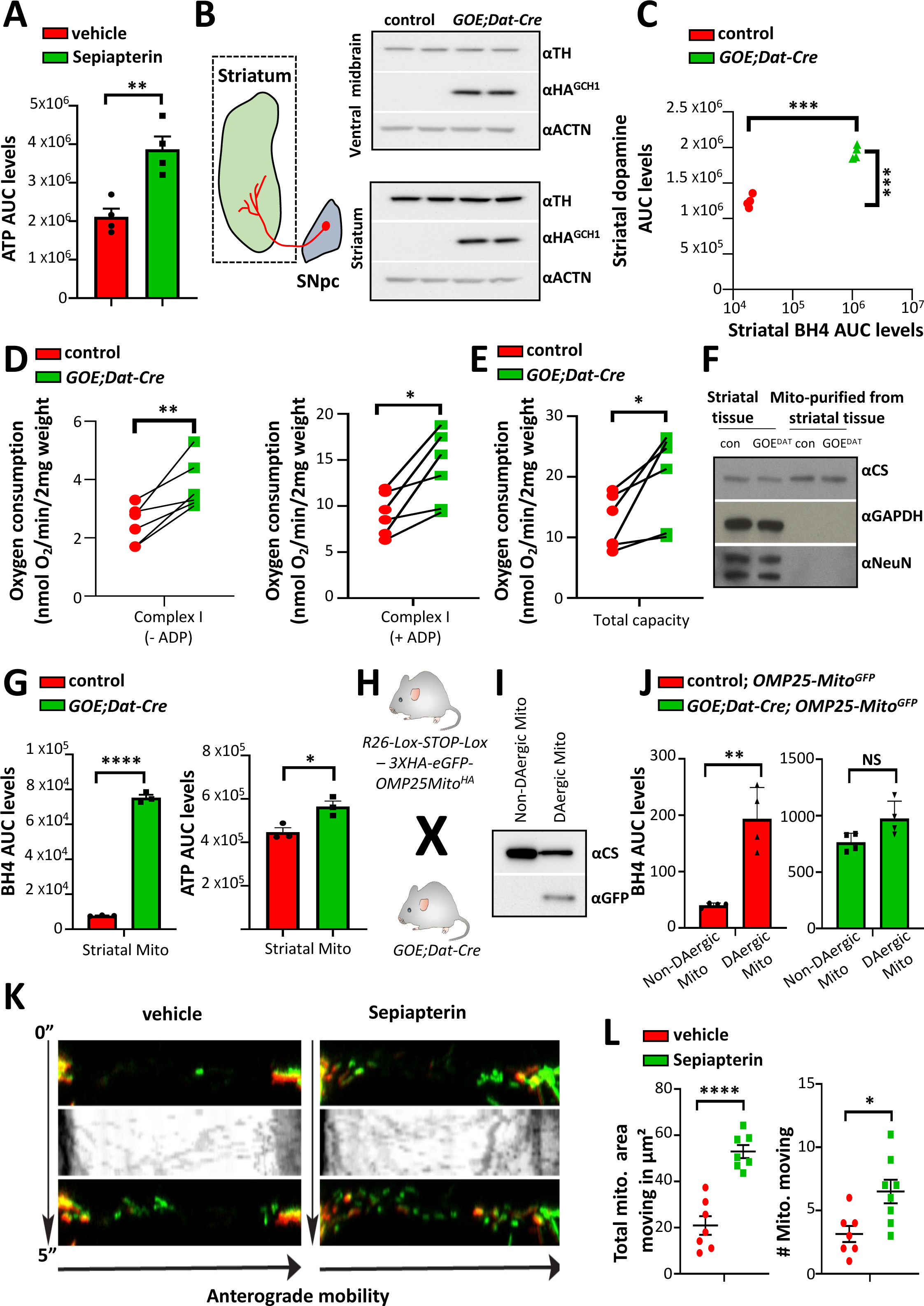
Mitochondrial BH4 regulates mitochondrial respiration, energy output and movement. **A**, ATP levels in SH-SY5Y cells treated with vehicle (DMSO) or sepiapterin (5μM) for 16 hours. Data are shown as means ± s.e.m. Individual samples for each genotype are shown. **P < 0.01 (Student’s t-test). AUC, area under the curve (see methods for quantification of metabolites). **B,** Schematic depicting the projections of DAergic neurons from the SNpc to the striatum (left). Dotted line indicates the extraction of the striatum, including projections from the SNpc DAergic neurons, but not the cell bodies. Western blot of TH and HA^GCH1^ in the ventral midbrain and striatal tissue isolated from control and *GOE;Dat-Cre* mice. Actin is used as a loading control. **C,** Dopamine and BH4 levels in the striatum of control and *GOE;Dat-Cre* mice. Data are shown as mean ± s.e.m. Individual samples for each genotype are shown. ***P < 0.001 (Student’s t-test). AUC, area under the curve (see methods for quantification of metabolites). **D,E,** Complex I mitochondrial activity **(D)** with (right panel) and without (left panel) ADP as well as total capacity of mitochondrial respiration **(E)** in striatal tissue from control and *GOE;Dat-Cre* mice. Data are shown as mean ± s.e.m. Individual mice for each condition are shown. *P < 0.05 (paired Student’s t-test). **F,** Western blot of the striatum as well as isolated mitochondria from the striatal tissue of control and *GOE;Dat-Cre* mice stained with a mitochondrial protein (citrate synthase (CS)), cytoplasmic protein (GAPDH) and a nuclear protein (NeuN). **G,** BH4 (left) and ATP (right) levels in the isolated mitochondria from striatal tissue of control and *GOE;Dat-Cre* mice. Data are shown as mean ± s.e.m. Individual mice for each genotype are shown. *P < 0.05; ****P < 0.0001 (Student’s t-test). AUC, area under the curve (see methods for quantification of metabolites). **H,** Schematic depicting the breeding of *GOE;Dat-Cre* with transgenic reporter mice (*R26-Lox-STOP-Lox – 3XHA-eGFP-OMP25Mito^HA^; referred to as OMP25-Mito^GFP^*) in which mitochondria are tagged with OMP25-GFP in DAergic neurons. **I,** Western blot of isolated mitochondria from the striatal tissue of control; OMP25-Mito^GFP,^ which were then separated into those originating form DAergic neurons in the SNpc versus all other mitochondria form the striatum. Citrate synthase (CS) and GFP were blotted. **J,** BH4 levels in mitochondria isolated from striatal tissue and separated into those originating from SNpc DAergic neurons and all others, from control; *OMP25-Mito^GFP^* (left) and *GOE;Dat-Cre; OMP25-Mito^GFP^* (right) mice. Data are shown as mean ± s.e.m. Individual mice for each genotype are shown. ***P < 0.001 (Student’s t-test). AUC, under the curve (see methods for quantification of metabolites). **K,L,** Time lapse images indicating mEOS2-labelled mitochondrial movement from the cell body along the axon in primary DRG sensory neurons. Kymographs depict mitochondrial movement over 5 minutes after photoconversion so that only newly transported (green-positive) mitochondria were analysed both for movement **(K)**. Quantitation of total mitochondrial area (left) and mitochondrial numbers (right) transported anterograde from soma to axons after 4 days of treatment with vehicle, and sepiapterin (5µM) **(L)**. **P < 0.01; ***P < 0.001; NS, not significant (One-way ANOVA with Dunnett’s multiple comparison test).

To explore whether increased striatal BH4 levels affect mitochondrial respiration, we analyzed oxygen consumption rates of striatal tissue from control and *GOE;Dat-Cre* mice. Enhanced *Gch1* levels, specifically in DAergic neurons, strongly increased mitochondrial complex I activity as well as total mitochondrial respiration (**Figure 6D,E**). Analysis of mitochondria isolated from the striatum of control and *GOE;Dat-Cre* mice showed that BH4 can be detected in purified mitochondria and that increased BH4, caused by *Gch1* overexpression, leads to more ATP production **(Figure 6F,G**). We next selectively separated axonal mitochondria from the DAergic neurons of the SNpc from all other mitochondria isolated from striatal tissue using the *OMP25-Mito^GFP^*reporter mouse line, in which the mitochondria are GFP-labelled specifically in DAT^+^ DAergic neurons (Bayraktar et al., 2019) (**Figure 6H,I; Figure S15D**). Using this approach, we directly confirmed BH4 localization in purified mitochondria and BH4 enrichment in axonal mitochondria originating from SNpc DAergic neurons (**Figure 6J**). Intriguingly, we detected increased BH4 in the non-DAergic mitochondria as well, similar to that in DAergic neurons of *GOE;Dat-Cre* mice (**Figure 6J**), suggesting that BH4 not only acts cell autonomously, but may also benefit mitochondrial health and dopamine synthesis in neighboring non-*Gch1* expressing cells.

To further confirm that increased BH4 levels are linked to mitochondrial fitness and function, we monitored mitochondrial motility as a readout, since ATP produced by mitochondria is critically required for axonal mitochondrial transport in neurons (Miller and Sheetz, 2004; Rintoul et al., 2003). Moreover, impaired mitochondrial motility is associated with reduced neuronal survival in multiple cellular models of familial PD (Godena et al., 2014; Li et al., 2013; Wang et al., 2011). Elevating BH4 in primary cultured neurons using sepiapterin treatment significantly increased mitochondrial movement from the soma to the axons (**Figure 6K,L; Figure S15E**). GCH1 and its metabolite BH4 promote, therefore, mitochondrial health and energetics in DAergic neurons.

### BH4 ameliorates mitochondrial dysfunction in human PD models

As *GCH1* SNPs are associated with the incidence of sporadic PD in humans, we examined whether our findings in mice could be translated to experimental human PD models. Differentiated SH-SY5Y cells express DAT through which MPP^+^ (MPTP-derived metabolite) causes mitochondrial dysfunction and ultimately cell death (Bezard et al., 1997; Dawson et al., 2010; Hallman et al., 1985; Meredith and Rademacher, 2011). MPP^+^ treatment induced the disruption of the mitochondrial network and function, as evidenced by the rounded morphology of the mitochondria and enhanced cell death in SH-SY5Y cells (**Figure 7A,B**). In this regard, enhancing BH4 levels through sepiapterin supplementation prevented mitochondrial fragmentation and rounding, and maintained mitochondrial network structure (**Figure 7A**). Importantly, sepiapterin treatment protected SH-SY5Y cells from MPP^+^-mediated cell death in a dose-dependent manner (**Figure 7B; Figure S16A**) as well as reduced MPP^+^-mediated mitochondrial superoxide reactive oxygen species (ROS) formation (**Figure 7C**). We excluded physical interaction between sepiapterin and MPP^+^ as our nuclear magnetic resonance spectroscopy study showed no chemical shifts in a mixture of the two compounds (**Figure S16B)**. BH4 exerts important dopamine-independent, mitochondria-protective effects against the complex I stressor MPP^+^ in a human neuronal cell line.

**Figure 7.**
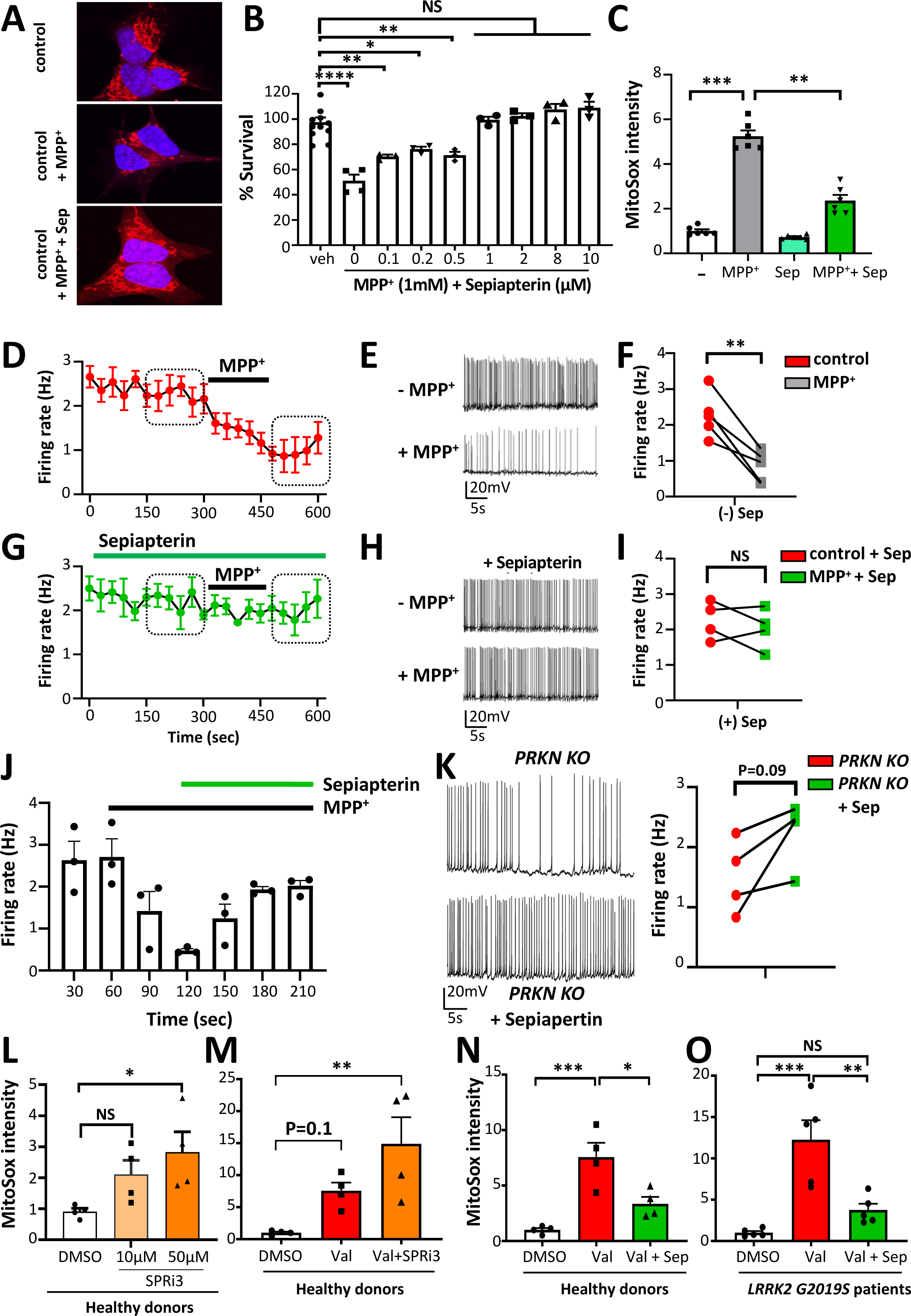
BH4 protects a DAergic human cell line, human midbrain organoids and PD patient-derived fibroblasts from mitochondrial stress. **A**, Representative confocal image of differentiated SH-SY5Y neuroblastoma cells treated with vehicle (control), MPP^+^ (1mM), and MPP^+^ plus sepiapterin (Sep, 2μM) stained with Mitotracker-Red to image mitochondrial morphology and networks after 24 hours of treatment. **B,** Survival quantification of differentiated SH-SY5Y neuroblastoma cells treated for 48 hours with DMSO vehicle (control), MPP^+^ (1mM), and MPP^+^ plus different doses of sepiapterin (as indicated in the figure). Data are shown as mean ± s.e.m. Individual samples for each condition are shown. *P < 0.05; **P < 0.01; ****P < 0.0001; NS, not significant (One-way ANOVA with Dunnett’s multiple comparison test). **C,** Normalised MitoSOX intensity measurements of SH-SY5Y cells untreated or treated for 16 hours with 1mM MPP^+^ with or without sepiapterin (5μM). Data are shown as mean ± s.e.m. Individual samples for each condition are shown. **P < 0.01; ***P < 0.001 (One-way ANOVA with Tukey’s multiple comparison test). **D,** Time course of the effect of MPP^+^ (20 μM) treatment on spontaneous AP firing frequencies in TH-GFP^+^ neurons within hMLOs (horizontal bar indicates the timing of MPP^+^ bath application). **E,** Representatives traces of spontaneous APs from TH-GFP^+^ neurons before (top panel) and after 3 minutes of MPP^+^ application (20 μM, bottom). **F,** Quantification of the effect of MPP^+^ treatment on the frequencies of spontaneous APs recorded from TH-GFP^+^ neurons. Data are shown as mean ± s.e.m. Individual recordings from hMLOs are shown. **P < 0.01 (paired Student’s t-test). **G,** Time course of the effect of MPP^+^ treatment in the presence of 10 mM sepiapterin on spontaneous AP firing frequencies recorded from TH-GFP^+^ neurons (horizontal bar indicates the timings of sepiapterin or MPP^+^ application). **H,** Representative traces of spontaneous APs from TH-GFP^+^ neurons before (top panel) and 3 minutes after bath application of MPP^+^ in the presence of sepiapterin (10 mM). **I,** Quantification of the effect of MPP^+^ on TH-GFP^+^ neurons in the presence of sepiapterin. Data are shown as mean ± s.e.m. Individual recordings from hMLOs are shown. NS, not significant (paired Student’s t-test). **J**, Firing rates of TH^+^ neurons in hMLOs treated with MPP^+^ and subsequently treated with sepiapterin as shown. Applications of MPP^+^ (black line) and sepiapterin (green line) are indicated. **K,** Representative traces of spontaneous APs recorded from DAergic neurons within hMLOs derived from isogenic *Parkin2* knock-out (*PARK2* KO) hESCs before (top) and after (bottom) treatment with sepiapterin (10 mM) and quantification of the effect of sepiapterin on DAergic neurons in *PARK2* KO organoids. Data are shown as mean ± s.e.m. Individual recordings from hMLOs are shown. P-value is shown (paired Student’s t-test). Sep, sepiapterin. **L-O**, Regulation of mitochondrial superoxide levels by BH4 in healthy human fibroblasts as well as in PD patient fibroblasts carrying the *LRRK2 p.G2019S* mutation. MitoSOX fluorescent levels were measured after 24 hours in healthy human fibroblasts treated with the SPR inhibitor SPRi3 **(L)**. MitoSOX levels were monitored in 24-hour valinomycin (10μM)-treated fibroblasts from healthy donors which were pre-treated with vehicle or SPRi3 (50μM) for 24 hours before valinomycin application **(M)**. MitoSOX levels were monitored in 24-hour valinomycin (10μM)-treated healthy donor **(N)** and *LRRK2 p.G2019S* PD patient **(O)** fibroblasts, pre-treated with vehicle or sepiapterin (25μM) for 24 hours before valinomycin application. Data are shown as mean± s.e.m. Data are normalized to DMSO-treated controls. AUC, under the curve (see methods for quantification of metabolites); Sep, sepiapterin. Individual patients for each genotype are shown. *P < 0.05; **P < 0.01; ***P < 0.001; NS, not significant (One-way ANOVA with Tukey’s multiple comparisons). Sep, sepiapterin.

Human midbrain-like organoids (hMLOs) recapitulate 3-dimensional physiological features of the *in vivo* midbrain and TH^+^ neurons in hMLOs express functional dopamine receptors and exhibit biochemical, transcriptional, and electrophysiological properties of mature DAergic neurons (Jo et al., 2016). Using hMLOs derived from H9 human embryonic stem cells (hESCs) with a knock-in *TH-GFP* reporter transgene, we were able to record pace-making firing specifically from TH-positive DAergic neurons (**Figure S17A-C**). Similar to DAergic neurons in the SNpc mouse brain slices, a QM385-mediated deficiency in BH4, as well as mitochondrial disruption by MPP^+^ treatment, reduced both the pace-making frequencies and ‘sag’ currents in DAergic hMLO neurons in a dose-dependent manner (**Figure 7D-F; Figure S17D-G**). Both pre-treatment and post-treatment with sepiapterin significantly mitigated the effects of MPP^+^ on the pace-making of DAergic neurons in hMLOs (**Figure 7G-J**; **Figure S17H,I**). These data from human midbrain organoids demonstrate that an enhancement of BH4 ameliorates MPP^+^-induced aberrant and diminished firing of DAergic neurons, which is an early hallmark of PD (Branch et al., 2016; Michel et al., 2016).

We next sought to explore if BH4 also protects against genetic PD stressors. Whereas the vast majority of human PD cases are sporadic (Savitt et al., 2006), approximately 5 % are inherited (Lesage and Brice, 2009). A loss-of-function mutation in *Parkin* (*PRKN*, also known as *PARK2*) causes early-onset human PD. *PRKN* plays an important role in mitochondrial quality control, mobility, function and turn-over (Dodson and Guo, 2007; Palacino et al., 2004; Pickrell and Youle, 2015). Importantly, DAergic neurons in hMLOs derived from homozygous isogenic *PRKN* mutated hESCs exhibited aberrant pace-making firing activity with reduced frequencies and increased irregularity, which was ameliorated by sepiapterin treatment (**Figure 7K)**. Thus, in addition to its protective effects in human model systems stressed with MPP^+^, increased BH4 can revert impaired pace making activity of *PRKN-* mutant hMLO neurons.

A second genetic mutation associated with PD is in Leucine-rich repeat kinase 2 (*LRRK2*). *LRRK2* is an extensively studied familial PD gene, and fibroblasts from familial, as well as sporadic, *LRRK2-*mutant PD patients are more sensitive to mitochondrial stressors than healthy donor-derived cells (Smith et al., 2016). We therefore quantified mitochondrial superoxide ROS levels to assess mitochondrial dysfunction in fibroblasts from four healthy donors and five *LRRK2 p.G2019S* mutant PD patients. In healthy donor fibroblasts, production of a BH4 deficiency caused by SPR inhibition with SPRi3 (Latremoliere et al., 2015) increased mitochondrial superoxide production (**Figure 7L**). When the mitochondrial membrane depolarizer, valinomycin, was applied to healthy donor fibroblasts, BH4 pathway inhibition augmented this mitochondrial dysfunction, and further increased superoxide ROS levels (**Figure 7M**), again highlighting the synergistic effects of BH4 inhibition with additional PD-related stress. BH4 enhancement using sepiapterin significantly reduced the amount of mitochondrial superoxide in healthy donor fibroblasts stressed with valinomycin (**Figure 7N**). Similarly, enhancing BH4 levels with sepiapterin significantly reduced the elevated level of mitochondrial superoxide in cells from *LRRK2 p.G2019S* PD patients (**Figure 7O)**. Modulation of the BH4 pathway regulates mitochondrial superoxide levels in fibroblasts from healthy donors and enhancing BH4 ameliorates the mitochondrial dysfunctions in *LRRK2 p.G2019S* PD patient fibroblasts to increase energy levels and reduce oxidative stress.

## Discussion

Several studies have linked variants at the *GCH1* locus with movement disorders, such as DRD and PD, whose pathologies affect the nigrostriatal dopamine system (Bandres-Ciga et al., 2020; Mencacci et al., 2014; De Vasconcellos et al., 2015; Wijemanne and Jankovic, 2015). We used *in silico* analysis to identify mutations within the *GCH1* locus that are linked to compromised GCH1 protein stability or impaired activity. Therefore, we created mouse models of *Gch1/*BH4 deficiency through genetic and pharmacological methods to assess their relevance to human pathologies.

Previous mouse models of BH4 deficiency, such as the *hph1* (hypomorphic *Gch1*) mouse, demonstrates reduced BH4 and catecholamine production (modest ∼12% reduction in striatal dopamine) but did not exhibit any movement disorders (Hyland et al., 2003; Nasser et al., 2013). Another mouse model, the *Spr*-deficient mouse (Yang et al., 2006) showed locomotor defects but also had severe reduction in catecholamine neurotransmitters in various brain regions and non-brain areas, making it difficult to assess BH4 effect on specific brain regions or cell types (Yang et al., 2006). To address these limitations, we used the *Dat-Cre* line to selectively remove *Gch1* in DAergic neurons during development and generate tamoxifen-inducible *Gch1*-deficient mice to bypass the requirement for dopamine signaling during embryonic and postnatal development (Zhou and Palmiter, 1995).

The tamoxifen-inducible *Gch1*-deficient mice displayed progressive worsening of motor functions and coordination skills, reduced weight gain, but also behaviors indicative of depression and/or anxiety, resembling non-motor symptoms which are major factors impacting overall health in PD and DRD patients (Chaudhuri et al., 2006; Fujishiro et al., 2008; Katzenschlager and Lees, 2004). These mice also displayed a delayed latency to move, reminiscent of the ‘freezing’ or slow hesitant movements (bradykinesia) behavior seen in DRD and PD patients. The *ERT-Th-Cre* line relies on tamoxifen administration for deletion, which is not as efficient in the brain (Badea et al., 2009; Jahn et al., 2018). We observed a ∼40-50 % reduction of BH4 in inducible adult-onset *Gch1^flox/flox^;ERT-Th-Cre* mice, which produces behavioral and biochemical effects, but wanted to increase the deficiency of BH4 further. Stereotactic injections of AAV5-*Th*-Cre directly into the SN of control and *Gch1^flox/flox^* mice resulted in greater reductions in BH4 and dopamine levels, accompanied by even stronger movement defects, suggesting that the extent of BH4 deficiency may determine the severity of disease associated with *GCH1* mutations. There are several reports which suggest that BH4 may actually be toxic to DAergic neurons (Choi et al., 2006; Enzinger et al., 2002; Lee et al., 2007). Our genetic in vivo data herein show conclusively that BH4 deficiency, and not overexpression, in DAergic compromises neuronal health.

Both DRD and PD are caused by dysfunction of nigrostriatal DAergic neurons, as indicated by the dramatic therapeutic effects of L-Dopa. However, a distinguishing feature between these two conditions is that these neurons degenerate over time in PD while they remain intact in DRD (Tadic et al., 2012). Importantly, our analyses indicate that BH4 is necessary for the health of DAergic neurons and that the extent to which BH4 is diminished may determine whether neurodegeneration will occur or not. Through the analysis of single-cell RNA sequencing data, combined with our *Gch1*-GFP reporter line, we show that *Gch1* is expressed in only a subset (∼30 %) of midbrain DAergic neurons. These *Gch1*-expressing DAergic neurons seem to be enriched for molecular processes associated with dopamine release and oxidative stress responses and may provide a local supply of BH4 to the area for carrying out its numerous cofactor and non-cofactor functions. Indeed, in tamoxifen-treated *Gch1^flox/flox^;ERT-Th-Cre* mice, there was enough BH4 to maintain cell viability even up to 12 months, although its levels were reduced by 50%; however, when BH4 was reduced by >90 % after AAV5-*Th-Cre* treatment, we observed reduced mRNA and proteins levels of several DAergic markers, increased lipid peroxidation levels (a distinctive marker for ferroptosis), loss of DAergic neurons in the SNpc, most likely via ferroptosis, and ultimately reduced survival of the animals. GCH1 acts as a 10-subunit decamer (Nar et al., 1995); thus one can imagine how various SNPs or mutations within the GCH1 locus could produce a spectrum of activity defects in the structure of the decamer alone. We purified wild type GCH1 and two mutants, one associated with DRD and another associated with PD; while both mutants showed diminished GCH1 activity compared to wild type GCH1, the mutant associated with PD displayed a greater defect to the DRD-associated mutation. Altogether, these data demonstrate how various GCH1 mutations could lead to DRD and PD, with the extent of BH4 pathway disruption induced by the mutations determining the presence or absence of neuronal degeneration.

BH4 is known to be a cofactor necessary for the activity of aromatic amino acid hydroxylases such as tyrosine hydroxylase (TH) during the synthesis of dopamine. Our study now highlights cofactor-independent functions of BH4 in DAergic neurons, such as its protective role against ferroptosis. Ferroptosis is an iron-dependent cell death pathway characterized by increased lipid peroxidation (Dixon et al., 2012). DAergic neurons are particularly sensitive to ferroptosis given the relative importance of iron metabolism in these neurons (Matak et al., 2016; Oakley et al., 2007). The GCH1/BH4 pathway is protective against ferroptosis (Kraft et al., 2020) and we now show that with strong BH4 deficiency, lipid peroxidation and ferroptosis is induced, which may be responsible for the observed neurodegeneration. Furthermore, we show that BH4 mediates other non-cofactor/TH-related functions in DAergic neurons such as maintaining energy and ROS levels in the mitochondria. Mitochondrial health is associated with the firing rates of DAergic neurons, enabling them to release dopamine (Liss et al., 2005; Masi et al., 2015; Yee et al., 2014). BH4 reduction in mouse brain slices and human midbrain-like organoids (MLOs) markedly impaired the spontaneous firing of DAergic neurons (pacemaker effects). Further analysis is required to discern whether these pacemaker effects are dependent on additional, non-mitochondrial BH4 functions. BH4 can also act as a superoxide scavenger (Bailey et al., 2017; Cronin et al., 2018; Nakamura et al., 2001a), protecting DAergic neurons from the damaging effects of ROS (Nakamura et al., 2000, 2001b), and fibroblasts from *GCH1* mutant individuals have increased superoxide levels (Terbeek et al., 2021). Increased BH4 levels help maintain mitochondrial structure and lower mitochondrial superoxide ROS levels in SH-SY5Y cells challenged with the mitochondrial complex I toxin MPP^+^. Additionally, we extended our findings to include primary fibroblasts (with stressed mitochondria) from healthy donors and *LRRK2 p.G2019S* patients, confirming that increased BH4 effectively reduced detrimental mitochondrial ROS levels in cells from human PD patients.

BH4 levels in the cerebrospinal fluid correlated inversely with the progression of neurodegeneration in PD patients (Ichinose et al., 2018). Given human genetic association studies linking *GCH1* mutations and PD, as well as the link between BH4 levels and mitochondrial function, we hypothesized that BH4 deficiency may render DAergic neurons more susceptible to PD stressors. Indeed, pre-treatment of SH-SY5Y cells with QM385 to inhibit SPR and reduce BH4, made cells more susceptible to the mitochondrial destabilizer and neurotoxin MPP^+^. These data were confirmed *in vivo* with the systemic administration of QM385, which increased the efficacy of low dose MPTP in reducing striatal dopamine levels as well as TH and DAT protein levels that are indicative of neuronal loss. In addition to this chemical-induced PD model, we have also demonstrated that *Gch1* deficiency increases the susceptibility of mice to *a*Syn overexpression. Previous studies in fruit flies have shown that knockdown of the *Gch1* orthologue increased motor defects in *a*Syn transgenic flies (Sarkar et al., 2020). Conversely, mice overexpressing *Gch1* specifically in DAergic neurons, rescues disease manifestation caused by MPTP as well as human *a*Syn overexpression. These findings in mice could be replicated in fibroblasts derived from PD patients and, importantly, human MLOs. Overall, in all these different experimental models, higher BH4 levels exerted neuroprotective effects. Our results further indicate that induction of the BH4 pathway not only prevents disease initiation but also restores cellular function after disease progression, making the pathway a potential therapeutic target for both early and late stages of PD. Moreover, we show with systemic QM385 treatment that serotonin levels are also reduced upon BH4 deficiency. As BH4 is also required for serotonin production (via tryptophan hydroxylase), it is possible that BH4 treatment would ameliorate PD-associated non-motor symptoms such as depression.

In the 80s, given the prominent role of BH4 in dopamine synthesis, PD patients were actually administered BH4. The results have been conflicting ranging from no effect to improvement of hypokinesia, rigidity, and tremors (Curtius et al., 1984; LeWitt et al., 1986). BH4 being unstable, expensive, and scarce at the time, large scale trials were not pursued. More recently, a lentiviral gene therapy which delivers TH, DDC and GCH1 to non-dopaminergic striatal neurons was evaluated in clinical studies (Jarraya et al., 2009; Palfi et al., 2014; S et al., 2018). This gene therapy was well tolerated, yet only moderate motor improvements were observed in a small phase I/II trial. Our data now proposes a conceptual framework whereby the functions of BH4 can be broadly separated into two fundamental roles in DAergic neurons: cofactor-dependent (TH stability and TH activity) for dopamine synthesis and, arguably more important, cofactor-independent (ROS scavenging and protection against ferroptosis) for maintaining mitochondrial function/health. Under BH4 deficiency, independent roles are more important and would be prioritized to ensure survival of the DAergic neurons and so BH4 is diverted to the mitochondria resulting in reduced TH levels. If the deficiency is so strong that levels drop below a certain threshold where both functions of BH4 are compromised, the DAergic neuron will degenerate. Simply therefore co-delivering GCH1 and TH may increase dopamine levels; but may not be the best strategy to maintain overall DAergic neuronal health (cofactor-dependent plus cofactor-independent roles of BH4) as the BH4 will be used for the excessive TH quantities. Our data suggests the best strategy may be to target the DAergic neurons in the SNpc with strategies to increase BH4 alone; so that in those targeted DAergic neurons, BH4 is boosted to increase all beneficial roles of the metabolite thus simultaneously increasing dopamine synthesis and providing essential protection in the neurons. Interestingly a phase I clinical trial administering a single oral dose of sepiapterin to healthy volunteers increased plasma BH4 levels (Smith et al., 2019). Will be interesting to know whether such treatment gets assess to the brain to increase BH4 levels as a possible future treatment for DAergic dysfunctions.

In summary, our data underscore the evolutionary conservation and multi-faceted roles for the GCH1/BH4 metabolic pathway in maintaining the health of DAergic neurons. These roles include facilitating dopamine synthesis, regulating TH levels, protecting neurons from ferroptosis, scavenging ROS, supporting mitochondrial function, and maintaining neuronal excitability. Consequently, we argue that disruptions of this pathway can lead to diverse neurological conditions, such as DRD and PD, whereas augmentation of neuronal BH4 can protect against PD.

## Acknowledgements

We thank the patients and volunteers who participated in this study, including the NETI Group from Universidade Federal de Santa Catarina. We are also thankful to Helena Dresch Vascouto and Prof. Dr. Maria Fernanda B. A. da Costa for their help in collecting blood samples and applying the clinical scales. We are grateful to Dr. Dorota Szumska for embryo magnetic resonance imaging. We acknowledge the Vienna BioCenter Core Facilities (VBCF) for their phenotyping, metabolomics, and histopathology work. We also thank our many colleagues for their critical input.

## Author contributions

SJFC, together with KC, HSJ and JMP, conceived the project. SJFC performed and designed the experiments with much help from coauthors as follows: WY and HSJ with electrophysiology of midbrain mouse slices and hMLOs; AH, MJC,KC with *Gch1* flox mice and analysis with *Th*-Cre; SLM, DJM with mitochondrial trafficking experiments; AW, AVK with mitochondrial respiration; JAK, OI, PJH with human LRRK2 patient fibroblast experiments; SB with in silico GCH1 mutant modeling and purifications; EOT, TH with single cell RNA sequencing analysis; MSC, MS, RK with NMR analysis; MO, MA, CGD, JDF,DC,VN with qPCR and Western blotting; BLT, DLS with metabolic measurements; LT with GCH1 activity assay; AW,HR, SR, AK,CP with histological work; NA, AL, MC, with GOE mouse genetics and BH4; FCF, RW with human patient cohort sampling.

## Funding

JMP is supported by the Austrian Federal Ministry of Education, Science and Research, the Austrian Academy of Sciences and the City of Vienna and grants from the Austrian Science Fund (FWF) Wittgenstein award (Z 271-B19), the T. von Zastrow foundation, a Canada 150 Research Chairs Program (F18-01336), and CIHR grant (168899). HSJ is supported by the Singapore National Medical Research Council Open-Fund Individual Research Grant (NMRC-OFIRG21jun-0037), the Singapore Ministry of Education Academic Research Fund (MOE-T2EP30121-0032), National Research Foundation Competitive Research Program (NRF-CRP17-2017-03), and Duke-NUS Signature Research Program Block Grant. KMC is supported by the British Heart Foundation (RG/17/10/32859 and CH/16/1/32013). CJW is supported by NIH R35NS105076. RW and AL are supported by the Brazilian National Council for Science and Technology (fellows of CNPq) and the Foundation for Research and Innovation of Santa Catarina State (FAPESC). ALa (JHU) is funded by R01NS112266. TH is supported by the Swedish Research Council (2018-02838); Novo Nordisk Foundation (NNF20OC0063667); Hjärnfonden (FO20190277) and the European Research Council (FOODFORLIFE, 2020-AdG-101021016) and intramural funds of the Medical University of Vienna. EOT is supported by a scholarship from the Austrian Science Fund (FWF, DOC 33-B27). Metabolomics was performed by the VBCF Metabolomics Facility which is funded by the City of Vienna through the Vienna Business Agency. Behavioural assays were performed by the Preclinical Phenotyping Facility at VBCF, a member of the Vienna BioCenter (VBC), Austria and funded by the Austrian Federal Ministry of Education, Science & Research, and the City of Vienna.

## Declaration of interest

The authors declare no conflict of interest.

## Materials and Methods

### Mice

Mice expressing EGFP under the *Gch1* promoter were used to label TH-expressing DAergic neurons (Latremoliere et al., 2015). Mice with a *Gch1-*floxed mice have been previously reported (Chuaiphichai et al., 2014; Latremoliere et al., 2015). For loss-of-function experiments, we bred *Gch1-*floxed mice with DAergic neuron-specific lines including *Th-Cre*, *Dat-Cre,* or tamoxifen-inducible *ERT-Th-Cre* animals (Badea et al., 2009). Human alpha-synuclein overexpression (aSyn) mice (Line ‘61’) have been previously described (Chesselet et al., 2012). Mice with a *Cre*-dependent *GCH1*-HA overexpression cassette to induce BH4 overproduction have been previously reported (Latremoliere et al., 2015). All animal experiments were approved by the Austrian Animal Care and Use Committee and Sing Health Institutional Animal Care and Use Committee in Singapore (2019/SHS/1495, 2020/SHS/1556).

### Behavior assays

Mice were transferred to the preclinical phenotyping facility of the Vienna Biocenter Core Facilities GmbH (VBCF) at least 1 week prior to experiments and housed at a 14hours light-10hours dark cycle in IVC racks, with access to food and water *ad libitum*. At the start of each experiment, mice were allowed to habituate to the experimental room for at least 30 minutes prior to any testing.

### Open field

Naïve mice were allowed to explore an open field arena (www.tse-systems.com) sized 50cm (width)x50cm (length)x29.5cm (height) for 30min and were video tracked using TSE VideoMot 3D software (version 7.01). In the software, the “center” zone was defined as a central square 25 cm x 25cm in size, the rest being the “border zone”. Light conditions were about 100 lux in the center zone. The time spent in each zone, distance travelled, number of center visits and rearing were recorded as readout parameters. All open field tests were performed during the morning (9am-1pm).

### Rotarod

Up to four mice were placed simultaneously on the rod compartments of a RotaRod device (model no. 3375-M5)—developed by TSE-Systems (Germany) and operated by TSE RotaRod v4.2.5 (4695) software,— first without any rotations (0rpm) facing the experimenter and habituated to sitting on the rod for up to 1min or until they fell (pre-trial 1). During pre-trials, 2 or 3 mice were placed on the rod and allowed to rotate at 4rpm, facing the experimenter (running forward) for up to 1min or until they fell. During trials, 1-4 mice were placed on the rod and allowed to rotate at 4rpm first and then accelerated from 4 to 40rpm, for up to 5min or until they fell. Passive rotations (clinging onto the rod without running) were treated like a fall. There was a 15 min break between each pre-trial and trial. The latency period until the mice fell and the speed at which they fell were recorded via TSE RotaRod v4.2.5 (4695) software. All rotarod tests were performed in the morning (9am-1pm).

### Beam walk

Mice were trained to walk from the home cage across a self-made wooden beam (cylindrical with, 20mm diameter, 1m length, 33cm height above the floor) to reach a wooden platform with a red igloo over 3 consecutive days, by repeatedly placing them at the end of the beam and letting them traverse the beam for as many repeats as achieved within 3 minutes. On day 4, all mice were tested using the 20mm diameter beam, on which they were allowed to traverse until they reached the platform, or fell, or the 60sec duration was completed (1 trial). When all the mice were tested on the 20mm diameter beam, the same procedure was repeated for the 16mm-, 14mm-, and 10mm-diameter beams. Following a four-day interval, training was resumed on Day 8, where the mice were tested on different diameter beams. All training and test sessions were recorded with a Sony Handycam HDR-CX200, and the behavior was evaluated by subsequent video analysis using “The Observer XT version 11.5” software by Noldus (The Netherlands); the distance reached, time needed to transverse, number of falls, slips, turns, stops and the time spent stopping were evaluated. The light conditions in the room were about 200 lux. All beam-walk experiments were performed during the afternoon (12pm-6pm).

### Elevated plus maze

Mice were placed in the center zone (6.5 x 6.5cm), facing an open arm of a custom-built elevated plus maze (elevated 54cm above the floor) that contains two open arms (OA, 30cm length, 7cm width) and two wall-enclosed arms (closed arms, CA, 30cm length, 6cm width, walls 14.5cm high) and allowed to explore freely for 5 minutes. Their path was video tracked using Topscan software (Cleversys, Inc., VA, USA) and the amount of time spent, and distance travelled in the open arms, closed arms and center zone were evaluated. Lux levels were about 100 lux in the center zone and open arms and about 35 lux in the closed arms.

### Y-maze

The Y-maze was performed as a test of working memory using a custom-built Y-shaped maze (made by workshop of Research Inst. of Molecular Pathology-IMP and Institute of Molecular Biotechnology-IMBA, Austrian Academy of Sciences, Vienna, Austria) with grey, opaque walls and floor and the following dimensions: arm length: 30cm, arm width: 6cm, wall height: 14.5cm. After 30 minutes of habituation to the test room (lux levels approx. 180 lux, visual cues on walls) mice were placed individually at the ends of one of the three arms (arm A, B, C), facing the wall at the end of the arm, and allowed to explore the maze for five minutes while being video tracked with the Topscan software (Cleversys Inc, USA). The experimenter watched the videos in the same room, behind a curtain and scored the latency to leave the starting arm (which was alternated between the mice) and the number and sequence of arm entries. This sequence was evaluated in triplets: Three arm entries in a row were scored as either correct spontaneous alternations (SA, e.g., BAC, CBC, ABC), erroneous alternate arm returns (AARs; e.g. BAB, CBC, ABA), or erroneous same arm returns (SARs, e.g. BAA, CCB, AAC). After each triplet was scored, the start of the analysis was shifted by 1 entry and the next triplet sequence was scored. Two such shifts of analysis result in overlapping triplets and the scoring of all possible decision points of the mouse. Spontaneous alternation performance (SAP) was calculated as [spontaneous alternations (SA)/(total arm entries – 2)].

### Light Dark transition test

**The** mice were placed in the dark zone and allowed to explore the light-dark arena (open field arena from TSE-Systems modified with custom-built dark zone boxes) freely for 20 minutes. Their path was video-tracked in the light zone using Videomot 2 software (TSE-Systems GmbH, Germany) and the amount of time spent in the light versus dark zones, the distance travelled in the light zone, as well as the latency until they transitioned to the light zone for exploration were evaluated. Lux levels were about 400 lux in the light zone and about 0 lux in the dark zone. Each zone (light zone and dark zone) was 24.5cm x 50cm in size.

### Grip strength test

Grip strength was measured using a grip strength meter (Bioseb, USA). For the forelimb measurement, the mouse was gently lowered over the top of a grid so that only its front paws could grip the grid. When the animal was gently pulled back and when it released the grid (which is connected to a sensor), its maximal grip strength value of the animal was displayed on the screen and noted. For the forelimb and hind limb measurement, the mouse was gently lowered over the top of the grid so that both its front and hind paws could grip the grid. The torso was kept parallel to the grid and the mouse was pulled back gently and steadily (not jerking), until it released its grip down the complete length of the grid. The maximal grip strength value of the animal was displayed on the screen and noted. Both, “forelimb only” and “fore- and hindlimb” tests were performed in an alternating fashion three times/mouse with a 15-minute inter-trial interval and the values were averaged across the three trials.

### Compounds

Sepiapterin (Sep, 11.225) was purchased from Schircks Labs, Switzerland. QM385 was developed previously and used as instructed (Cronin et al., 2018). SPRi3 developed previously and used as instructed (Latremoliere et al., 2015). Retinoic acid (Sigma, R2625); PMA (Sigma, P1585); MPP^+^ (Sigma, D048); Carbidopa (Sigma, C1335); Benserazide (Sigma, B7283); MPTP (Cayman Chemicals, 16377).

### Culturing and differentiating SHSY5Y cells

SHSY5Y cells (ATCC, CRL2266) were seeded at a concentration of 2000 cells/well in 500 µl DMEM/F12 10% FBS medium in a TC 48-well plate (Nunclon). The next day, 500 µL of pre-warmed DMEM/F12 10% FBS + 2X retinoic acid media was added and cells incubated for an additional three days. The media was changed to prewarmed DMEM/F12 10% FBS + PMA media and incubated for another three days to induce differentiation. The cells were then treated with 1mM MPP^+^ and sepiapterin and mitochondrial integrity was analyzed with Mitotracker Red CMXRos (Cell Signaling, 9082) 24 hours later, and survival assessed 48 hours later after compound addition.

### Targeted Metabolomics

Metabolites were extracted from tissue or cell pellets using a MeOH:ACN:0.1% DTT in water (2:2:1, v/v) (MeOH= Methanol; ACN=Acetonitrile) ice-cold solvent mixture. 500 μL of the solvent were added to 50 mg of tissue in an Eppendorf tube, vortexed for 30 s, incubated in liquid nitrogen for 1 min, and subjected to vigorous vortex shaking for 5 min. Then, the cells were disrupted using a pellet mixer (2 min) at low temperature, followed by centrifugation at 4,000 × g for 10 min at 4 °C. The supernatant was collected and transferred to another tube. 100 μL MeOH:ACN: 0.1% DTT in water (2:2:1, v/v) were added to the pellet, which was homogenized again using a pellet mixer for 2 min, following which it was centrifuged at the same conditions as described above. After that, both supernatants were combined (approx. 600 μL), centrifuged again at 4,000 × g for 10 min at 4 °C, and transferred to a new tube. Reversed phase liquid chromatography-tandem mass spectrometry (LC-MS/MS) was used for the quantification of BH4, sepiapterin, dopamine, neopterin and serotonin. Briefly, 1 μl of the extract was injected on a RSLC ultimate 3000 (Thermo Fisher Scientific) directly coupled to a TSQ Vantage mass spectrometer (Thermo Fisher Scientific) via electrospray ionization. A Kinetex C18 column was used (100 Å, 150 x 2.1 mm) at a flow rate of 80 µl/min. LC-MS/MS was performed by employing the selected reaction monitoring (SRM) mode of the instrument using the transitions (quantifiers) 242.1 *m/z* → 166.1 *m/z* (BH4); 238.1 *m/z* → 192.1 *m/z* (sepiapterin); 154.1 *m/z* → 137.1 *m/z* (dopamine); 254.1 *m/z* → 206.1 *m/z* (neopterin) and 177.1 *m/z* → 192.1 *m/z* (serotonin) in the positive ion mode. A 7-minute-long linear gradient was used, starting from 100% A (1 % acetonitrile, 0.1 % formic acid in water) to 80% B (0.1 % formic acid in acetonitrile). Freshly prepared dithiothreitol (1mg/ml) was used for stabilizing BH4. Authentic metabolite standards (Merck) were used for determining the optimal collision energies for LC-MS/MS and for validating experimental retention times. ATP, ADP, pyruvate, and lactate levels were measured using HILIC (hydrophilic interaction chromatography). Each sample was injected, at a flow rate of 100 µl/min, onto a SeQuant ZIC-pHILIC HPLC column (Merck, 100 x 2.1 mm; 5 µm) operated with an Ultimate 3000 HPLC system (Dionex, Thermo Fisher Scientific) and directly coupled to a TSQ Quantiva mass spectrometer (Thermo Fisher Scientific). A 15-minute gradient to 100% B (A: 95% acetonitrile 5% 10 mM aqueous ammonium acetate; B: 50% acetonitrile 50% 10 mM aqueous ammonium acetate) was used for separation. The following transitions were used for quantitation in the negative ion mode: 506.1 *m/z* → 159.1 *m/z* (ATP); 426.1.1 *m/z* → 134.1 *m/z* (ADP); 89.1 *m/z* → 43.1 *m/z* (lactate); 87.1 *m/z* → 34.1 *m/z* (pyruvate). The total intensity of the ion counts of a specific transition is proportional to the amount of that metabolite. It should be noted that throughout the project, various labs were involved in measuring BH4 and related monoamines and hence the reason for different labels. With the LC-MS/MS described above, the area under the curve (AUC) for the peak of the respective metabolite was used as comparative analysis. Within each preparation the results can be comparatively analyzed but not between different experiments due to the use of different machines and day-to-day variations in metabolite preparations.

### Nuclear magnetic resonance (NMR) spectroscopy

1H NMR experiments were carried out at 298K on a Bruker Avance III HD+ 600 MHz spectrometer equipped with a TXI probe. The spectra were measured using a Watergate experiment with 64 scans and 3-minute measurement time. The samples contained 500 μM compounds, 5 % DMSO, 90% PBS and 5% DMSO-d6 for the lock signal.

### Brain embedding and sectioning

A coronal cut was made on the mouse brains ventral to the median eminence with a sharp razor blade; the anterior part was frozen for dissecting the striatum on dry ice and the posterior part postfixed in 4% paraformaldehyde in phosphate buffered saline (PBS) at 4°C for approximately 2 weeks, washed twice with PBS and left in PBS at 4°C until it was embedded into paraffin. For embedding, brains were washed with ice-cold PBS, the majority of the cerebellum was cut away and the brain was dehydrated and paraffinated automatically (Miles Scientific Tissue-TEK VIP) using increasing concentrations of ethanol followed by xylol and paraffin. Paraffin-embedded midbrain was cut into 20 μm slices using a microtome (Reichert). When no cerebellum was visible under the microscope, slices were mounted to Starfrost ® microscope slides. When the medial habenular nucleus could be seen under the microscope, mounting of the slices was stopped. The slices were fixed on the microscope slides by heating to 60 °C for 4 h. For cryoembedding, brans were extracted and fixed in 4% PFA for two days at room temperature. Brains were then transferred to 30% sucrose solution for 4 additional days and embedded in OCT (Tissue-tek). For midbrain sectioning, Bregma range −2.68 to −3.79 was used; paraffin section thickness was 3-4 μm while cryo sections were 10-12 μm.

### Histology staining

20-μm thick cryosections were blocked with 1% bovine serum albumin (Sigma-Aldrich)/ 0.1%Triton X-100 in 0.1 M phosphate buffered saline (PBS) and then incubated with primary antibodies overnight at 4°C. After 3 washes in PBS for 10 minutes each, sections were incubated with secondary antibody for 1 hour at room temperature, washed 3 times in PBS (10 minutes each) and mounted using Vectashield Plus with DAPI mounting medium (H-2000, Vectashield). Primary antibodies and detection reagents used: tyrosine hydroxylase (anti-rabbit, Abcam, ab137869; anti-mouse, Millipore, MAB318), dopamine transporter (Abcam, ab184451), dopamine decarboxylase (Sigma, HPA017742), anti-VMAT2 (Progen, 16085), anti-4-HNE (JaICA, MHN-100P). Secondary antibodies used: Alexa Fluor 488 anti-rabbit (A21206, Invitrogen), 1:500; Alexa Fluor 555 anti-mouse (A32773, Invitrogen), 1:500 and Alexa Fluor 555 anti-rabbit (A32794, Invitrogen), 1:500. All images were assembled for publication using Fiji. For comparative analysis, fluorescence intensity, exposure time, and other parameters were consistent for all conditions in the same experiment.

### GCH1 enzymatic activity assay

GCH1 activity was measured spectrophotometrically (Infinite M Plex, Tecan, CA, USA), according to Kolinski and Gross (Kolinsky and Gross, 2004), with some modifications. Briefly, protein solutions were diluted in fresh 50 mM Tris-HCL buffer pH 7.5. The activity was assessed by following the formation of 7,8-dihydroneopterin triphosphate (NH2) in the presence of 2 mM GTP and 0.1 mg/mL of protein at 340 nm. The extinction coefficient of ɛ340 = 1820 M^−1^ cm^−1^ was used for the calculations after correction for multi-well plates. Activities are depicted as mU / mg protein, and 1U corresponds to 1 μmol formed NH2 / min at 37°C.

### Mitochondrial respiration

Mitochondrial respiratory parameters were measured with high-resolution respirometry (Oxygraph-2k, Oroboros Instruments, Innsbruck, Austria) (Mkrtchyan et al., 2018) by incubating striatal tissue homogenate in a buffer containing 115 mM KCl, 5 mM KH_2_PO_4_, 20 mM Tris-HCl, 1 mM diethylenetriamine penta acetic acid, and 1 g/L fatty acid-free bovine serum albumin at 37°C (pH 7.2). Complex I-linked State 2 respiration was induced by addition of 5 mM glutamate/5 mM malate. Transition to State 3 respiration was induced by addition of 1 mM adenosine diphosphate. Total capacity was measured by titration of carbonyl cyanide-4-(trifluoromethoxy)-phenylhydrazone in steps of 0.5 µM. Respiration rates were obtained by calculating the negative time derivative of the measured oxygen concentration.

### Mitochondrial tracking

DRG neuronal cell bodies were infected at seeding with a Lentivirus-expressing mitochondria-targeted mEOS2 (titer 8.2 × 109 cfu/ml) using a multiplicity of infection (MOI) of 20. Live imaging of mitochondria in DRG axons was performed at DIV14 using a Zeiss LSM 880 confocal microscope equipped with Airyscan, incubation chamber and a 20x water dipping objective (Zeiss Plan-Apochromat 1.0 NA W DIC objective, Zeiss, Germany). Untreated control DRGs or DRGs treated with the BH4 activator Sepiapterin (5µM) and inhibitor (QM385, 5µM) for 4 days were imaged in this experiment. Mitochondria in the most proximal 50 µm segment of the DRG axon were photoconverted using a 405 nm laser at 3% laser power for 20 s. Immediately after photoconversion, the 85 µm-long proximal axonal segment was imaged every 15 seconds for 6 min. An 8–12-μm stack was created from the images at every 0.5 μm in depth. Time lapse images, exported from the Zeiss software, were used to generate videos, while all further analysis was done in Fiji(Schindelin et al., 2012). Newly transported mitochondria (in green), either appearing from the neuronal cell body to the most proximal 20 μm axonal segment or distal axonal mitochondria appearing in the 20 μm-long axonal segment of the most distal photoconverted segment, were counted visually and confirmed using the kymograph. Kymographs of the time lapse images were generated by using the ImageJ plugin KymographClear2.0 (Mangeol et al., 2016).

### hMLO electrophysiology of brain slices

Whole-cell patch-clamp recordings of neurons from acute mouse midbrain slices and hMLOs (wholemount) were performed. Midbrain slices were prepared from C57BL/6 mice (Jax) at postnatal day 60-90 as previously described(Guzman et al., 2018). After the mice were anaesthetized by inhalation with 5% isoflurane, they were decapitated and their brains were quickly removed and chilled in an ice-cold high-Mg^2+^ cutting solution containing the following (in mM): 110 mM choline chloride, 25 mM NaHCO_3_, 20 mM glucose, 2.5 mM KCl, 1.25 mM NaH_2_PO_4_, 1 mM sodium pyruvate, 0.5 mM CaCl_2_, 7 mM MgCl_2_, 0.57 mM ascorbate; the pH of the solution was adjusted to 7.4 by saturating with carbogen (95% O_2_–5% CO_2_), and was maintained at an osmolality of approximately 300 Osmol L^−1^. The isolated brain was glued onto the stage of a vibrating blade of Compresstome (VF-200, Precisionary), and 320 μm-thick slices were cut. The resulting slices were incubated at 34 °C for 30 min in artificial cerebrospinal fluid (aCSF) containing the following: 125 NaCl mM, 25 NaHCO_3_ mM, 20 mM glucose, 2.5 mM KCl, 1.25 mM NaH_2_PO_4_, 1 mM sodium pyruvate, 1 mM CaCl_2_, 0.5 mM MgCl_2_, and 0.57 mM ascorbate, and bubbled with 95% O_2_–5% CO_2_, and thereafter maintained at room temperature. SNpc neurons were visually identified (Olympus BX51WI upright microscope equipped with a 60x water-immersion objective lens with DIC-IR) based on size, somatodendritic morphology, and the presence of regular spiking with a frequency between 1 and 4 Hz(Guzman et al., 2018). Whole-cell patch-clamp recordings (one cell per slice or organoid) were performed at 32 ± 1 °C, and the rate of aCSF perfusion was maintained at 1–1.5 ml min^−1^. Recordings were obtained using a Multiclamp 700B amplifier, a Digidata 1550 digitizer, and pClamp 10 acquisition software (Molecular Devices, San Jose, CA, USA). Recorded signals were sampled at 10 kHz and filtered at 2 kHz. Patch pipettes for current-clamp mode were filled with internal solutions containing the following components: 130 mM K-gluconate, 20 mM KCl, 10 mM Na2-phosphocreatine, 10 mM HEPES, 0.1 mM EGTA, 2 mM Mg-ATP, and 0.3 mM NaGTP. We recorded series resistance throughout the experiments and excluded neurons with series resistance > 20 MΩ from data analysis. Membrane potential values were presented as recorded without correcting for liquid junction potentials. Perforated-patch experiments were performed with pipette fronts filled with a solution containing 130 mM KMeSO_4_, 10 mM NaCl, and 10 mM HEPES and then back-filled with the same solution containing 20 mg/ml gramicidin-D(Guzman et al., 2018).

### Intracranial stereotactic injections

Wild-type and *Gch1^fl/fl^* male mice (2-3 months old) were anaesthetized with isoflurane (5%, Abbot Laboratories) and fixed in a stereotaxic frame (Kopf). Anesthesia throughout the procedure was delivered via mouth tubing and maintained between 1.6 % and 2.2% of isoflurane in the air. The body temperature of the mouse was monitored with a rectal thermometer and kept constant at 36°C using a heating pad (DC temperature controller). The mouse head was shaved to expose the skull, following which it was cleaned and drilled with small holes bilaterally above the SN, at coordinates: AP −2.8, ML ± 1.3, DV −4.4, relative to bregma (Paxinos and Franklin, 2001). The mice were injected with 120nl/injection site of *AAV5; Th-Cre* (1 × 10^12^ GC/mL) (v659, Viral Vector Facility, ETH, Switzerland) viral vectors. Vectors were delivered to the SN via a thin glass pipette and the injection was controlled by a Micro4 Micro Syringe Pump (World Precision Instruments; injection volume 120 nl/site, delivery rate 30 nl/min). To prevent backflow after injection, the pipette was left at the injection site for 5 min. The mice were let to recover for at least one week after the surgery, during which time their drinking water was supplied with enrofloxacine 0.1mg/mL (Baytril, Bayer) and caprofen 0.2mg/mL (Rimadyl, Pfizer).

### LRRK2 patient fibroblast assay

Previously described (Smith et al., 2016, Korecka & Thomas et al., 2019) deidentified healthy and LRRK2 G2019S patient fibroblasts were obtained from the Coriell and the NIA Aging Cell biorepositories or the Mayo clinic. Conducting research with patient fibroblasts was not considered to be human subject research as it does not involve intervention or interaction with the individual and the samples do not contain identifiable private information. The average age of the control and the mutant fibroblasts was matched to 66 and 67 years, respectively. Mitochondrial super oxide (MitoSOX™ Red, ThermoFisher Scientific, M36008) levels, induced by 24-hour treatment with the mitochondrial depolarization agent valinomycin (10μM, SigmaAldrich, V0627), were measured in human fibroblasts obtained from healthy individuals (N=4) and PD patients carrying a LRRK2 G2019S mutation (N=5), in the presence of sepiapterin or SPRi3 (pretreatment for 24-hours before addition of valinomycin), using the live cell imaging system IncuCyte.

### Protein blotting

Protein blotting was carried out using standard protocols. Blots were blocked for 1 h with 5% milk in PBST (1× PBS and 0.1% Tween-20) and were then incubated overnight at 4 °C with primary antibodies diluted in 5% milk in PBST (1:1,000 dilution). Blots were washed three times in PBST for 15 min and were then incubated with HRP-conjugated secondary antibodies (1:2,500 dilution; GE Healthcare, NA9340V) for 45 min at room temperature, washed three times in PBST for 15 min and visualized using enhanced chemiluminescence (ECL Plus, Pierce, 1896327). The following antibodies were used in the present study: anti-TH (Abcam, ab137869; Millipore, MAB318); anti-DAT (Proteintech, 22524-1-AP); anti-DDC (Sigma, 91089); anti-VMAT2 (Abcam, ab259970); and monoclonal anti-mouse actin (Sigma, A5316).

### Reverse transcriptase quantitative polymerase chain reaction (RT-qPCR)

Total RNA was extracted from the ventral midbrain tissue of mice with the indicated genotypes. RNA extraction was performed using the RNAeasy Mini Kit (Qiagen) and included DNase I (TurboDNase) digestion to avoid potential DNA contamination. Then, 3 μg of total RNA was reverse transcribed using LunaScript cDNA Synthesis Kit (New England Biolabs). Gene-expression levels were quantified by real-time quantitative PCR (iCycler iQ BioRad) using the Luna qPCR Mastermix (New England Biolabs) and normalized to *Gapdh* housekeeping gene. mRNA fold changes were calculated using the ΔΔ*C*t method. The following mouse primer pairs were used:

GTP cyclohydrolase I (*Gch1*):

ACAAGCAAGTCCTTGGTCTCA; GTGAGGCGCTCTTGAACTTG

Glyceraldehyde 3-phosphate dehydrogenase *(Gapdh)*:

CATCACTGCCACCCAGAAGACTG; ATGCCAGTGAGCTTCCCGTTCAG

Tyrosine hydroxylase (*Th*):

CGGTGTACTGGTTCACTGTG; CGCATGCAGTAGTAAGATGT

Dopamine transporter (*Dat*):

GGTGCTGATTGCCTTCTCCAGT; GACAACGAAGCCAGAGGAGAAG

Vesicle monoamine transporter 2 (*Vmat2*):

CCTCTTACGACCTTGCTGAAGG; GCTGCCACTTTCGGGAACACAT

Dopamine decarboxylase (*Ddc*):

GGAGCCAGAAACATACGAGGAC; GCATGTCTGCAAGCATAGCTGG

### Patients

Outpatients with PD from the Public Movement Disorders Clinic of Santa Catarina State at the Hospital Governador Celso Ramos, Florianópolis city, southern Brazil were included in this study. Diagnosis was based on the Movement Disorders Society Guidelines (Goetz et al., 2008). Patients had bradykinesia, and at least 2 of the following symptoms: 1) resting tremor (frequency between 4 and 6 Hz); 2) muscle rigidity; and 3) postural instability not caused by visual or vestibular or cerebellar proprioceptive dysfunction. All patients had unilateral resting tremors onset; and showed substantial improvement in symptoms in response to dopamine agonists or levodopa treatment. The disease stage was evaluated using the Hoehn and Yahr scale (Hoehn and Yahr, 1967). Anxiety and depressive symptoms were determined by the Hospital Anxiety and Depression scale (Mehta et al., 2018) validated for Brazilian patients (de Lemos Zingano et al., 2015; Zingano, 2019). Exclusion criteria used for PD diagnosis were as follows: previous history of cerebrovascular disease, brain trauma, infection or tumor, oculogyric crisis, neuroleptic use, prolonged symptoms remission, strictly unilateral symptoms after 3 years of disease, supra-nuclear palsy, cerebellar symptoms, early onset dysautonomia or dementia, pyramidal signs, brain tumor, and hydrocephalus, use of anti-inflammatory medication (steroids or non-steroids) in the last 3 weeks; acute disease in the last 60 days; be not in the best “on” state (Mehta et al., 2018). Blood samples were collected from all patients at their first visit in the research study. Healthy persons (without neurologic or psychiatric diseases) were matched for age, gender, and education level (p > 0.70). The project was approved by the Ethics Committee (CEP-HGCR 2009-26) and signed informed consent was obtained from all participants.

### Blood and plasma collection

After the collection of clinical and psychological data, 4 mL of blood were drawn from the median cubital vein and distributed into sterile vacutainer tubes containing 10 % EDTA. The blood was immediately centrifuged at 400 × *g* for 10 min at room temperature and plasma was collected for biochemical analyses. For BH4 quantitation, 200 μL of plasma were immediately added to one volume (1:1, v/v) of 5 % trichloroacetic acid and centrifuged at 16,000 × *g* for 10 min at 4 °C. The supernatant was frozen and stored at – 86 °C. Plasma BH4 levels were analyzed immediately (within 6 h).

### Measurements of BH4 levels in patients

BH4 levels were determined by high performance liquid chromatography (HPLC) and quantified using electrochemical detection as previously described with some modifications (Latini et al., 2018). Plasma samples were thawed on ice and protected from light. Afterwards, samples were centrifuged (16,000 × *g*; 10 min; 4 °C) and 20 μL of supernatant were transferred to an HPLC vial for analysis. The HPLC analysis of BH4 was carried out in a HPLC system (Alliance e2695, Waters, Milford, USA) by using a Waters Atlantis dC18 reverse phase column (4.6 × 250 mm; 5 μm particle), with the flow rate set at 0.7 mL/min and an isocratic elution of 6.5 mM NaH_2_PO_4_, 6 mM citric acid, 1 mM sodium octyl sulfate, 2.5 mM diethylenetriaminepentaacetic acid, 160 μM dithiothreitol and 8 % acetonitrile, at pH 3.0. The temperature of the column compartment was set at 35 °C. The identification and quantification of BH4 was performed by an electrochemical detector (module 2465, Waters, Milford, USA) using a voltage of +450 mV. The results were expressed as ηmol/L.

### Measurements of neurotransmitters levels in patients

Catecholamines levels were determined by HPLC and quantified by using electrochemical detection as previously described (Scheffer et al., 2019). Plasma samples were precipitated by the addition of one volume (1:1, v/v) of 0.1 M perchloric acid containing 0.02 % sodium metabisulfite and centrifuged (16,000 × *g*; 10 min; 4 °C). Monoamines and metabolites present in the supernatants were assessed by HPLC (Alliance e2695, Waters, Milford, USA), with electrochemical detection (Waters 2465, Waters, Milford, USA). Twenty microliters of the supernatants were analyzed in a 150 mm x 2.0 mm, 4µm, C18 column (Synergi Hydro, California, USA) and an isocratic elution of 90 mM sodium phosphate, 50 mM citric acid, 2.3 mM sodium 1-heptane-sulfonate, 50 µM ethylenediaminetetraacetic acid, and 10 % acetonitrile, at pH 3.0 and with a flow rate of 0.20 mL/min. The results were expressed as ηmol/mL.

### Single-cell sequencing analysis

Data from Zeisel et al., 2018, which is the biggest (to our knowledge) scRNA-seq resource for the ventral midbrain published so far (9273 cells), was used for our analysis (Zeisel et al., 2018). Standard single-cell variance stabilizing transformation pipeline ((Hafemeister and Satija, 2019)– sctransform v. 0.3.2; Seurat v. 4.0.6) was used to correct unique molecular identifier (UMIs) counts used for visualization on density-preserving UMAP 2D-embedding (Kobak and Linderman, 2021) as *ln*(*x* + 1): normal logarithm with pseudo count 1 and to derive Pearson’s residuals (visualized as a heatmap). An impact of variability due to the role of *Gch1* in mitochondrial physiology was eliminated by regressing out the ratio between mitochondrial and ribosomal fractions of transcripts. Low resolution clustering ((Traag et al., 2019)– leidenalg v. 0.8.8) on full dataset was used to tidy up dopaminergic clusters (MBDOP1 and MBDOP2) in the original annotation. Our main interest in dopaminergic neurons of ventral tegmental area and substantia nigra represented in stable cluster MBDOP2 versus more variable MBDOP1 cluster of periaqueductal gray (PAG), which was used as control. To investigate the association of members of BH4-pathway we examined Pearson’s correlation with markers of dopaminergic neurons within these cell populations. We observed a highly significant correlation of *Gch1* with *Slc18a2 (Vmat2), Slc6a3 (Dat)* and *Th*. Also, we examined differential gene expression of *Gch1* positive versus *Gch1^-^* within strictly dopaminergic neurons (subset includes 13 MBDOP1 and 56 MBDOP2 neurons) using Wilcoxon rank sum test – co-expression subsets were visualized as UpSet plot (Conway et al., 2017). For differential comparison between *Gch1*-negative and *Gch1*-expressing midbrain DAergic neurons, GO overrepresentation analysis of top 250 genes (**Table S4**) was performed in R using clusterProfiler and analysis results were shown as an enrichment map in R using enrichplot.

### Computational analysis of published *GCH1* mutations

To compute how the various mutations in GCH1 affect the predicted vibrational energy and stability of the protein, the following software programs were used: mCSM (Pires et al., 2014a), DUET (Pires et al., 2014b), DynaMut (Rodrigues et al., 2018), ENCoM (Frappier et al., 2015), and SDM (Pandurangan et al., 2017).

### Statistical analyses

All values are expressed as mean +/- s.e.m and analysis performed by GraphPad Prism software. Details of the statistical tests used are stated in the figure legends. Briefly, Student’s t-test was used to compare two groups. One-way ANOVA followed by Dunnett’s post-hoc test was used for analysis between multiple groups. 2-way ANOVA was used to compare two groups over time. In all the tests, P ≤ 0.05 was considered significant.

## Notes

### Competing Interest Statement

The authors have declared no competing interest.

