## Supplemental data for "Crucial neuroprotective roles of the metabolite BH4 in dopaminergic neurons"

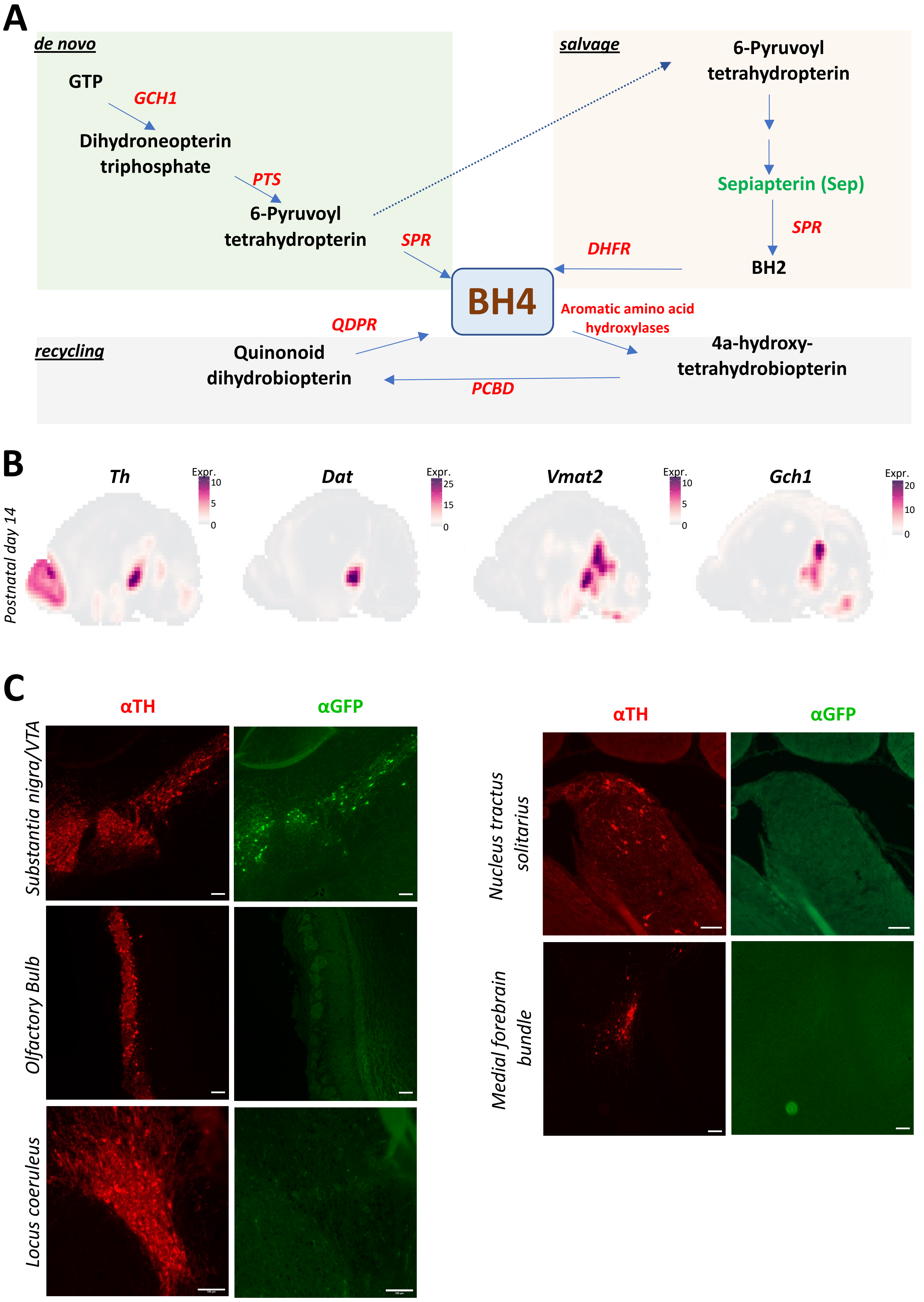
A
de novo
GTP
GCH1
Dihydroneopterin triphosphate
PTS
6-Pyruvoyl tetrahydropterin
SPR
salvage
6-Pyruvoyl tetrahydropterin
Sepiapterin (Sep)
SPR
BH2
DHFR
BH4
Aromatic amino acid hydroxylases
QDPR
recycling
Quinonoid dihydrobiopterin
4a-hydroxy-tetrahydrobiopterin
PCBD
B
Postnatal day 14
Th
Dat
Vmat2
Gch1
Expr.
10
5
0
25
15
0
20
10
C
αTH
αGFP
Substantia nigra/VTA
Olfactory Bulb
Locus coeruleus
Nucleus tractus solitarius
Medial forebrain bundle

Fig.S1

**Supplemental Figure Legends.**

**Supplemental Figure 1. *Gch1*-GFP expression is restricted to TH-expressing neurons of the ventral midbrain.**

**A,** Schematic representation of the de novo, salvage and recycling arms of the BH4 pathway. Dotted arrow indicates non-enzymatic reactions while solid arrows indicate enzymatic reactions. *GTP*, guanosine triphosphate; *PTPS*, 6-Pyruvoyl tetrahydropterin synthase; *DHFR*, dihydrofolate reductase; *SPR*, sepiapterin reductase; *QDPR*, quinoid dihydropteridine reductase; *PCDB*, pterin-4alpha-carbinolamine dehydratase. Sepiapterin (green) is an intermediatory metabolite of the salvage pathway and can be used to enhance BH4 levels. **B,** *In situ* intensities from the Allen Brain Atlas of various DAergic markers (*Th*, *Dat*, *Vmat2*) as well as *Gch1* in the brains of P14 mice. **C,** Representative brain images of prominent TH-expressing neuronal regions throughout the brain of *Gch1*-GFP reporter mice and costained with anti-GFP. Scale bar, 100µm.

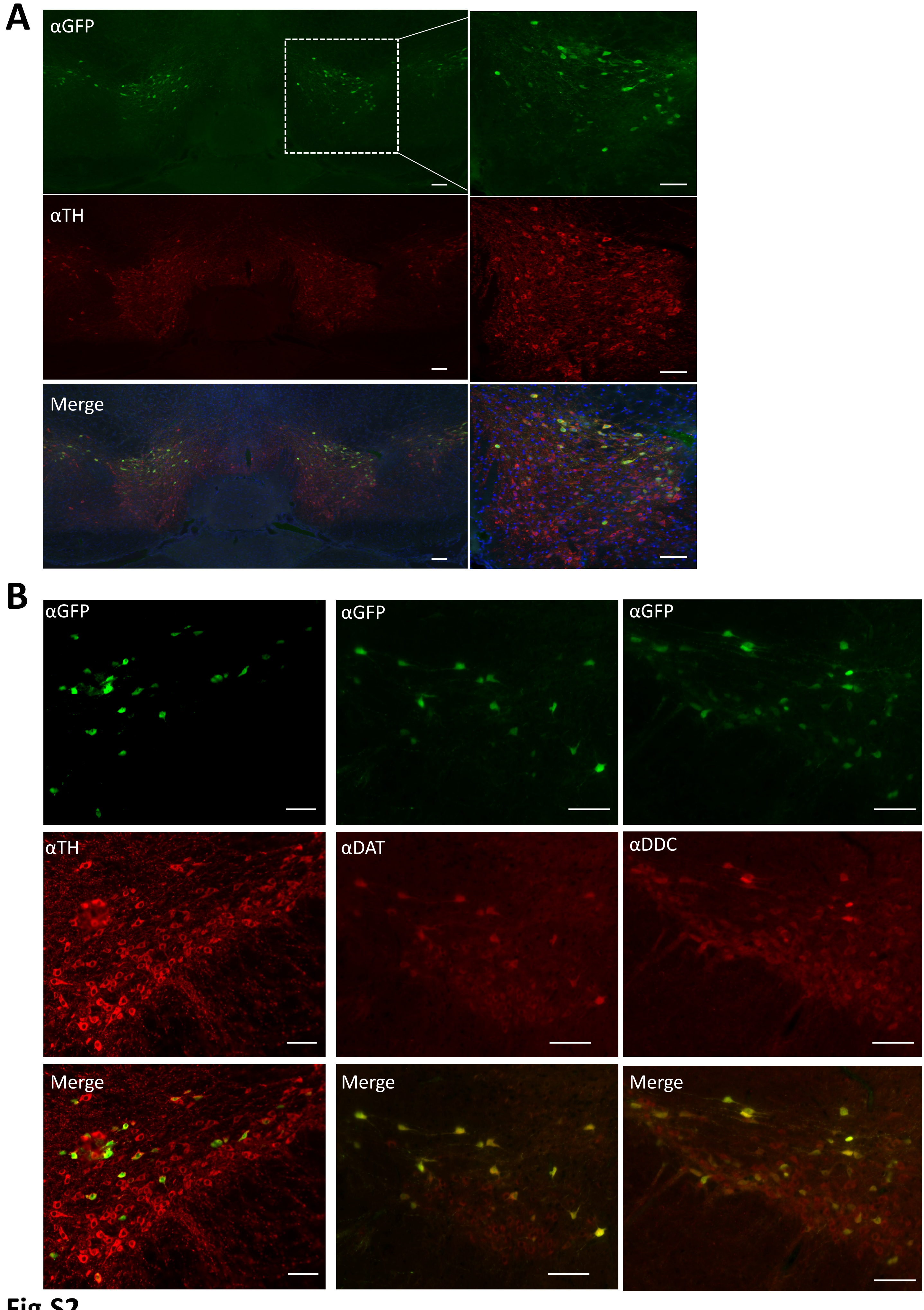
A
αGFP
αTH
Merge
B
αGFP
αGFP
αGFP
αTH
αDAT
αDDC
Merge
Merge
Merge

Fig.S2

**Supplemental Figure 2. *Gch1* is expressed in a subset of DAergic neurons.**

**A,** Representative images of coronal sections of the SNpc and VTA regions from *Gch1*-GFP reporter mice stained with anti-TH and anti-GFP. Scale bar, 100µm. **B,** Representative images of coronal sections of the SNpc from *Gch1*-GFP reporter mice stained with anti-TH, anti-DAT, anti-DDC and anti-GFP. Scale bar, 100µm.

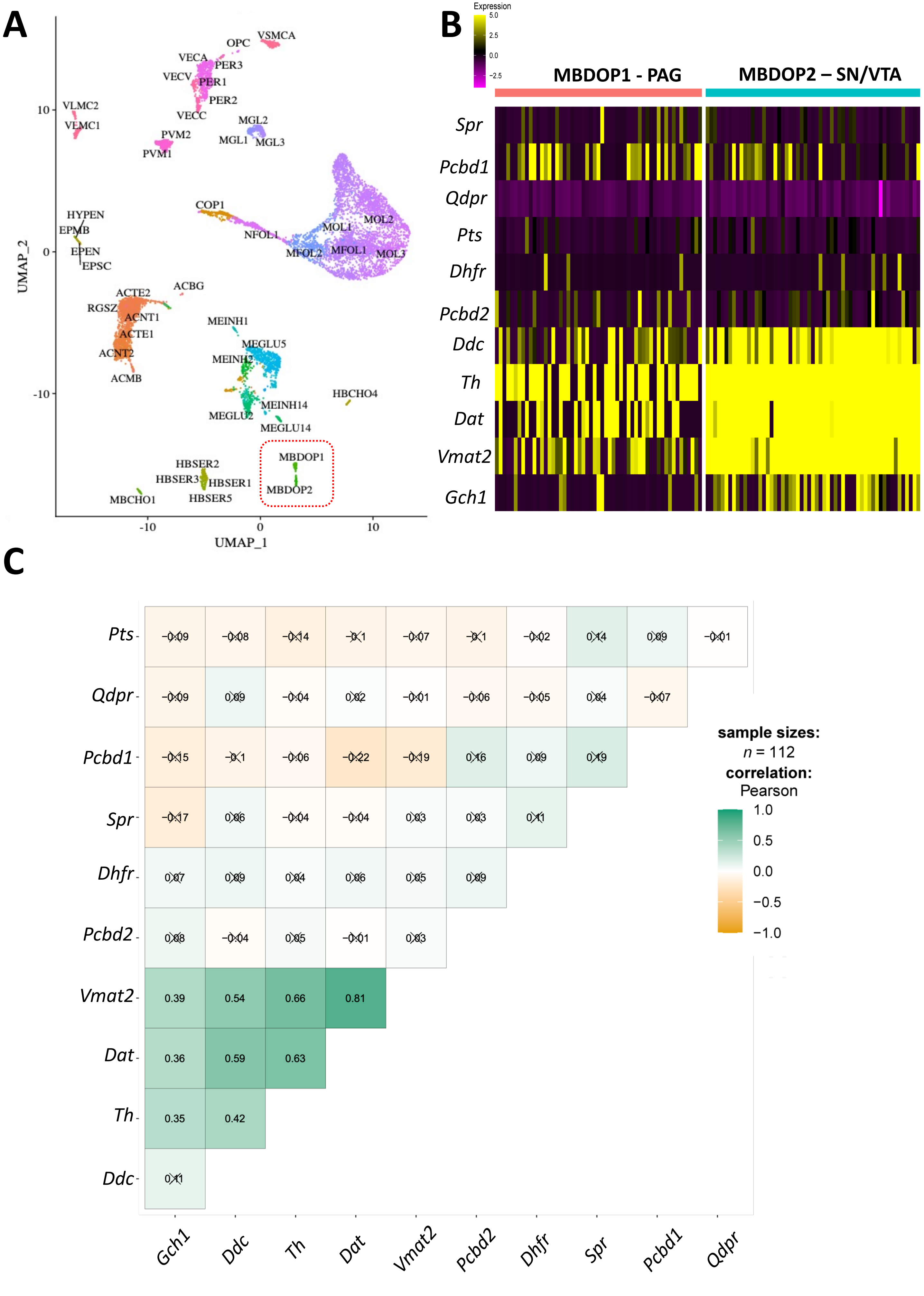
A
B
C
UMAP_1
UMAP_2
VSMCA
OPC
VECA
PER3
VECV
PER1
PER2
VLMC2
VEMC1
VECC
MGL2
PVM2
MGL1
MGL3
PVM1
COP1
HYPEN
EPMB
EPEN
EPSC
NFOL1
MOL1
MOL2
MFOL2
MFOL1
MOL3
ACBG
ACTE2
RGSZ
ACNT1
ACTE1
ACNT2
ACMB
MEINH1
MEGLU5
MEINH2
HBCHO4
MEINH14
MEGLU2
MEGLU14
MBDOP1
HBSER2
HBSER3
HBSER1
MBDOP2
HBSER5
MBCHO1
Expression
MBDOP1 - PAG
MBDOP2 – SN/VTA
Spr
Pcbd1
Qdpr
Pts
Dhfr
Pcbd2
Ddc
Th
Dat
Vmat2
Gch1
sample sizes:
n = 112
correlation:
Pearson

Fig.S3

**Supplemental Figure 3. Single cell RNAseq analysis of ventral midbrain DAergic populations reveal significance correlation between *Gch1* expression and SNpc/VTA DAergic neurons.**

**A,** DensMAP plot depicting original annotation by (Zeisel et al., 2018). Dotted red box contains the DAergic neurons of the PAG (MBDOP1) and the VTA/SN (MBDOP2). **B,C,** Heatmap of variance-stabilized expression levels as Pearson's residuals **(B)** and correlation matrix **(C)** of DAergic and BH4 pathways genes of original PAG and VTA/SN neuronal clusters annotation.

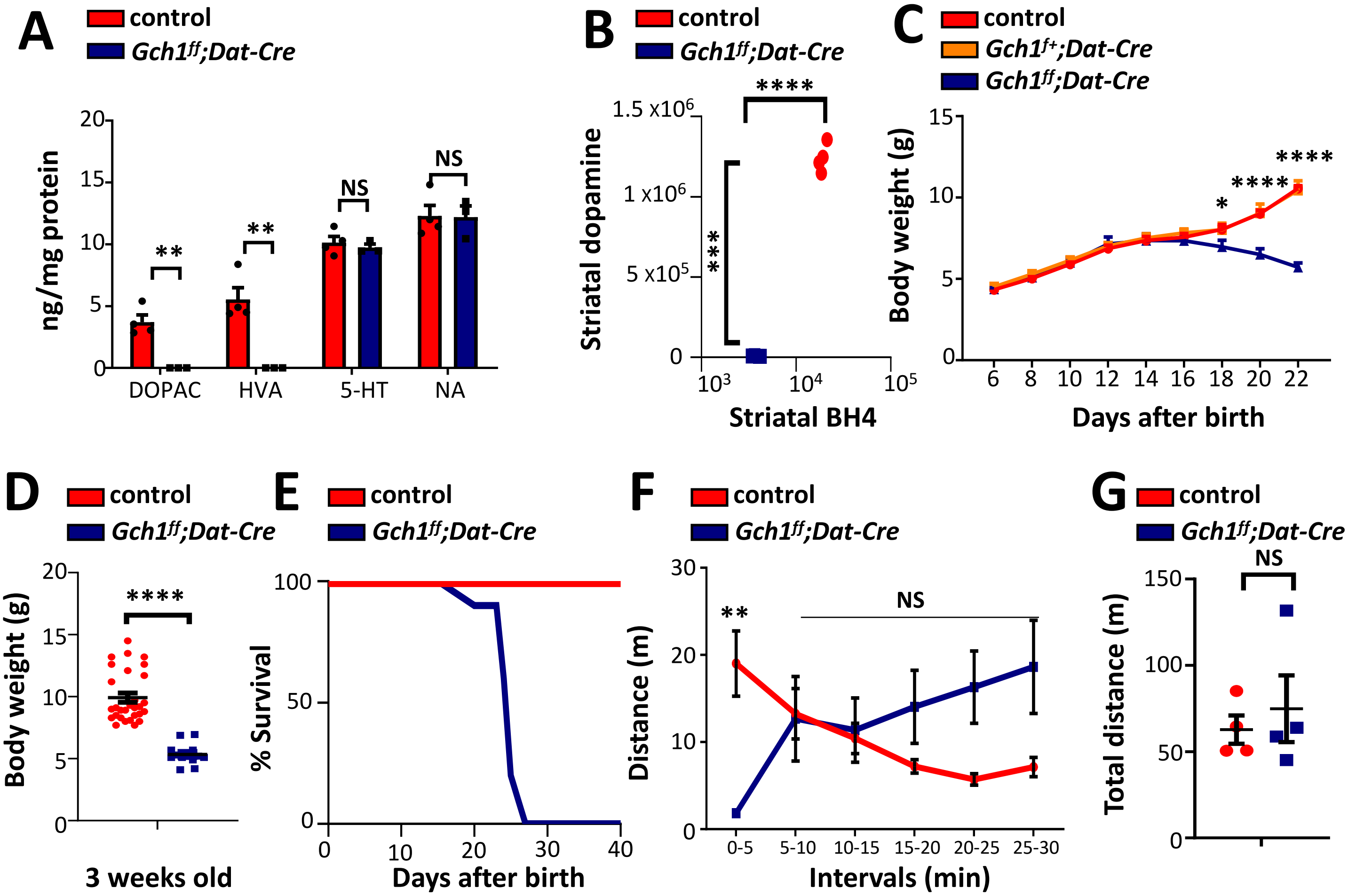

A
control
Gch1ff;Dat-Cre
ng/mg protein
DOPAC
HVA
5-HT
NA
**
**
NS
NS
B
Striatal dopamine
Striatal BH4
****
***
C
Gch1f+;Dat-Cre
Body weight (g)
Days after birth
*
****
****
D
Body weight (g)
3 weeks old
****
E
% Survival
Days after birth
F
Distance (m)
Intervals (min)
**
NS
G
Total distance (m)
NS


FIG.S4

**Supplemental Figure 4. Loss of DAergic neuron-specific *Gch1* results severe motor defects and premature death.**

**A,** Dopamine metabolites in the brain of control and *Gch1*$^{flox/flox}$*;Dat-Cre* mice. Data are shown as means ± s.e.m. Individual mice for each genotype are shown. **P < 0.01; NS, not significant (multiple t-test). **B,** Striatal BH4 and dopamine levels of control and *Gch1*$^{flox/flox}$*;Dat-Cre* mice. Data are shown as means ± s.e.m. Individual mice for each genotype are shown. ***P < 0.001; ****P < 0.0001; NS, not significant (Student's t-test). **C,D,** Body weights of control, *Gch1*$^{flox/flox}$*;Dat-Cre* and heterozygous *Gch1*$^{flox/+}$*;Dat-Cre* mice after birth (**C**) and at 3-weeks old age (**D**). Data are shown as means ± s.e.m. *P < 0.05; ****P < 0.0001 (Two-way ANOVA with Tukey's multiple comparisons). **E,** Survival curve of control and *Gch1*$^{flox/flox}$*;Dat-Cre* mice. **F,G,** Distance travelled of the open field testing in 5-minute intervals (**F**) and in 30 minutes (**G**) of 2-week old control and *Gch1*$^{flox/flox}$*;Dat-Cre* mice. Data are shown as means ± s.e.m. **P < 0.001; NS, not significant (Student's t-test with multiple comparisons).

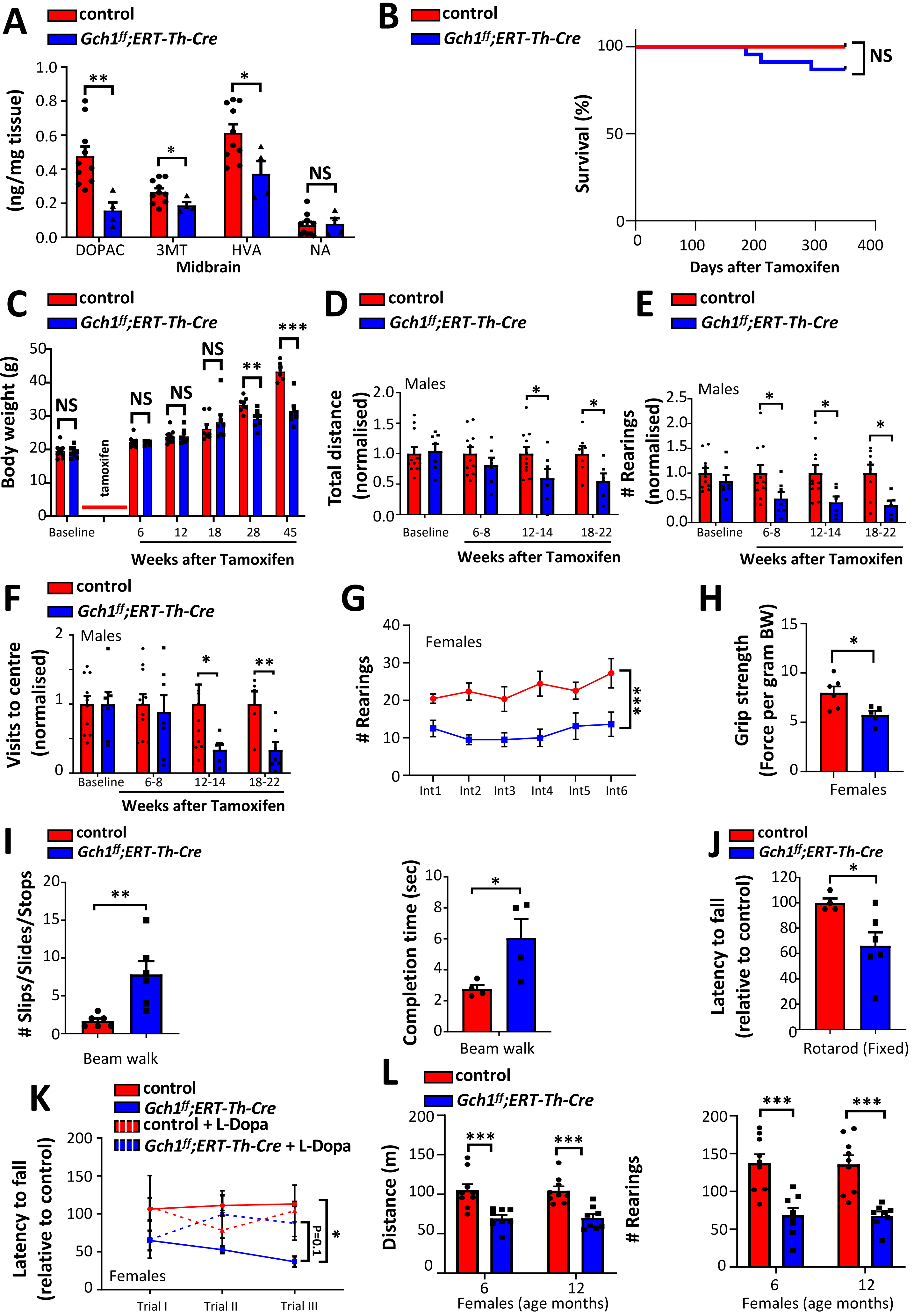
A
control
Gch1ff;ERT-Th-Cre
(ng/mg tissue)
DOPAC
3MT
HVA
NA
Midbrain
NS
B
control
Gch1ff;ERT-Th-Cre
Survival (%)
Days after Tamoxifen
NS
C
control
Gch1ff;ERT-Th-Cre
Body weight (g)
tamoxifen
Baseline
Weeks after Tamoxifen
D
control
Gch1ff;ERT-Th-Cre
Males
Total distance (normalised)
Baseline
6-8
12-14
18-22
Weeks after Tamoxifen
E
control
Gch1ff;ERT-Th-Cre
Males
### Rearings (normalised)
Weeks after Tamoxifen
F
control
Gch1ff;ERT-Th-Cre
Males
Visits to centre (normalised)
Weeks after Tamoxifen
G
Females
### Rearings
Int1
Int2
Int3
Int4
Int5
Int6
H
Grip strength (Force per gram BW)
Females
I
control
Gch1ff;ERT-Th-Cre
### Slips/Slides/Stops
Beam walk
Completion time (sec)
Beam walk
J
control
Gch1ff;ERT-Th-Cre
Latency to fall (relative to control)
Rotarod (Fixed)
K
control
Gch1ff;ERT-Th-Cre
control + L-Dopa
Gch1ff;ERT-Th-Cre + L-Dopa
Latency to fall (relative to control)
Females
Trial I
Trial II
Trial III
P=0.1
L
control
Gch1ff;ERT-Th-Cre
Distance (m)
Females (age months)
### Rearings
Females (age months)

**Supplemental Figure 5. Progressive motor dysfunction in tamoxifen-inducible *Gch1$^{flox/flox}$;ERT-Th-Cre* mice.**

**A,** Quantification of dopamine metabolites in the midbrains of control and *Gch1$^{flox/flox}$;ERT-Th-Cre* mice. Data are shown as means ± s.e.m. Individual mice for each genotype are shown. *P < 0.05 (multiple t-test). **B,** Kaplan-Meier survival curves of control and *Gch1$^{flox/flox}$;ERT-Th-Cre* mice after tamoxifen administration as indicated. NS, not significant (Log-rank (Mandel-Cox) test). **C**, Body weight measurements of male control and *Gch1$^{flox/flox}$;ERT-Th-Cre* mice after tamoxifen administration. Data are means ± s.e.m. Individual mice for each genotype are shown. **P < 0.01; ***P < 0.001 (multiple t-test). **D-F**, Open field behavioral testing of male control and *Gch1$^{flox/flox}$;ERT-Th-Cre* mice at indicated timepoints after tamoxifen administration for total distance travelled **(D)**, numbers of rearings **(E)** and visits to the center **(F)**, assessed over a 30-minute observational period. Data are means ± s.e.m. Individual mice for each genotype are shown. *P < 0.05; **P < 0.01; NS, not significant (Multiple t-test). **G**, Open field behavioral testing of female control and *Gch1$^{flox/flox}$;ERT-Th-Cre* mice after tamoxifen administration for numbers of rearings in 5-minute intervals assessed over a 30-minute observational period. Data are means ± s.e.m. Individual mice for each genotype are shown. ***P < 0.001 (Two-way ANOVA). **H**, Grip strength of all paws from control and *Gch1$^{flox/flox}$;ERT-Th-Cre* female mice 28 weeks post tamoxifen administration. Data are means ± s.e.m. Individual mice for each genotype are shown. *P < 0.05 (Student's t-test). **I,** Beam walk behavioral testing of control and *Gch1$^{flox/flox}$;ERT-Th-Cre* mice 28 weeks post tamoxifen administration. Number of stops, slips and slides during crossing (left) as well as the time taken to cross the 10cm-beam successfully (right) were measured. Data are shown as means ± s.e.m. Individual mice for each genotype are shown. *P < 0.05; **P < 0.01 (Student's t-test). **J,** Fixed (4 rotation per minute) rotarod analysis of control and *Gch1$^{flox/flox}$;ERT-Th-Cre* mice 28 weeks post tamoxifen administration. Data are shown as means ± s.e.m. Individual mice for each genotype are shown. *P < 0.05 (Student's t-test). **K**, Accelerated (4-40km/hr) rotarod analysis of control and *Gch1$^{flox/flox}$;ERT-Th-Cre* female mice 28 weeks post tamoxifen administration before and 45 minutes after administration of L-Dopa (50mg/kg) plus benserazide, (12.5mg/kg). Data are means ± s.e.m. Individual mice for each genotype are shown. *P < 0.05 (2-way ANOVA with Tukey's multiple comparison test).

**L,** Open field behavioral testing of female control and *Gch1*$^{flox/flox}$*;ERT-Th-Cre* mice at 6 and 12 months after tamoxifen administration for total distance travelled (left), and numbers of rearings (right) assessed over a 30-minute observational period. Data are means ± s.e.m. Individual mice for each genotype are shown. ***P < 0.001 (Multiple t-test).

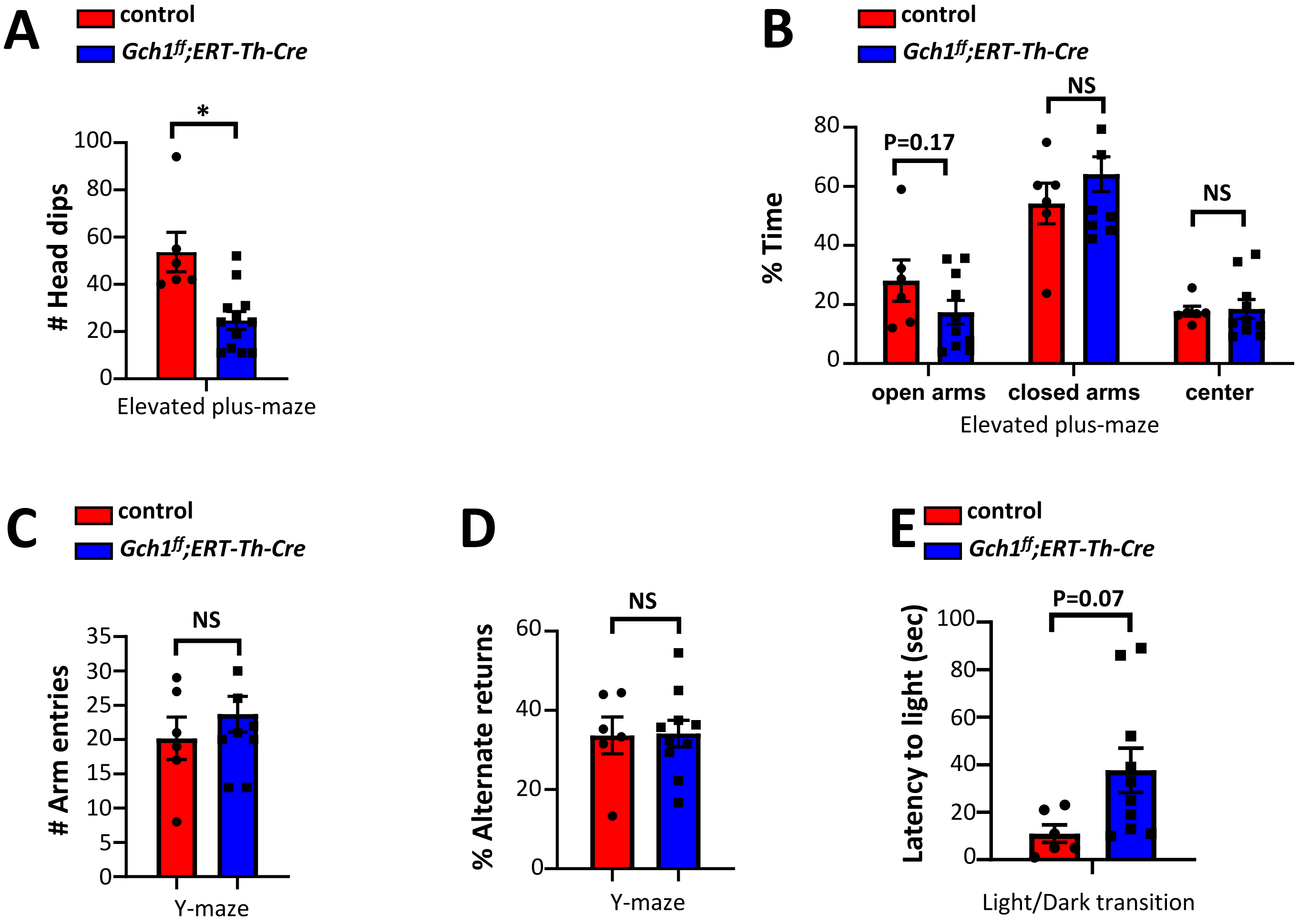

A
control
Gch1ff;ERT-Th-Cre
*
### Head dips
100
80
60
40
20
0
Elevated plus-maze
B
control
Gch1ff;ERT-Th-Cre
% Time
80
60
40
20
0
P=0.17
NS
NS
open arms
closed arms
center
Elevated plus-maze
C
control
Gch1ff;ERT-Th-Cre
NS
### Arm entries
35
30
25
20
15
10
5
0
Y-maze
D
NS
% Alternate returns
60
40
20
0
Y-maze
E
control
Gch1ff;ERT-Th-Cre
P=0.07
Latency to light (sec)
100
80
60
40
20
0
Light/Dark transition


Fig.S6

**Supplemental Figure 6. Increased anxiety-related behaviors observed in tamoxifen-inducible *Gch1^flox/flox^;ERT-Th-Cre* mice.**

**A,B,** Elevated plus maze behavior of control and *Gch1*$^{flox/flox}$*;ERT-Th-Cre* mice 28 weeks post tamoxifen administration looking at number of head dips in the open arms **(A)** and percent of time spent in open versus closed arms **(B)**. Data are shown as means ± s.e.m. Individual mice for each genotype are shown. *P < 0.05; NS, not significant (Student's t-test). **C,D,** Y-maze behavior testing in which number of arm entries **(C)** as well as percentage of alternate arm returns **(D)** were measured. Data are shown as means ± s.e.m. Individual mice for each genotype are shown. NS, not significant (Student's t-test). **E**, Light/Dark box behavioral analysis in which initial latency of movement into the light compartment were recorded. Data are shown as means ± s.e.m. Individual mice for each genotype are shown. NS, not significant (Student's t-test).

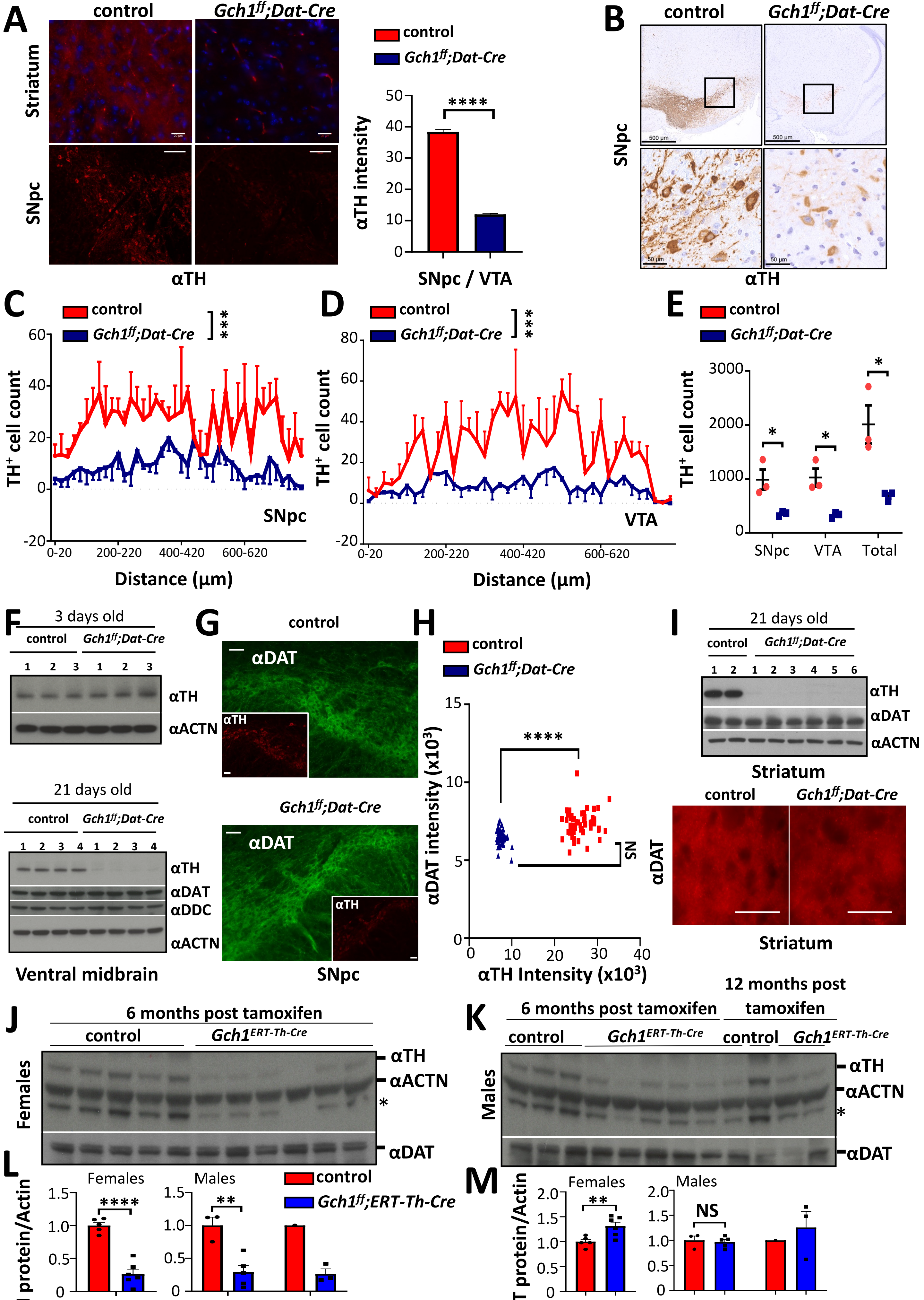
A
control
Gch1ff;Dat-Cre
Striatum
SNpc
αTH
control
Gch1ff;Dat-Cre
αTH intensity
****
SNpc / VTA
B
control
Gch1ff;Dat-Cre
SNpc
αTH
C
control
Gch1ff;Dat-Cre
***
TH+ cell count
SNpc
Distance (μm)
D
control
Gch1ff;Dat-Cre
***
TH+ cell count
VTA
Distance (μm)
E
control
Gch1ff;Dat-Cre
TH+ cell count
SNpc
VTA
Total
F
3 days old
control
Gch1ff;Dat-Cre
αTH
αACTN
21 days old
control
Gch1ff;Dat-Cre
αTH
αDAT
αDDC
αACTN
Ventral midbrain
G
control
αDAT
αTH
Gch1ff;Dat-Cre
αDAT
αTH
SNpc
H
control
Gch1ff;Dat-Cre
****
NS
αDAT intensity (x10³)
αTH Intensity (x10³)
I
21 days old
control
Gch1ff;Dat-Cre
αTH
αDAT
αACTN
Striatum
control
Gch1ff;Dat-Cre
αDAT
Striatum
J
6 months post tamoxifen
control
Gch1ERT-Th-Cre
αTH
αACTN
*
αDAT
Females
K
6 months post tamoxifen
12 months post tamoxifen
control
Gch1ERT-Th-Cre
control
Gch1ERT-Th-Cre
αTH
αACTN
*
αDAT
Males
L
Females
Males
control
Gch1ff;ERT-Th-Cre
****
**
TH protein/Actin
Months after Tamoxifen
M
Females
Males
**
NS
DAT protein/Actin
Months after Tamoxifen

FIG.S7

**Supplemental Figure 7. BH4 deficiency leads to reduced TH levels.**

**A,** Representative immunofluorescence of TH-staining of DAergic neurons in the striatum and SNpc (left) and quantification of SNpc/VTA neuronal TH intensity (right) in midbrain from 3-week old control and *Gch1$^{flox/flox}$;Dat-Cre* mice. Scale bar, 100µm. Data are shown as means ± s.e.m. Intensities of >100 TH$^{+}$ neurons pooled from n=5 animals of each genotype. ****P < 0.0001 (Student's t-test). **B-E,** Representative image of TH immunohistochemistry in the SNpc of 3-week old control and *Gch1$^{flox/flox}$;Dat-Cre* mice **(B)** and quantification of TH$^{+}$ cells through 20µm consecutive sections through the SNpc **(C)** and the ventral tegmental area (VTA) **(D)** and total TH$^{+}$ cell counts in the SNpc, VTA and SNpc+VTA **(E)** from control and *Gch1$^{flox/flox}$;Dat-Cre* mice. Data are shown as means ± s.e.m. Individual mice for each genotype are shown. *P < 0.05; ***P<0.001; NS, not significant (Student's t-test with multiple comparisons). **F,** Western blot analysis of TH, DAT and DDC in the ventral midbrain of control and *Gch1$^{flox/flox}$;Dat-Cre* mice at 3 days (top panel), and 21 days (bottom panel) after birth. N=3/4 per genotype per time point. Actin is used as a loading control. **G,H,** Representative immunofluorescence of DAT (main) and TH (inset) neurons in the SNpc **(G)** and quantification of antibody intensities **(H)** of 3-week old control and *Gch1$^{flox/flox}$;Dat-Cre* mice. Data are shown as means ± s.e.m. Individual cell intensities pooled from n=5 mice (H, top) and from individual mice for each genotype are shown (H, bottom). ***P<0.001; ****P < 0.0001; NS, not significant (Student's t-test with multiple comparisons). **I,** top**,** Western blot analysis of TH and DAT in striatal tissue from control and *Gch1$^{flox/flox}$;Dat-Cre* mice at 21 days after birth. N=2/6 per genotype. Actin is used as a loading control. **I,** bottom**,** Representative immunofluorescence of DAT staining in the striatum of 3-week old control and *Gch1$^{flox/flox}$;Dat-Cre* mice. **J,K,** Western blot of TH and DAT in the striatal tissue of female **(J)** and male **(K)** control and *Gch1$^{flox/flox}$;ERT-Th-Cre* mice at 6 and 12 months after tamoxifen administration. Actin is used as a loading control. Asterix indicates non- specific band which is recognized by the anti-TH antibody in these preparations. **L,M,** Normalized quantification of TH **(L)** and DAT **(M)** proteins in the striatal tissue of female and male control and *Gch1$^{flox/flox}$;ERT-Th-Cre* mice at 6 and 12 months after tamoxifen administration. Data are shown as means ± s.e.m. Individual mice for each genotype are shown. **P < 0.01; ****P < 0.0001; NS, not significant (Multiple t-test).

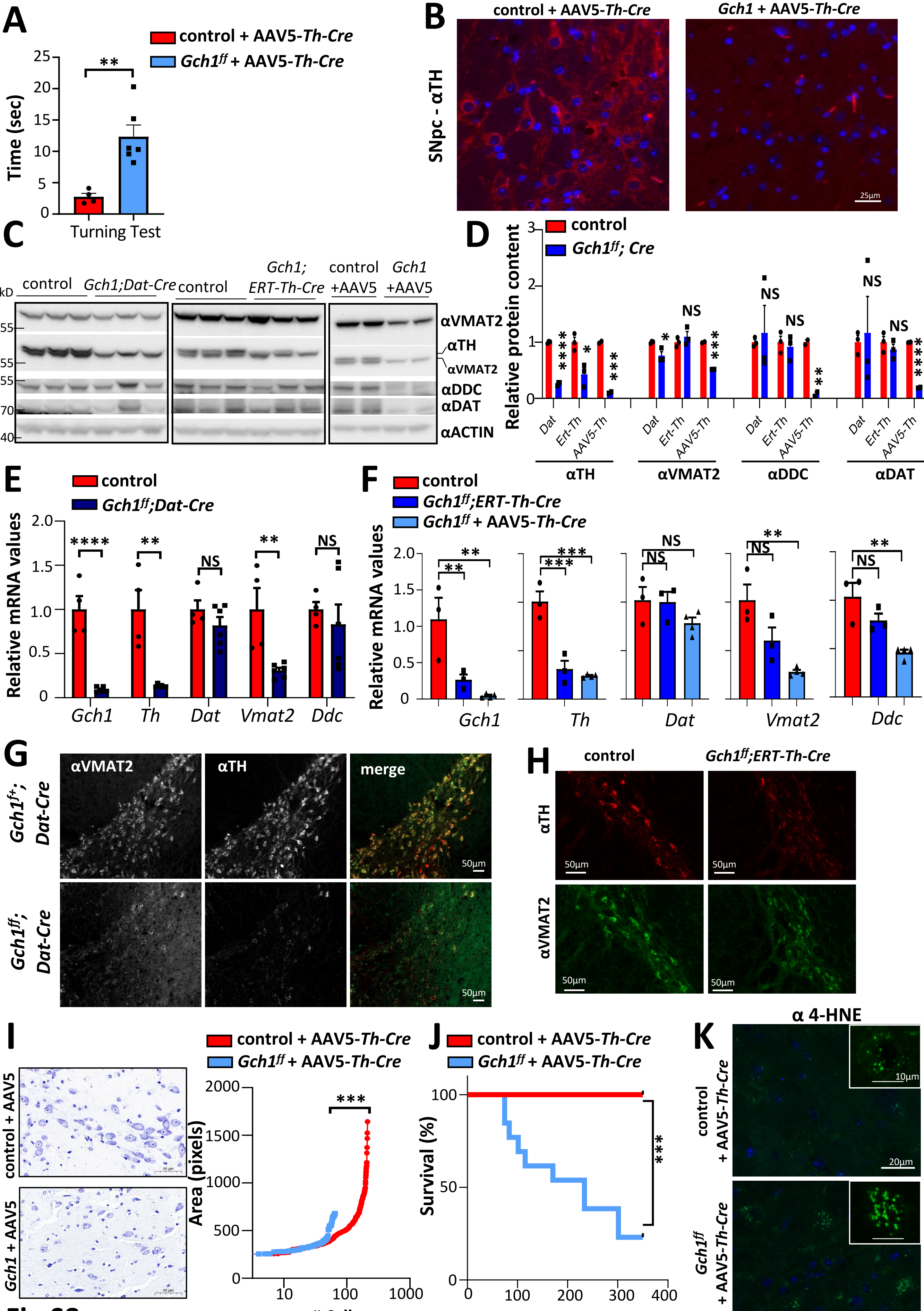

A
control + AAV5-Th-Cre
Gch1ff + AAV5-Th-Cre
Time (sec)
Turning Test
B
control + AAV5-Th-Cre
Gch1 + AAV5-Th-Cre
SNpc - αTH
25μm
C
control
Gch1;Dat-Cre
control
Gch1; ERT-Th-Cre
control +AAV5
Gch1 +AAV5
kD
αVMAT2
αTH
αVMAT2
αDDC
αDAT
αACTIN
D
control
Gch1ff; Cre
Relative protein content
Dat
Ert-Th
AAV5-Th
αTH
αVMAT2
αDDC
αDAT
NS
E
control
Gch1ff;Dat-Cre
Relative mRNA values
Gch1
Th
Dat
Vmat2
Ddc
NS
F
control
Gch1ff;ERT-Th-Cre
Gch1ff + AAV5-Th-Cre
Relative mRNA values
Gch1
Th
Dat
Vmat2
Ddc
NS
G
αVMAT2
αTH
merge
Gch1f/+; Dat-Cre
Gch1ff; Dat-Cre
50μm
H
control
Gch1ff;ERT-Th-Cre
αTH
αVMAT2
50μm
I
control + AAV5
Gch1 + AAV5
control + AAV5-Th-Cre
Gch1ff + AAV5-Th-Cre
Area (pixels)
### Cells
J
control + AAV5-Th-Cre
Gch1ff + AAV5-Th-Cre
Survival (%)
Days after AAV5-Th-Cre
K
α 4-HNE
control + AAV5-Th-Cre
Gch1ff + AAV5-Th-Cre
10μm
20μm


Fig.S8

**Supplemental Figure 8. Extent and effects of BH4 deficiency on DAergic neuronal markers.**

**A,** Turning test evaluation of control and *Gch1*$^{flox/flox}$ mice 3 months after AAV5-*Th-Cre* administration. Data are means ± s.e.m. Individual mice for each genotype are shown. **P < 0.01 (Student's t-test). **B**, Representative images of anti-TH immunofluorescence in the SNpc of control and *Gch1*$^{flox/flox}$ mice 4 months after AAV5-*Th-Cre* injection. Scale bar, 25µm. **C,D,** Western blot for TH, VMAT2, DDC and DAT in striatal tissue from control, *Gch1*$^{flox/flox}$*;Dat-Cre, Gch1*$^{flox/flox}$*;ERT-Th-Cre* and *Gch1*$^{flox/flox}$ mice injected with AAV5-*Th-Cre* mice. Actin is used as a loading control (**C**) and quantification of relative expression (**D**). Data are shown as means ± s.e.m. Individual mice for each genotype are shown. *P<0.05; **P<0.01; ***P<0.001; ****P<0.0001; NS, not significant (Multiple t-test comparison to relative controls). **E,F,** RT-qPCR of *Gch1*, *Th*, *Dat, Ddc* and *Vmat2* mRNA from the ventral midbrain of control, *Gch1*$^{flox/flox}$*;Dat-Cre, Gch1*$^{flox/flox}$*;ERT-Th-Cre* and *Gch1*$^{flox/flox}$ + AAV5-*Th-Cre* mice. Data are shown as means ± s.e.m. Individual mice for each genotype are shown. **P<0.01; ***P<0.001; ****P<0.0001; NS, not significant (One-way ANOVA with Dunnett's multiple comparison test). **G**, Representative images of anti-VMAT2 and anti-TH immunofluorescence in the SNpc of control and *Gch1*$^{flox/flox}$*;Dat-Cre* mice. **H**, Representative images of anti-TH and anti-VMAT2 immunostaining in the SNpc of control and *Gch1*$^{flox/flox}$*;ERT-Th-Cre* mice 28 weeks post tamoxifen administration. **I,** Representative images of cresyl violet staining (left) and quantification of number and pixel area of cells (right) in the SNpc of control and *Gch1*$^{flox/flox}$ mice 4 months after AAV5-*Th-Cre* injection. Scale bar, 50µm. Data for individual cells are shown and pooled from n=3 mice for each genotype. ***P<0.001 (Student's t-test). **J,** Kaplan-Meier survival curves of control (n=10) and *Gch1*$^{flox/flox}$ (n=10) mice after AAV5-*Th-Cre* injection as indicated. ***P < 0.001 (Log-rank (Mandel-Cox) test). **K,** Representative 4-HNE immunohistochemistry as a marker for lipid peroxidation (and ferroptosis) in the SNpc of control and *Gch1*$^{flox/flox}$ mice 1 month after AAV5-*Th-Cre* injection. 4-HNE, 4-Hydroxynonenal. Scale bar, 25µm. Inset shows a more magnified representative cell from the SNpc region in control and *Gch1*$^{flox/flox}$ mice 1 month after AAV5-*Th-Cre* injection. Scale bar, 10 µm.

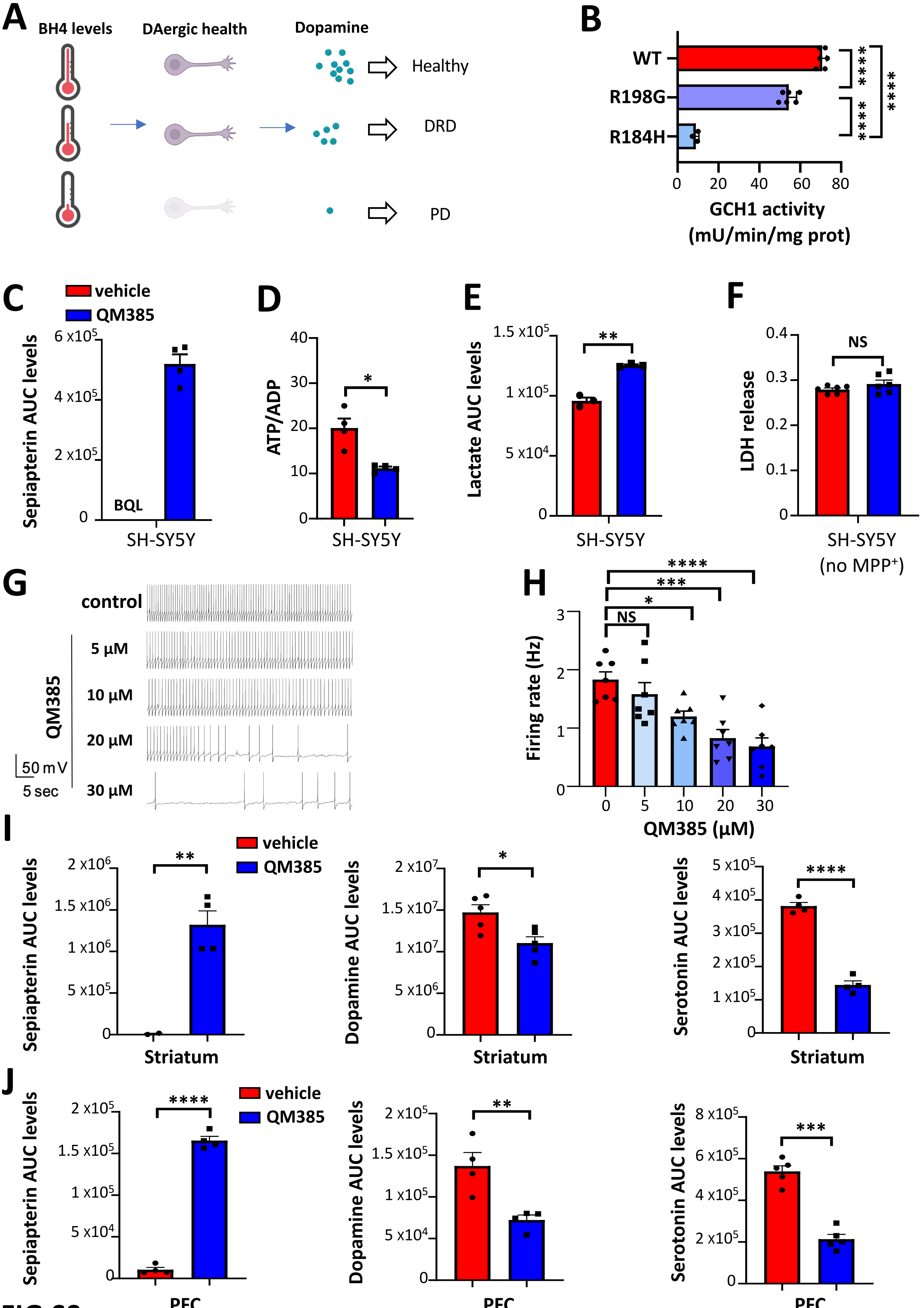
A
BH4 levels
DAergic health
Dopamine
Healthy
DRD
PD
B
WT
R198G
R184H
****
****
****
0 20 40 60 80
GCH1 activity
(mU/min/mg prot)
C
vehicle
QM385
Sepiapterin AUC levels
BQL
SH-SY5Y
D
ATP/ADP
SH-SY5Y
E
Lactate AUC levels
SH-SY5Y
F
NS
LDH release
SH-SY5Y
(no MPP⁺)
G
control
5 µM
10 µM
20 µM
30 µM
QM385
50 mV
5 sec
H
Firing rate (Hz)
QM385 (µM)
I
Sepiapterin AUC levels
Dopamine AUC levels
Serotonin AUC levels
Striatum
J
PFC

FIG.S9

**Supplemental Figure 9. Metabolic profiling of QM385-treated SH-SY5Y cells and mice.**

**A,** Schematic depicting the hypothesis that brain levels of BH4 may dictate the severity of diseases associated with nigrostriatal DAergic dysfunction. **B,** Activity of purified wild type GCH1 and disease-associated GCH1 mutants (R198 (DRD) and R184 (PD)). Data are shown as means ± s.e.m. Individual samples are shown. ****P < 0.0001 (One-way ANOVA with Tukey's multiple comparison test). **C-E,** Metabolic profiling of sepiapterin levels (**C**), ATP/ADP ratios (**D**) and lactate levels (**E**) in SH-SY5Y cells treated with vehicle (DMSO) and QM385 (5µM) for 16 hours. BQL, below quantifiable levels. Data are shown as means ± s.e.m. Individual samples are shown. *P < 0.05; **P < 0.01; NS, not significant (Student's t-test). AUC, under the curve (see methods for quantification of metabolites). **F,** Viability as determined by lactate dehydrogenase (LDH) release from SH-SY5Y cells treated with vehicle (DMSO) and QM385 (5µM) for 48 hours. Data are shown as means ± s.e.m. Individual samples are shown. NS, not significant (Student's t-test). **G,H,** Representative traces of spontaneous pace-making action potentials (APs) from DAergic neurons in mouse SNpc under dose response of QM385 treatments (**G**) and quantification of the firing rates under each condition (**H**). Data are shown as means ± s.e.m. Individual brain slice recordings are shown. *P < 0.05; ***P < 0.001; ****P < 0.0001; NS, not significant (One-way ANOVA with Dunnett's multiple comparison test). **I,J,** Measurements of sepiapterin, dopamine and serotonin levels in the striatum **(I)** and prefrontal cortex (PFC) **(J)** of mice treated for 5 days with QM385, 5mg/kg, or vehicle i/p, twice daily. Tissues were extracted 6 hours after the final treatment. Data are shown as means ± s.e.m. Individual mice are shown. *P < 0.05; **P < 0.01; ***P < 0.001; ****P < 0.0001 (Student's t-test). AUC, under the curve (see methods for quantification of metabolites).

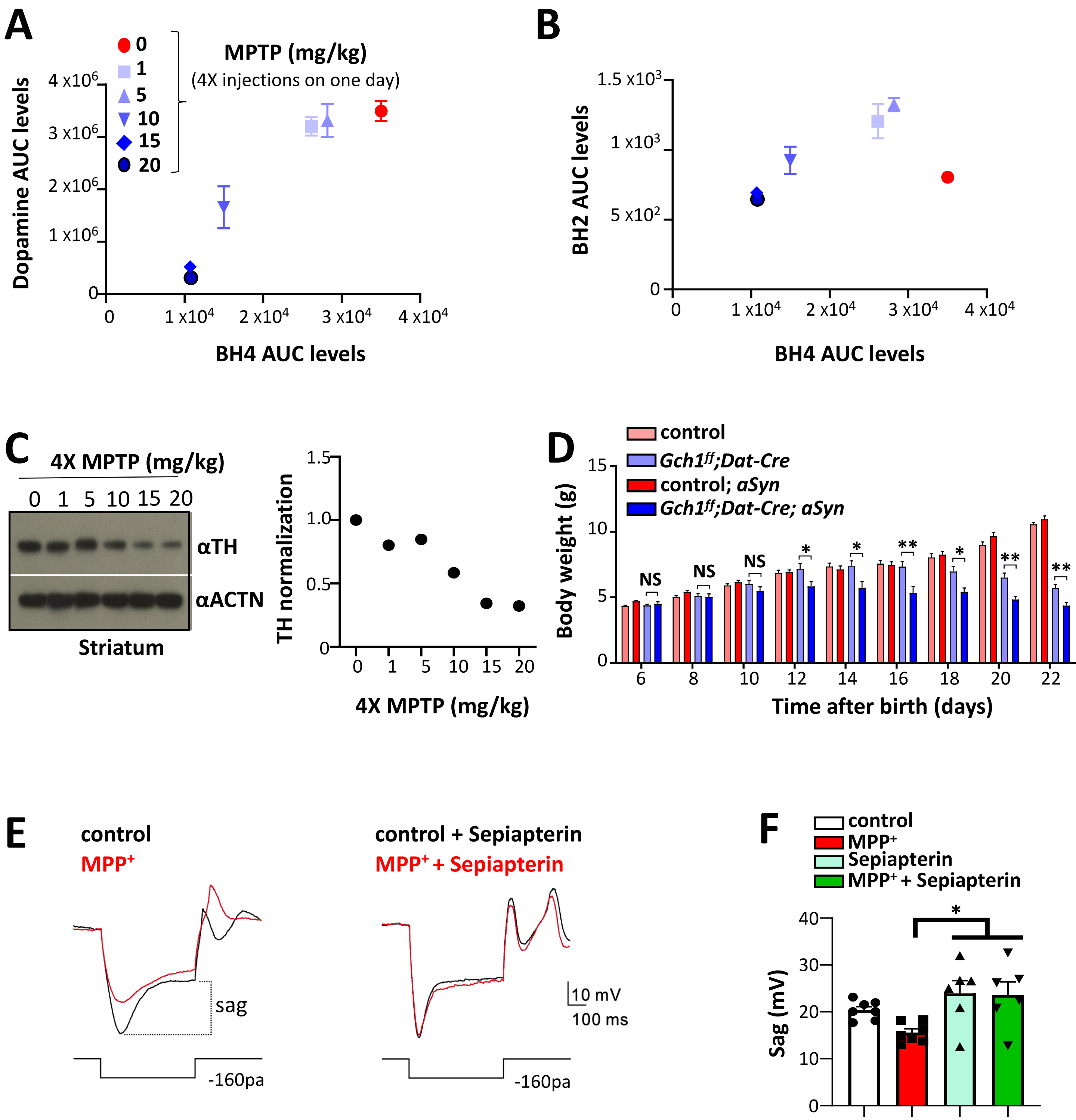

A
MPTP (mg/kg)
(4X injections on one day)
0
1
5
10
15
20
Dopamine AUC levels
BH4 AUC levels
B
BH2 AUC levels
BH4 AUC levels
C
4X MPTP (mg/kg)
0 1 5 10 15 20
αTH
αACTN
Striatum
TH normalization
4X MPTP (mg/kg)
D
control
Gch1ff;Dat-Cre
control; aSyn
Gch1ff;Dat-Cre; aSyn
Body weight (g)
NS
Time after birth (days)
E
control
MPP+
control + Sepiapterin
MPP+ + Sepiapterin
sag
10 mV
100 ms
-160pa
F
control
MPP+
Sepiapterin
MPP+ + Sepiapterin
Sag (mV)


FIG.S10

**Supplemental Figure 10. The BH4 pathway regulates the pacemaking activity of mouse midbrain slices.**

**A-C,** Effect of in vivo administration of MPTP (indicated doses (mg/kg), i.p.) on striatal dopamine, BH4 and BH2 levels (**A,B**) as well as on TH protein levels. Actin is used as a loading control (**C**).**D,** Body weight measurements after birth of control, *Gch1;Dat-Cre*, control; *aSyn*, and *Gch1$^{flox/flox}$;Dat-Cre; aSyn* mice. Data are shown as mean ± s.e.m. *P < 0.05; **P < 0.01; NS, not significant (Student's t-test with multiple comparisons between *Gch1$^{flox/flox}$;Dat-Cre* and *Gch1$^{flox/flox}$;Dat-Cre; aSyn* mice). **E,** Representative traces showing sag voltage in response to current injection (-160 pA) recorded from mouse SNPC DAergic neurons. The sag is indicated. **F,** Quantification of the sag voltage under each condition (with and without 10 μM sepiapterin, 20 μM $MPP^+$). Data are shown as means ± s.e.m. Individual slice recordings for each condition are shown. *P < 0.05 (One-way ANOVA with Tukey's multiple comparisons).

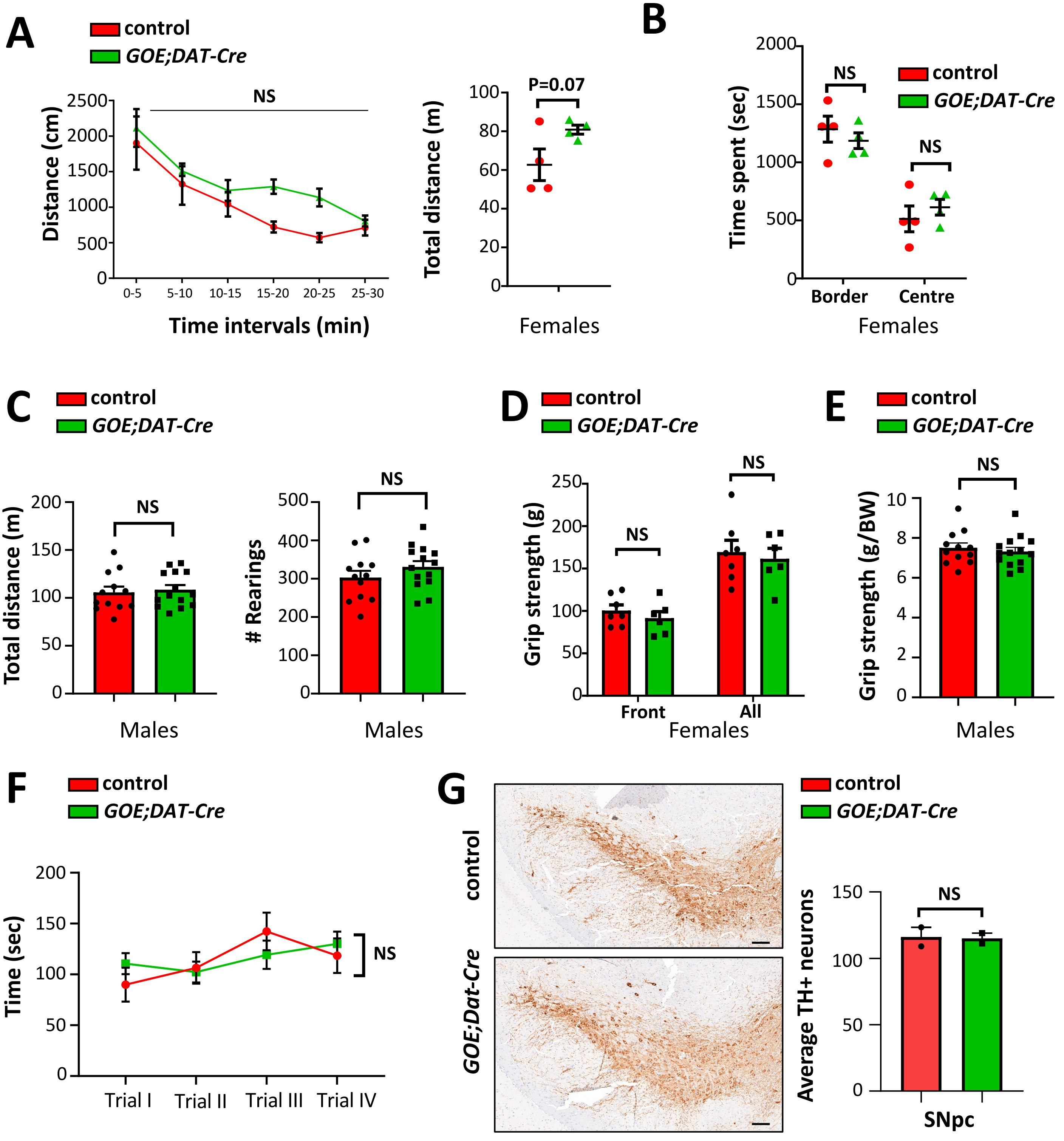

A
control
GOE;DAT-Cre
NS
Distance (cm)
Time intervals (min)
0-5
5-10
10-15
15-20
20-25
25-30
P=0.07
Total distance (m)
Females
B
NS
control
GOE;DAT-Cre
Time spent (sec)
Border
Centre
Females
C
control
GOE;DAT-Cre
Total distance (m)
Males
### Rearings
Males
D
control
GOE;DAT-Cre
Grip strength (g)
Front
All
Females
E
control
GOE;DAT-Cre
Grip strength (g/BW)
Males
F
control
GOE;DAT-Cre
Time (sec)
Trial I
Trial II
Trial III
Trial IV
NS
G
control
GOE;Dat-Cre
Average TH+ neurons
SNpc


Fig.S11

**Supplemental Figure 11. *Gch1* overexpression in DAergic neurons does not affect baseline motor behavior nor DAergic neuronal numbers.**

**A,B,** Quantification of interval distance (**A,** left graph) and total distance (**A,** right graph) travelled as well as total time spent at the border/center (**B**) of female control and *GOE;DAT-Cre* mice in open field testing during a 30-minute observational period. Data are shown as means ± s.e.m. NS, not significant (multiple t-test). **C**, Total distance travelled (left) and number of rearings (right) of male control and *GOE;DAT-Cre* mice in open field testing during a 30-minute observational period. Data are shown as means ± s.e.m. Individual mice for each genotype are shown. NS, not significant (Student's t-test). **D,E,** Grip strength of female (**D**) and male (**E**) control and *GOE;DAT-Cre* mice. Data are shown as means ± s.e.m. Individual mice for each genotype are shown. NS, not significant (Student's t-test and multiple t-test). **F,** Accelerated (4-40rpm) rotarod testing over various trials of control and *GOE;DAT-Cre* mice. Data are shown as means ± s.e.m. NS, not significant (Two-way ANOVA). **G,** Representative, and quantification of, TH staining in the SNpc of controls and *GOE;DAT-Cre* mice. NS, not significant (Student's t-test).

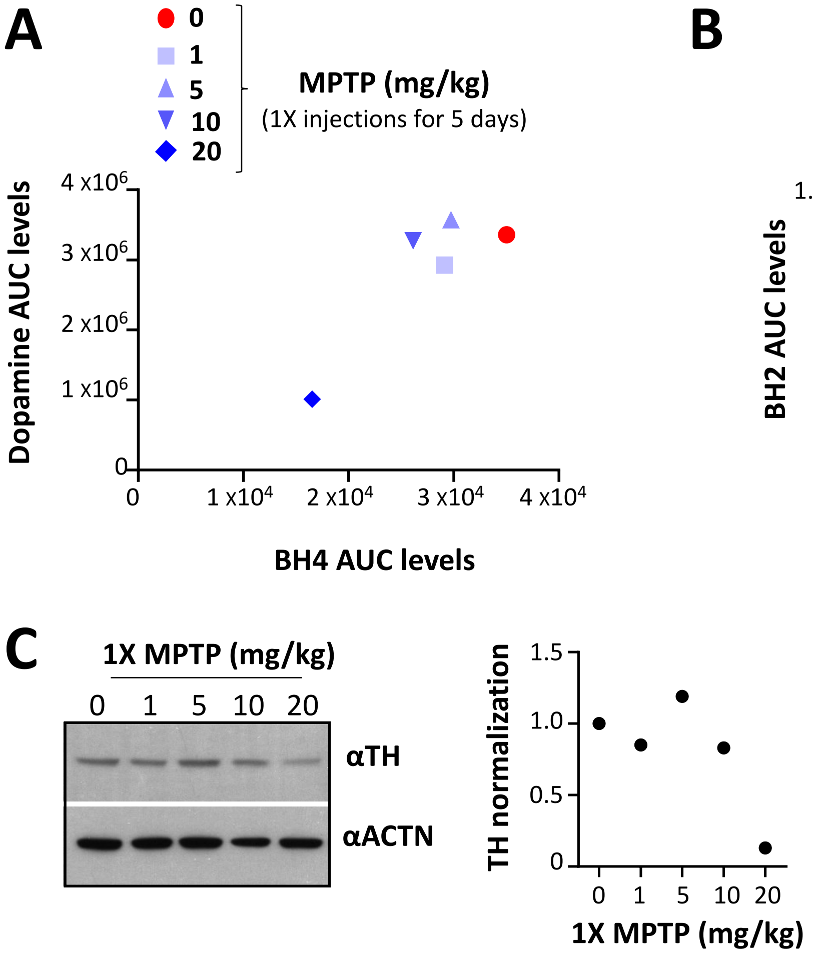
A
0
1
5
10
20
MPTP (mg/kg)
(1X injections for 5 days)
Dopamine AUC levels
4 x10^6
3 x10^6
2 x10^6
1 x10^6
0
1 x10^4
2 x10^4
3 x10^4
4 x10^4
BH4 AUC levels
B
BH2 AUC levels
1.5 x10^3
1 x10^3
5 x10^2
0
1 x10^4
2 x10^4
3 x10^4
4 x10^4
BH4 AUC levels
C
1X MPTP (mg/kg)
0 1 5 10 20
αTH
αACTN
TH normalization
1.5
1.0
0.5
0
0 1 5 10 20
1X MPTP (mg/kg)

Fig.S12

**Supplemental Figure 12. Response of varying doses of MPTP on striatal dopamine, BH4, BH2 and TH levels.**

**A,B,** Striatal BH4, dopamine (**A**) and BH2 (**B**) levels in wild type mice one week post MPTP treatment at the indicated doses. AUC, under the curve (see methods for quantification of metabolites). **C,** Western blot (left) and quantification (right) of tyrosine hydroxylase (TH) protein levels in the striatum of wild type mice one week post MPTP treatment at the indicated doses. Actin was used as a loading control.

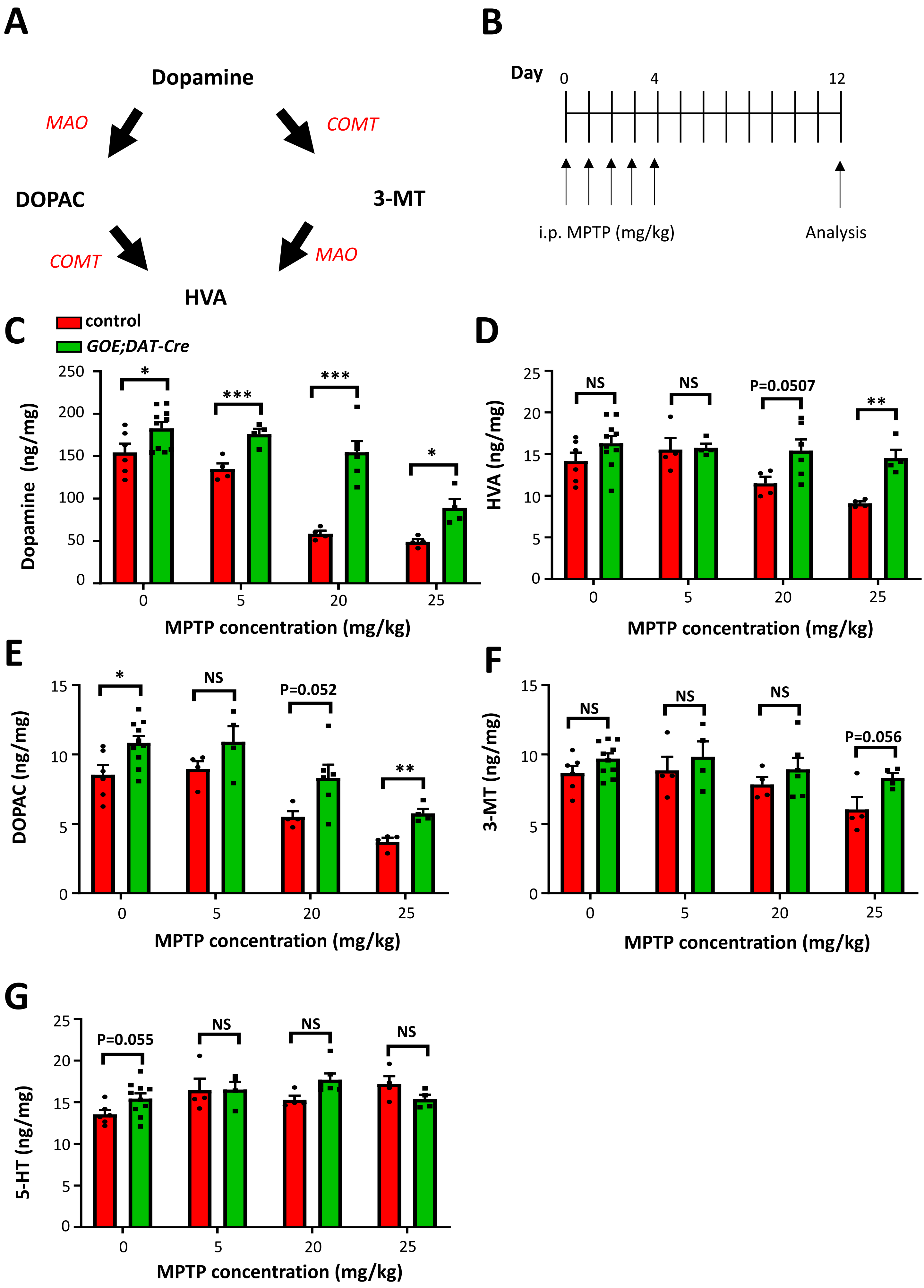
A
Dopamine
MAO
COMT
DOPAC
3-MT
COMT
MAO
HVA
B
Day
0
4
12
i.p. MPTP (mg/kg)
Analysis
C
control
GOE;DAT-Cre
Dopamine (ng/mg)
MPTP concentration (mg/kg)
D
HVA (ng/mg)
NS
NS
P=0.0507
MPTP concentration (mg/kg)
E
DOPAC (ng/mg)
NS
P=0.052
MPTP concentration (mg/kg)
F
3-MT (ng/mg)
NS
NS
NS
P=0.056
MPTP concentration (mg/kg)
G
5-HT (ng/mg)
P=0.055
NS
NS
NS
MPTP concentration (mg/kg)

Fig.S13

**Supplemental Figure 13. Enhanced BH4 in DAergic neurons ameliorates MPTP-mediated dopamine decline in mice.**

**A,** Schematic depicting dopamine metabolism. MAO, monoamine oxidase; COMT, Catechol-O-methyltransferase; DOPAC, 3,4-Dihydroxyphenylacetic acid; HVA, homovanillic acid; 3-MT, 3-methoxytyramine. **B,** Schematic depicting administration of MPTP (i.p.) *in vivo* each day for 5 contiguous days. **C-G,** Dopamine (**C**), and its metabolites HVA (**D**), DOPAC (**E**) and 3-MT (**F**) as well as serotonin (5-HT) (**G**) levels in the striatum of control and *GOE;DAT-Cre* mice after varying doses of MPTP treatment as outlined in (B). Data are means ± s.e.m. Individual mice for each genotype are shown. *$P < 0.05$; **$P < 0.01$; ***$P < 0.001$; NS, not significant (t-test with multiple comparisons).

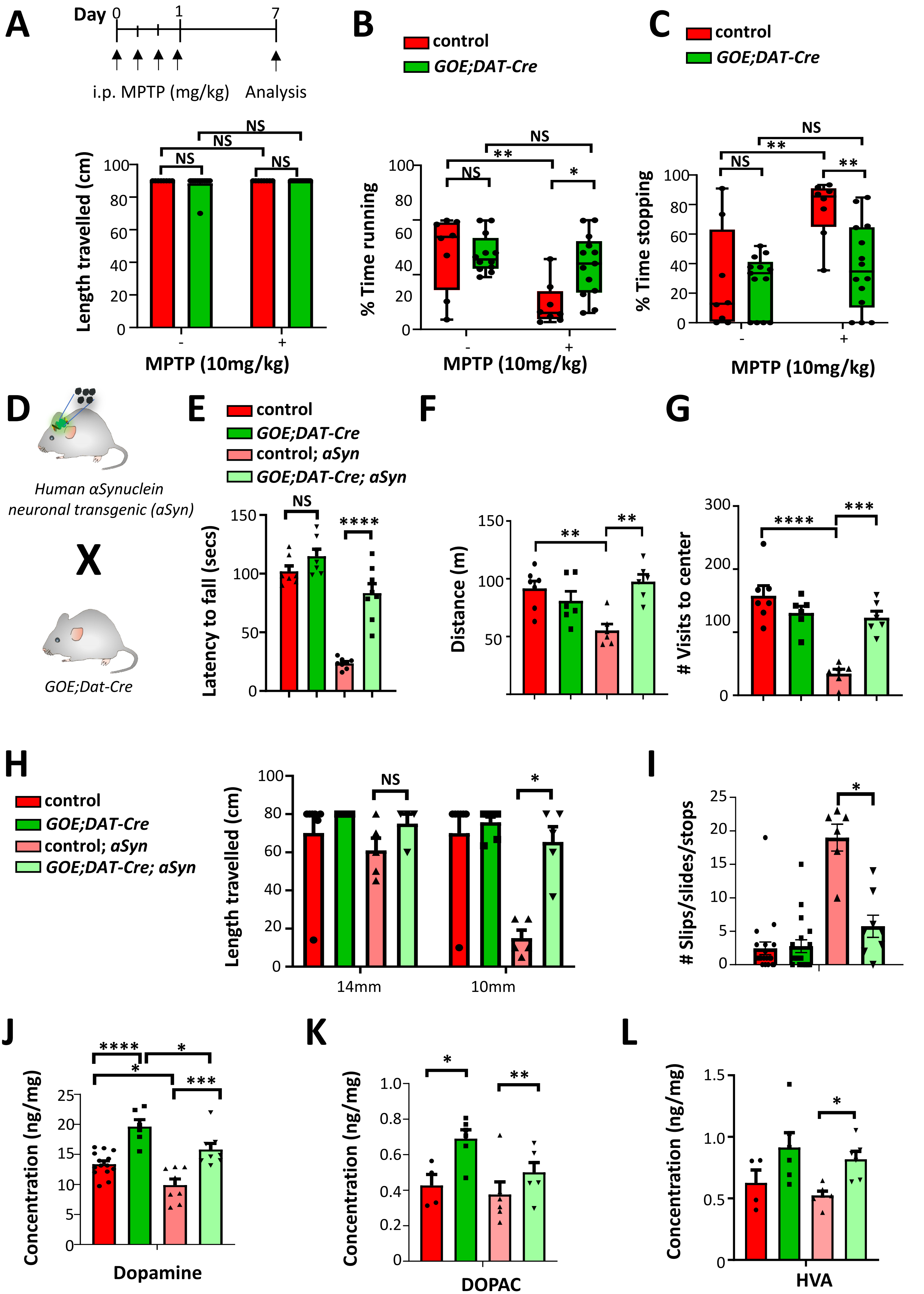
A
Day 0 1 7
i.p. MPTP (mg/kg)
Analysis
Length travelled (cm)
MPTP (10mg/kg)
B
control
GOE;DAT-Cre
% Time running
C
% Time stopping
D
Human αSynuclein neuronal transgenic (aSyn)
X
GOE;Dat-Cre
E
control; aSyn
GOE;DAT-Cre; aSyn
Latency to fall (secs)
F
Distance (m)
G
### Visits to center
H
Length travelled (cm)
14mm
10mm
I
### Slips/slides/stops
J
Concentration (ng/mg)
Dopamine
K
DOPAC
L
HVA

Fig.S14

**Supplemental Figure 14. Enhanced BH4 in DAergic neurons ameliorates motor decline caused by alpha-synuclein overexpression.**

**A-C,** MPTP injection scheme and beam walk behavioral testing of control and *GOE;DAT-Cre* mice, 14 days after MPTP treatment from (Figure 2B) in which distance travelled **(A)** and proportion of time running **(B)** as well as stopping **(C)** across a 20mm-diameter beam was recorded. **D**, Schematic depicting the breeding strategy for incorporating the human alpha-synuclein (aSyn) overexpression transgene into *GOE;Dat-Cre* mice. **E**, Rotarod analysis with constant (4km/hr) speed of control, *GOE;Dat-Cre*, control; *aSyn*, and *GOE;Dat-Cre;aSyn* mice. Data are means ± s.e.m. Individual mice for each genotype are shown. ****P<0.0001; NS, not significant (One-way ANOVA with Tukey's multiple comparisons). **F,G,** Open field behavioral testing in which total distance (**F**) and visits to center (**G**) were recorded in the 30-minute observational period of control, *GOE;Dat-Cre*, control; *aSyn*, and *GOE;Dat-Cre;aSyn* mice. Data are means ± s.e.m. Individual mice for each genotype are shown. **P<0.01; ***P<0.001; ****P<0.0001; NS, not significant (One-way ANOVA with Tukey's multiple comparisons). **H,I,** Beam walk behavioral testing of 8-month-old control, *GOE;Dat-Cre*, control; *aSyn*, and *GOE;Dat-Cre; aSyn* mice. Distance travelled crossing along beams of different diameters (**H**) and number of stops, slips and slides during crossing along beams of different diameters (**I**) were measured. Data are means ± s.e.m. Individual mice for each genotype are shown. *P < 0.05; NS, not significant (Student's t-test with multiple comparisons). **J-L,** Dopamine (**J**), DOPAC **(K)** and HVA (**L**) levels in the midbrain of control, *GOE;Dat-Cre*, control; *aSyn*, and *GOE;Dat-Cre;aSyn* mice. Data are shown as means ± s.e.m. Individual mice for each genotype are shown. *P < 0.05; ***P < 0.001; ****P < 0.0001; NS, not significant (One-way ANOVA with Tukey's multiple comparisons).

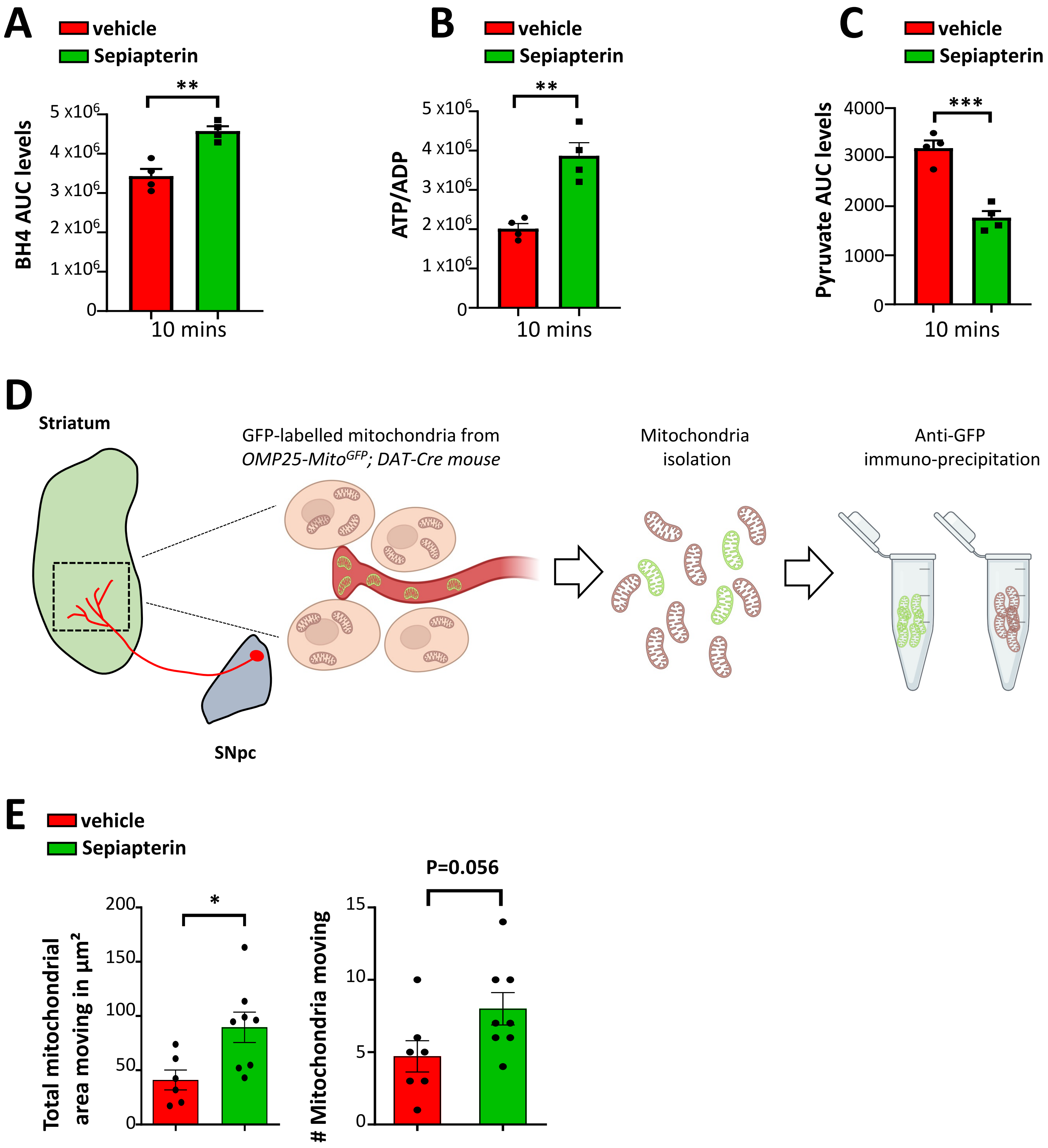
A
vehicle
Sepiapterin
**
BH4 AUC levels
5 x10^6
4 x10^6
3 x10^6
2 x10^6
1 x10^6
0
10 mins
B
vehicle
Sepiapterin
**
ATP/ADP
5 x10^6
4 x10^6
3 x10^6
2 x10^6
1 x10^6
0
10 mins
C
vehicle
Sepiapterin
***
Pyruvate AUC levels
4000
3000
2000
1000
0
10 mins
D
Striatum
SNpc
GFP-labelled mitochondria from
OMP25-MitoGFP; DAT-Cre mouse
Mitochondria
isolation
Anti-GFP
immuno-precipitation
E
vehicle
Sepiapterin
*
Total mitochondrial
area moving in µm²
200
150
100
50
0
P=0.056
### Mitochondria moving
15
10
5
0

**Fig.S15**

**Supplemental Figure 15. BH4 improves mitochondrial health and function.**

**A-C,** BH4 (**A**), ATP/ADP (**B**) and pyruvate (**C**) levels in SH-SY5Y cells treated for ten minutes with DMSO (vehicle) and sepiapterin (10µM). Data are shown as means ± s.e.m. Individual samples are shown. BQL, below quantifiable levels. AUC, under the curve (see methods for quantification of metabolites). **P < 0.01; ***P < 0.001 (Student's t-test). **D,** Schematic depicting the isolation of DAergic-specific mitochondria localized in projections from the SNpc in the striatum using the (*R26-Lox-STOP-Lox – 3XHA-eGFP-OMP25Mito$^{HA}$; referred to as OMP25-Mito$^{GFP}$*) reporter mouse. Total mitochondria are isolated from the striatal tissue and using anti-GFP microbeads, the mitochondria from SNpc DAergic neurons can be selectively isolated. Non-DAergic originated mitochondria from the striatum are also isolated. **E**, Quantitation of total mitochondrial area (left panel) and mitochondrial numbers (right panel) transported retrograde from axons to soma after 4 days of treatment with vehicle, and sepiapterin. Data are shown as means ± s.e.m. Individual imaged axons are shown *P < 0.05 (Student's t-test).

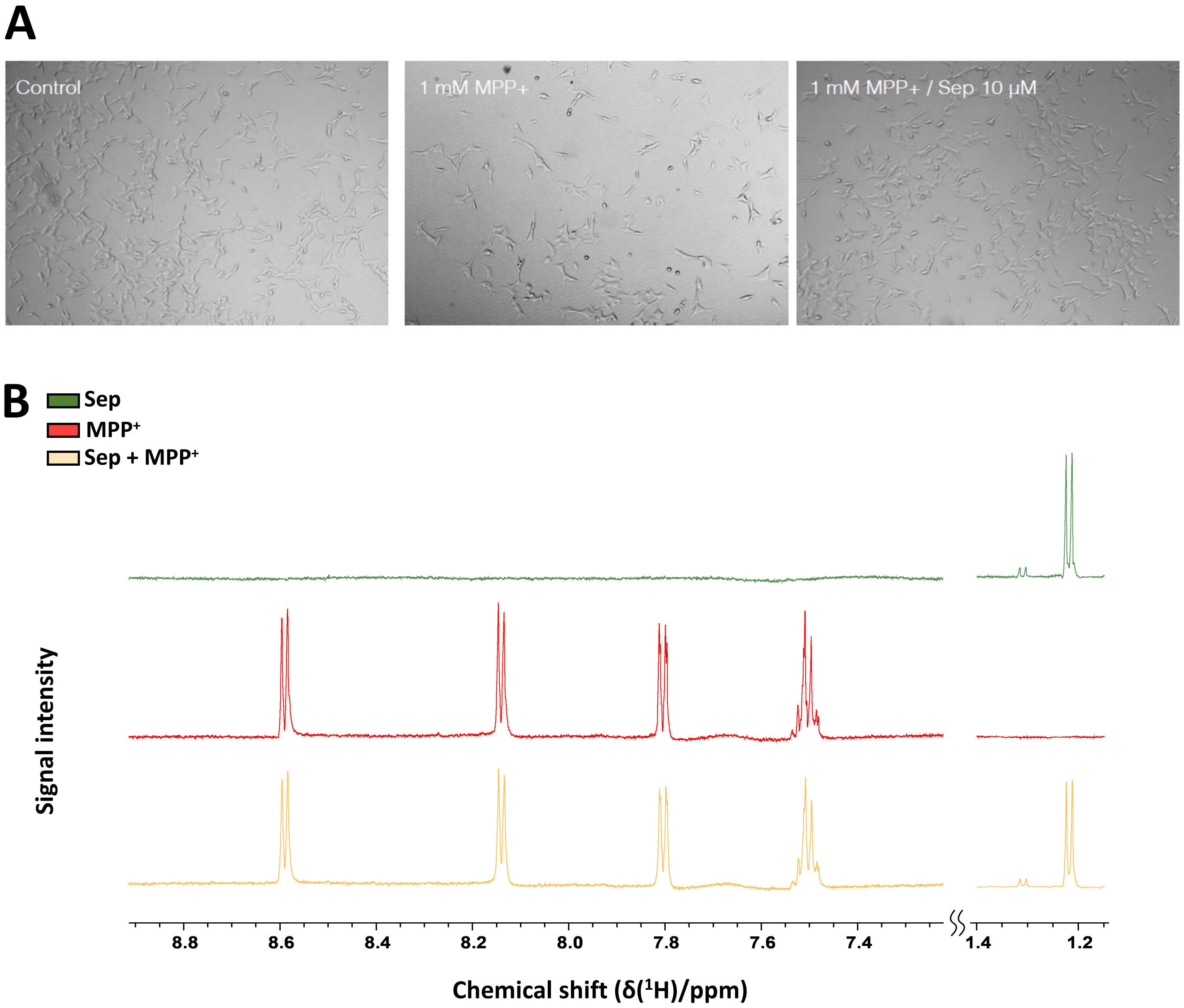
A
Control
1 mM MPP+
1 mM MPP+ / Sep 10 µM
B
Sep
MPP+
Sep + MPP+
Signal intensity
8.8
8.6
8.4
8.2
8.0
7.8
7.6
7.4
1.4
1.2
Chemical shift (δ(1H)/ppm)

Fig.S16

**Supplemental Figure 16. Increased BH4 protects against $MPP^+$-induced toxicity of DAergic neurons *in vitro.***

**A,** Representative brightfield images of SH-SY5Y cells untreated (control) or treated with $MPP^+$ (1mM) as well as $MPP^+$ and sepiapterin (Sep) for 48 hours. **B,** One-dimensional proton nuclear magnetic resonance (1D $H^1$ NMR) spectra of sepiapterin, $MPP^+$ and sepiapterin plus $MPP^+$ with the horizontal axes showing the chemical shifts of the compounds (units of ppm) and the y-axes depicting the intensity which is proportional to the concentration. Note that the NMR signals do not change upon mixing the compounds.

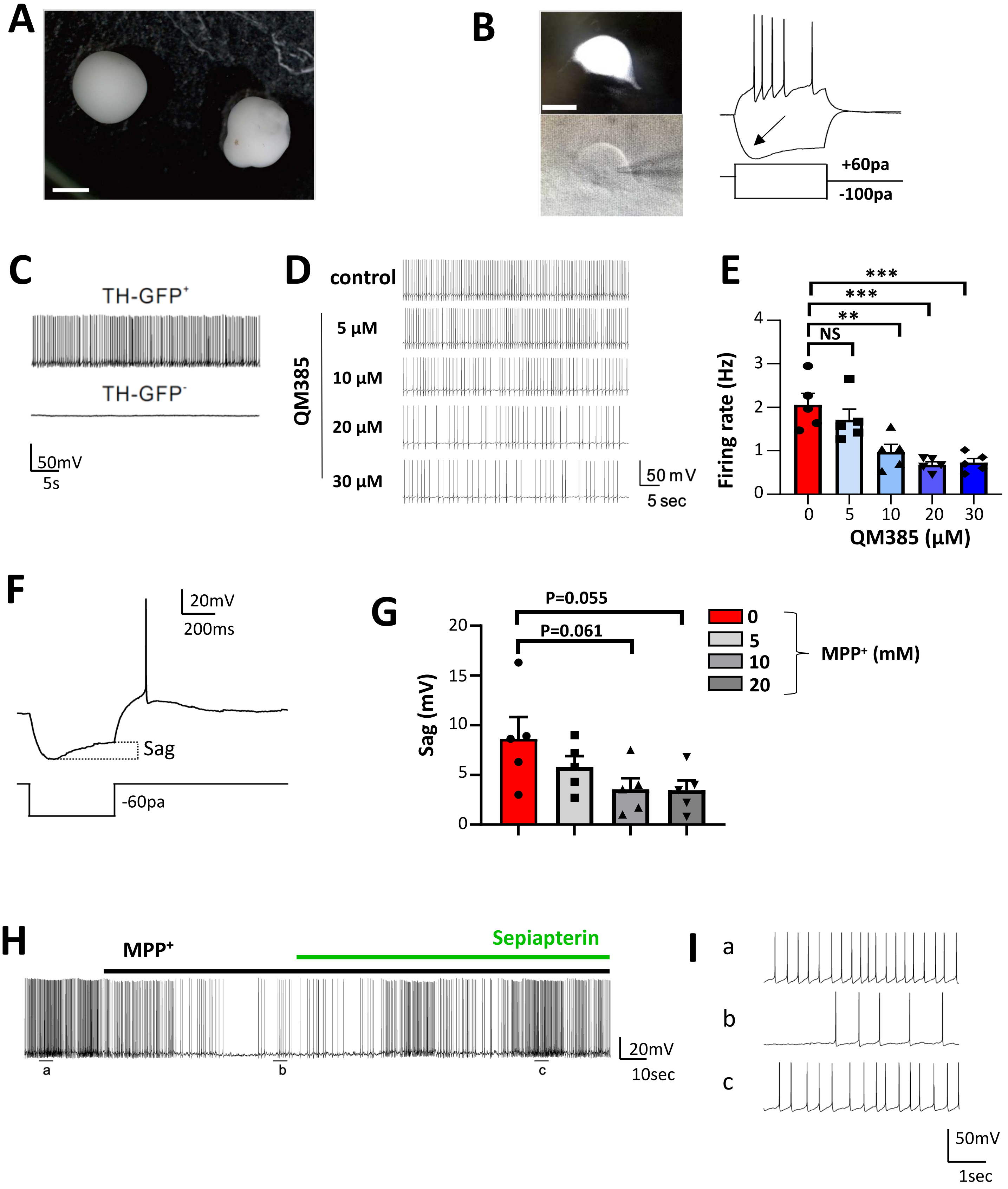
A
B
+60pa
-100pa
C
TH-GFP+
TH-GFP-
50mV
5s
D
control
QM385
5 μM
10 μM
20 μM
30 μM
50 mV
5 sec
E
Firing rate (Hz)
QM385 (μM)
NS
**
***
***
F
20mV
200ms
Sag
-60pa
G
Sag (mV)
P=0.055
P=0.061
MPP+ (mM)
0
5
10
20
H
MPP+
Sepiapterin
20mV
10sec
a
b
c
I
a
b
c
50mV
1sec

Fig.S17

**Supplemental Figure 17. The BH4 pathway regulates the pacemaking activity of human DAergic neurons.**

**A,** Brightfield images of human midbrain-like organoids (hMLOs). **B,** Differential interface contrast (DIC) images illustrate the typical morphology of recorded TH-GFP$^+$ neurons within hMLOs (left). Scale bars, 10 μm. Representative traces of multiple APs recorded from a TH-GFP$^+$ neuron in hMLOs at day 90 (right). Note the archetypal sag (arrow). **C,** Representative traces of spontaneous pace-making APs recorded from TH-GFP$^+$ neurons (top) and TH-GFP$^-$ neurons (bottom) within the same hMLO. **D**,**E,** Representative traces of spontaneous APs recorded from TH-GFP$^+$ neurons in hMLOs under different concentrations of QM385 (**D**) and quantification of frequencies of spontaneous APs recorded from TH-GFP$^+$ neurons within hMLOs under each condition (**E**). Error bars represent mean ± SEM. Data are shown as means ± s.e.m. Individual recordings for each condition are shown. **P < 0.01; ***P < 0.001; NS, not significant (One-way ANOVA with Dunnett's multiple comparisons). **F,** Representative traces of rebound AP recorded from TH-GFP$^+$ neurons within human MLOs. The rebound sag is indicated. G**,** Quantification of the effect of MPP$^+$ on sag voltage in a dose-dependent manner. Error bars represent mean ± SEM. Data are shown as means ± s.e.m. Individual neuronal recordings for each condition are shown. P values are indicated (One-way ANOVA with Dunnett's multiple comparisons). **H,** Representative traces of spontaneous APs recorded from TH-GFP$^+$ neuron within hMLOs under each condition (10 μM sepiapterin, 20 μM MPP$^+$). **I,** Magnified traces corresponding to each time frame marked in H (a-c). Applications of MPP$^+$ (black line) and sepiapterin (green line) are indicated.

**Supplemental Table S1**

| Variables | Controls n = 18 | PD Patients n = 18 | "p" values |
|---|---|---|---|
| **Characteristics of controls and PD patients** | | | |
| **Gender, n (%)** | | | |
| Male | 09 (39.1) | 14 (60.9) | |
| Female | 09 (69.2) | 04 (30.8) | 0.16 |
| **Race (%)** | | | |
| White | 16 (88.9) | 15 (83.3) | |
| Afro-Brazilians | 03 (11.1) | 02 (16.7) | 1.0 |
| **Age, years, Mean (SE)** | 61.4 (2.4) | 61.7 (2.3) | 0.93 |
| **Clinical comorbidities, n (%)** | | | |
| Ischemic heart disease | 01 (5.6) | 01 (5.6) | 1.0 |
| Atrial Fibrillation | 0 | 01 (5.6) | 1.0 |
| Systemic hypertension | 03 (16.7) | 06 (33.3) | 0.44 |
| Diabetes Mellitus type 2 | 02 (11.1) | 0 | 0.49 |
| Hypothyroidism | 0 | 01 (5.6) | 1.0 |
| Benign prostatic hyperplasia | 0 | 01 (5.6) | 1.0 |
| **Psychiatric comorbidities** | | | |
| Depression[a] | 0 | 10 (55.6) | < 0.0001 |
| Anxiety | 0 | 1 (5.6) | 1.0 |
| **Characteristics of PD patients** | | | |
| **Age at the diagnosis in years, Mean (SE)** | N.A. | 48.4 (2.1) | N.A. |
| **Time since diagnosis in years, Mean (SE)** | N.A. | 13.2 (1.2) | N.A. |
| **First PD symptom** | | | |

| | | | |
|---|---|---|---|
| Bradykinesia | N.A. | 05 (13.9) | N.A. |
| Tremor | N.A. | 07 (19.4) | N.A. |
| Rigidity | N.A. | 02 (5.6) | N.A. |
| Unknown | N.A. | 04 (11.2) | N.A. |
| **Pharmacological Treatment** | | | |
| L-Dopa only | N.A. | 05 (13.9) | N.A. |
| L-Dopa + Pramipexol | N.A. | 06 (33.3) | N.A. |
| L-Dopa + Amantadine | N.A. | 02 (5.6) | N.A. |
| L-Dopa + Pramipexol + Amantadine | N.A. | 05 (13.9) | N.A. |
| Daily Levodopa dose in mg/kg, Mean (SE) | N.A. | 14.1 (2.2) | N.A. |
| **Modified Hoehn and Yahr Scale** | N.A. | 4.4 (0.4) | N.A. |
| **Unified Parkinson's disease rating scale (UPDRS)** | N.A. | | |
| Part I – Non-Motor Aspects of Experience of Daily Living | N.A. | 13.6 (2.1) | N.A. |
| Part II - Motor Aspects of Experience of Daily Living | N.A. | 20.1 (2.7) | N.A. |
| Part III – Motor Examination | N.A. | 45.1 (5.0) | N.A. |
| Part IV – Motor Complications | N.A. | 6.8 (1.3) | N.A. |

[a] The controls and patients with depression were under treatment with serotonin reuptake inhibitors;

Categoric variables were analyzed by Fisher Exact test;

Continuous variables were analyzed by Student "t" test;

N.A. = Non-applied to controls.

**Supplemental Tables**

**Table S1. Clinical and demographic characteristics of PD patients from cohort 1**

| GCH1 mutation | Sex, Age at onset/presentation | Location | Notes | Reference(s) |
|---|---|---|---|---|
| **Dopamine-responsive dystonia** | | | | |
| c.631_632delAT (p.Met211ValfsTer38) | F, 2 | China | | Clinical and genetic heterogeneity in a cohort of Chinese children with dopa-responsive dystonia; Chen et al., Front. Pediatr 2020, 8: 83. doi: 10.3389/fped.2020.00083 |
| c.631_632delAT (p.Met211ValfsTer38) | M, 27 (father of above) | | P | |
| c.548A>C (p.E183A) | M, 5 | | | |
| c.350T>G (p.L117R) | F, Infant | China | | Dopa-responsive dystonia in Chinese patients: including a novel heterozygous mutation in the GCH1 gene with an intermediate phenotype and one case of prenatal diagnosis; Zhang W et al, Neurosci Lett 2017, 644: 48-54. doi: 10.1016/j.neulet.2017.01.019<br><br>A novel GTPCH deficiency mouse model exhibiting tetrahydrobiopterin-related metabolic disturbance and infancy-onset motor impairments; Jiang et al, Metabolism 2019, 94: 96-104. doi: 10.1016/j.metabol.2019.02.001 |
| c.551G>A (p.R184H) c.610G>A (p.V204I) Compound heterozygous | M, 16 months | India | Asymptomatic father and mother are carriers for R184H and V204I, respectively | A compound heterozygote for GCH1 mutation represents a case of atypical dopa-responsive dystonia; Giri S et al, J Mol Neurosci 2019, 68: 214-220. doi: 10.1007/s12031-019-01301-3 |
| c.738T>C (p.*251R) | NA, 41 | Denmark | Mild P | Heterozygous mutations in GTP-cyclohydrolase-1 reduce BH4 biosynthesis but not pain sensitivity; Nasser et al, Pain 2018, 159: 1012-1024. doi: 10.1097/j.pain.0000000000001175<br><br>The patients with the c.899T>C (p.*251R) mutation belong to the same family. |
| c.738T>C (p.*251R) | NA, 6 | | Mild P + D | |
| c.738T>C (p.*251R) | NA, 9 | | D | |
| c.738T>C (p.*251R) | NA, 15 | | Mild P | |
| c.738T>C (p.*251R) | NA, 6 | | Mild P | |
| c.738T>C (p.*251R) | NA, 8 | | No signs | |
| c.738T>C (p.*251R) | NA, 10 | | OI | |
| c.738T>C (p.*251R) | NA, 2.5 | | P + D | |
| c.738T>C (p.*251R) | NA, 10 | | Mild P | |
| c.738T>C (p.*251R) | NA, NA | | NS, OI | |
| c.738T>C (p.*251R) | NA, NA | | NS, OI | |
| Ex1del | NA, NA | | | |
| Ex1del | NA, NA | | NS | |
| Ex1del | NA, Childhood | | | |
| c.262delC (p.Q87fs) | NA, 16<br>NA, 16<br>NA, 10 | | | |
| c.268G>C (p.G90R) | NA, NA<br>NA, NA<br>NA, NA | | | |
| c.662_665delTGAA (p.M221Tfs*5) | NA, NA | | | |
| Ex1-6del (no protein) | NA, NA | | | |
| c.317C>T (p.T106I) | F, 8 | China | | Targeted gene capture sequencing in diagnosis of dystonia patients; Ma et al, J Neurol Sci 2018, 390: 36-41. doi: 10.1016/j.jns.2018.04.005 |
| c.608G>A (p.G203E) | F, 23 | | | |
| c.626+2T>G | F, 10 | Japan | | |
| c.544C>T (p.Q182X) | F, 10 | | | |

| | | | | |
|---|---|---|---|---|
| c.557C>A (p.T186K) | M, 19<br>F, 11<br>F, 10 | | | *GCH1* mutations in dopa-responsive dystonia and Parkinson's disease; Yoshino H et al, J Neurology 2018, 265: 1860–1870.<br>doi: 10.1007/s00415-018-8930-8<br><br>Deep brain stimulation shows high efficacy in two patients with *GCH1* variants; Daida et al, Parkinson Related Disord 2019, 65: 277-278.<br>doi: 10.1016/j.parkreldis.2019.06.002 |
| c.343+1G>T | F, 3 | | | |
| c.181G>T (p.E61X) | F, 34 | | | |
| c.509+2_3insT | F, 47 | | | |
| c.626+1G>C | F, 13 | | | |
| c.54C>A (p.C18X) | F, 8 | | | |
| c.706G>T (p.Glu236*) | F, 15<br>F, 19 | France | Mother and daughter | Mutation in the *GCH1* gene with dopa-responsive dystonia and phenotypic variability; Krim E et al, Neurol Genet 2018, 4: e231.<br>doi: 10.1212/NXG.0000000000000231 |
| c.283C>T (p.P95S), c.671A>G (p.K224R) Compound heterozygous | F, Infant | Spain | | Novel GCH1 compound heterozygosity mutation in infancy-onset generalized dystonia; Flotats-Bastardas M et al, Neuropediatrics 2018, 49: 296-297.<br>doi: 10.1055/s-0038-1626709 |
| c.679A>G (p.T227A) | M, infant? | China | | Dopa-responsive dystonia in Han Chinese patients: one novel heterozygous mutation in GTP cyclohydrolase 1 (GCH1) and three known mutations in TH; Yang K et al, Med Sci Monit 2018, 24: 751-757.<br>doi: 10.12659/msm.907288 |
| c.670 A>G (p.K224E) | M, 43 | Taiwan | No diurnal fluctuation; lacking response to levodopa | Atypical presentation of dopa-responsive dystonia in Taiwan; Weng YC, Wang CC, Wu YR. Brain Behav 2018, 8: e00906.<br>doi: 10.1002/brb3.906 |
| c.263G>T (p. R88L) | F, 15 | Taiwan | R88L also in father (dystonia) and asymptomatic sister | A novel missense mutation of the GTP cyclohydrolase 1 gene in a Taiwanese family with dopa-responsive dystonia: A case report; Yang C-C et al, Clin Neurol Neurosurg 2018, 165: 21-23.<br>doi: 10.1016/j.clineuro.2017.12.018 |
| c.209delA (p.N70Tfs*10) | F, 16<br>F, 2<br>M, 8 | Serbia | | *GCH1* mutations are common in Serbian patients with dystonia-parkinsonism: Challenging previously reported prevalence rates of DOPA-responsive dystonia; Dobričić V et al, Parkinsonism Related Disord 2017, 45: 81-84.<br>doi: 10.1016/j.parkreldis.2017.09.017 |
| c.228delG (p.S77Pfs*3) | F, 6<br>F, 10<br>M, 7 | | | |
| c.400delG (p.D134Tfs*2) | F, 12 | | | |
| c.470T>C (p.L157P) | All F, 13, 12, 7, 6, 20, 22 | | | |
| c.541 + 1G > C | F, 7 | | Intron 4 | |
| c.550C>T (p.R184C) | M, 45 | | 1 patient (Parkinsonism), 1 control | |
| c.571G>A (p.V191I) | NA, NA | | 1 control | |
| c.608G>A (p.G203E) | F, 15,<br>F, 7<br>M, 7 | | | |
| c.626 + 1G > A | M, 7 | | Intron 5 | |
| Single copy deletion of exon 1 | M, Infant | USA | Diurnal fluctuation of symptoms | Dopa-responsive dystonia: a male patient inherited a novel GCH1 deletion from an asymptomatic mother; Wang W et al 2020, J Mov Disord 13: 150-153.<br>doi: 10.14802/jmd.19069 |

| | | | | |
|---|---|---|---|---|
| c. 542T>G (p. V181G) | M, 27 | Japan | First amino acid of exon 5 | Guitarist's cramp as the initial manifestation of dopa-responsive dystonia with a novel heterozygous GCH1 mutation; Hasegawa T et al , F1000 Res 2021, 10:361. doi: 10.12688/f1000research.51433.1 |
| c.305T>A (p. M102K) | F, 19 | Ireland | Daily levodopa dose 900 mg | A case of GCH-1 mutation dopa-responsive dystonia requiring high doses of levodopa for treatment; Bradley M et al 2021, Tremor Other Hyperkinet Move 11:23 |
| c.604G>A (p.V202I) | F, Infant | China | Homozygous – very rare | Severe hypotonia without hyperphenylalaninemia caused by a homozygous *GCH1* variant: a case report and literature review; Chen Y et al, Front Genet 2022, 13: 929069. doi: 10.3389/fgene.2022.929069 |
| c.703C>G (p.R235G) | F, Infant | India | Developmental delay, irritability, oculogyric crisis | Disorders of tetrahydrobiopterin metabolism: experience from south India; Ray S et al, Metab Brain Dis 2022, 37:743-760. doi: 10.1007/s11011-021-00889-z |
| c.457C>T (p.H153Y) | M, 6 | | Dystonia | |
| **Isolated dystonia** | | | | |
| rs3759664 | | | Minor A allele associated with isolated limb dystonia | Association of *TOR1A* and *GCH1* polymorphisms with isolated dystonia in India; Giri S et al, J Mol Neurosci 2021, 71:325-337. doi: 10.1007/s12031-020-01653-1 |
| **Spastic paraplegia** | | | | |
| c.454-2A>G r.spl | F, 2<br>F, 3 | Netherlands | Mother and daughter | Autosomal dominant GCH1 mutations causing spastic paraplegia at disease onset; Wassenberg T et al, Parkinsonism Relat Disord 2020, 74: 12-15. doi: 10.1016/j.parkreldis.2020.03.019 |
| c.193del (p.Glu65fs) | F, 14 | | | |
| c.631_632del (p.Met211fs) | M, 7 | | | |
| c.229_246del (p.S77_L82del) | M, 4 (twin A<br>M, 9 (twin B) | Canada | Monozygotic twins (adopted – genetic heritage unknown) | GCH1 mutations in hereditary spastic paraplegia; Varghaei P et al, Clin Genet 2021, 100: 51-58. doi: 10.1111/cge.13955 |
| c.614T>A (p.V205E) | M, 6 | | French-Canadian | |
| c.607G>A (p.G203R) | F, 4 | France | 1 patient with severe hereditary spastic paraplegia | Heterozygous pathogenic variation in *GCH1* associated with treatable severe spastic tetraplegia; Ravel JM et al, Parkinsonism Relat Disord 2023, online ahead of print. doi: 10.1016/j.parkreldis.2023.105310 |
| **GCH1 deficiency** | | | | |
| c.703C>G (p.R235G) | F, Infant | India | | A novel GCH1 mutation in an Indian child with GTP cyclohydrolase deficiency; Gowda VK et al, Indian J Pediatrics 2019, 86: 752–753. https://doi.org/10.1007/s12098-019-02900-z |
| **Parkinson's disease** | | | | |
| c.745delA (p.R249Gfs*76) | NA, NA | China | 1 case E | GCH1 variants contribute to the risk and earlier age-at-onset of Parkinson's disease: a two-cohort case-control study; Pan et al, Translational Neurodegeneration 2020, 9: 31. https://doi.org/10.1186/s40035-020-00212-3 |
| c.726A>G (p.E242E) | NA, NA | | 1 case G | |
| c.693G>C (p.L231F) | NA, NA | | 1 control E | |
| c.678G>C (p.V226V) | NA, NA | | 1 case E | |

| | | | | |
|---|---|---|---|---|
| c.608G>A (p.G203E) | NA, NA | | 1 case E | 2 cohorts, labelled WES (age at onset ≤50 yrs or with a family history) and WGS (sporadic late-onset PD, AAO > 50 yrs). WES case n=1555, control n=2234. WGS case n=1962, control n=1279. In the WES cohort, a significant association of GCH1 coding variants with PD was found. In the cohort WGS, however, no association was observed for either all variants or deleterious variants. In Notes column to left, E=WES, G=WGS.<br><br>Among non-coding variants, "rs12323905 remained significantly associated with PD after multiple comparison correction in our data. … previously reported associated variants rs841 and GWAS signal rs11158026 were also found to be associated with PD in the same association direction, however, these associations did not remain significant after multiple comparison correction." |
| c.593G>A (p.R198Q) | NA, NA | | 1 case E | |
| c.582G>A (p.T194T) | NA, NA | | 1 case G | |
| c.579C>G (p.I193M | | | 2 cases E | |
| c.573A>G (p.V191V) | | | 3 controls E; 1 case + 1 control G | |
| c.562C>T (p.Q188X) | | | 1 case E | |
| c.552C>A (p.R184R) | | | 1 case E | |
| c.507G>A (p.A169A) | | | 1 control E | |
| c.458A>C (p.H153P) | | | 1 case E | |
| c.409A>G (p.M137V) | | | 1 case E | |
| c.362dupT (p.F122Ifs*1) | | | 1 case E | |
| c.304A>T (p.M102L) | | | 1 case E | |
| c.294G>T (p.A98A) | | | 2 cases + 1 control E; 1 control G | |
| c.259delC (p.Q87Sfs*29) | | | 1 case E | |
| c.256C>T (p.P86S) | | | 1 case G | |
| c.246G>T (p.L82L) | | | 1 case G | |
| c.239G>A (p.S80N) | | | 4 cases + 1 control E; 6 cases + 2 controls G | |
| c.230C>G (p.S77C) | | | 2 cases + 1 control E; 3 cases + 1 control G | |
| c.210C>A (p.N70K) | | | 2 controls G | |
| c.196C>T (p.L66L) | | | 1 control E; 2 cases + 3 controls G | |
| c.193G>T (p.E65X) | | | 1 case E | |
| c.170G>A (p.R57Q) | | | 1 case E | |
| c.166G>A (p.E56K) | | | 1 control G | |
| c.116C>T (p.P39L) | | | 1 case G | |
| c.111G>A (p.E37E) | | | 1 case G | |
| c.91G>A (p.G31R) | | | 1 control E; 1 control G | |
| c.87G>T (p.R29R) | | | 1 case E | |
| c.75G>C (p.R25R) | | | 1 case E | |
| c.68C>T (p.P23L) | | | 1 control E | |
| c.32A>C (p.E11A) | | | 1 case G | |
| c.221C>A (p.A74D) | F, 47 | Korea | | GCH-1 genetic variant may cause Parkinsonism by unmasking the subclinical nigral pathology; Shin JH et al, J Neurol 2020, 267: 1952-1959. doi: 10.1007/s00415-020-09788-2 |
| c.671A>G (p.K224R) | M, 58 | France, Canada (Quebec), USA (NY) | | Common and rare *GCH1* variants are associated with Parkinson's disease; Rudakou U et al, Neurobiology of Aging 2019, 73: 231.e1-231.e6. https://doi.org/10.1016/j.neurobiolaging.2018.09.008 |
| c.671A>G (p.K224R) | M, 64 | | | |
| c.671A>G (p.K224R) | NA, NA | | | |
| (p.R184C) | M, 48 | | | |
| c.662T>C (p.M221T) | F, 61 | | | |
| c.626+2T>G | F, 64 | Japan | | *GCH1* mutations in dopa-responsive dystonia and Parkinson's disease; Yoshino H et al, J Neurology 2018, 265: 1860–1870. doi: 10.1007/s00415-018-8930-8 |
| c.626+1_2insT | F, 59 | | | |
| c.626C>T (p.T209I) | F, 31 | | | |
| c.646C>T (p.R216X) | F, 15 | | | |
| c.344-5T>A | F, 8 | | | |

| | | | | |
|---|---|---|---|---|
| | | | | Deep brain stimulation shows high efficacy in two patients with *GCH1* variants; Daida K et al, Parkinson Related Disord 2019, 65: 277-278.<br>doi: 10.1016/j.parkreldis.2019.06.002 |
| c.239G > A (p.S80N) | M, 50 | China | | Study of GCH1 and TH genes in Chinese patients with Parkinson's disease;<br>Yan Y-P et al, Neurobiol Aging 2018, 68: 159.e3-159.e6.<br>doi: 10.1016/j.neurobiolaging.2018.02.004 |
| rs11158026 | | China | No association of rs11158026 with PD within cohort, but meta-analysis showed association | Association analyses of variants of *SIPA1L2*, *MIR4697*, *GCH1*, *VPS13C,* and *DDRGK1* with Parkinson's disease in East Asians; Zou M et al, Neurobiol Aging 2018, 68: 159.e7-159.e14.<br>doi: 10.1016/j.neurobiolaging.2018.03.005 |
| c.170G>A (p.R57Q) | F, 41 | China | | Rare *GCH1* heterozygous variants contributing to Parkinson's disease; Xu Q et al, Brain 2017, 140: e41.<br>doi: 10.1093/brain/awx110 |
| c.230C>G (p.S77C) | M, 60 | | | |
| c.239G>A (p.S80N) | F, 58<br>M, 56 | | | |
| c.409A>G (p.M137V) | F, 41 | | | |
| c.593G>A (p.R198Q) | F, 47 | | | |
| c.608G>A (p.G203E) | F, 35 | | | |
| c.695G>A (p.G232D) | M, 51 | | | |
| | | | | A meta-analysis of genome-wide association studies identifies 17 new Parkinson's disease risk loci; Chang D et al, Nature Genet 2017, 49: 1511-1516.<br>doi:10.1038/ng.3955 |
| rs11158026<br>(C/C or C/T or T/C or T/T) | | USA | T allele: higher PD risk, 5 years earlier onset | Aging modifies the effect of GCH1 RS11158026 on DAT uptake and Parkinson's disease clinical severity; Webb J and Willette AA, Neurobiol Aging 2017, 50: 39-46.<br>doi: 10.1016/j.neurobiolaging.2016.10.006 |
| rs11158026 | | China | No association found of rs11158026 with PD | Polymorphism in *MIR4697* but not *VPS13C, GCH1,* or *SIPA1L2* is associated with risk of Parkinson's disease in a Han Chinese population; Yang X et al, Neurosci Lett 2017, 650: 8-11.<br>doi:10.1016/j.neulet.2017.04.003 |
| | Mean age at onset 56.7 years | Norway and Sweden | No putatively pathogenic GCH1 mutations found | Low frequency of GCH1 and TH mutations in Parkinson's disease; Rengmark A et al, Parkinsonism Relat Disord 2016, 29: 109-111.<br>doi: 10.1016/j.parkreldis.2016.05.010 |
| rs11158026 | | Taiwan | rs11158026 minor allele (C allele) frequency significantly higher in PD cases than in controls | Association of GCH1 and MIR4697, but not SIPA1L2 and VPS13C polymorphisms, with Parkinson's disease in Taiwan;<br>Chen C-M et al, Neurobiol Aging 2016, 39: 221.e1-5.<br>doi: 10.1016/j.neurobiolaging.2015.12.016<br><br>"T" is minor allele in Caucasians but "C" is minor allele in Asians |
| rs11158026 | | China | rs11158026 does not influence the risk of developing PD | Association of four new candidate genetic variants with Parkinson's disease in a Han Chinese population; Wang L et al, Am J Med Genet B Neuropsychiatr Genet 2016, 171B: 342-347.<br>doi: 10.1002/ajmg.b.32410 |

| | | | | |
|---|---|---|---|---|
| None found | | Spain | No mutation carriers were found for *GCH1* | Analysis of the genetic variability in Parkinson's disease from Southern Spain; Bandrés-Ciga S et al, Neurobiol Aging 2016, 37: 210.e1-210.e5. doi: 10.1016/j.neurobiolaging.2015.09.020 |
| rs11158026 | | Iran | rs11158026 T allele significantly more prevalent among PD cases | *SIPA1L2*, *MIR4697*, *GCH1* and *VPS13C* loci and risk of Parkinson's diseases in Iranian population: A case-control study; Safaralizadeh T et al, J Neurol Sci 2016, 369: 1-4. doi: 10.1016/j.jns.2016.08.001 |
| **Alzheimer's disease** | | | | |
| rs72713460 | | China | | Identification of genetic risk factors in the Chinese population implicates a role of immune system in Alzheimer's disease pathogenesis; Zhou X et al, Proc Natl Acad Sci 2018, 115: 1697-1706. doi: 10.1073/pnas.1715554115 |
| **Schizophrenia spectrum** | | | | |
| rs10137071 G/A | | | n=85 A allele (22 F, 42 M); n =21 GG 7 F, 14 M) | Regulation of cortical and peripheral *GCH1* expression and biopterin levels in schizophrenia-spectrum disorders; Clelland JD et al, Psychiatry Research 2018, 262: 229-236. doi: 10.1016/j.psychres.2018.02.020 |

D = dystonia, NA = not available, NS = no symptoms, OI = only interviewed, P = Parkinsonism, PD = Parkinson's disease

**Table S2. Recently published mutations (2016-present) in GCH1 which have been associated with dopa-responsive dystonia (DRD) and/or Parkinson's disease (PD)**

Synonymous and missense GCH1 mutations associated with neurological conditions, reported in papers primarily focusing on DRD and PD, published since 2016.

**Supplemental Table S3.**

| Mutation | DDG_Dyna | DDG_ENCoMencom | DDS_vib_ENCoM | DDG_mCSM | DDG_SDM | DDG_DUET | Summary | |
|---|---|---|---|---|---|---|---|---|
| E56K | -0,832 | -0,051 | 0,064 | 0,24 | -0,58 | 0,465 | 4d, 2s | |
| R57Q | -0,219 | 0,054 | -0,067 | 0,055 | -0,52 | 0,129 | 3d, 3s | |
| R59G | -0,466 | -0,443 | 0,554 | -0,436 | 0,49 | -0,16 | 5d, 1s | 1s SDM |
| P69L | 0,929 | 0,113 | -0,141 | -0,602 | 1,57 | 0,15 | 2d, 4s | |
| N70K | -0,15 | -0,034 | 0,042 | -0,067 | 0,81 | 0,576 | 4d, 2s | |
| L71Q | -0,49 | -0,288 | 0,36 | -1,893 | -1,17 | -1,802 | 6d, 0s | |
| A73D | 0,113 | 0,024 | -0,03 | -0,615 | -1,1 | -0,408 | 4d, 2s | 2s Dyn, DDS_vib |
| A74D | 0,054 | 0,059 | -0,073 | -0,336 | -0,94 | -0,076 | 4d, 2s | |
| A74V | 0,33 | 0,081 | -0,102 | -0,412 | -1,03 | -0,268 | 4d, 2s | |
| Y75C | -1,154 | -0,575 | 0,719 | -1,483 | -0,15 | -1,26 | 6d, 0s | |
| Y75S | -2,208 | -0,95 | 1,188 | -2,309 | -1,55 | -2,335 | 6d, 0s | |
| S77C | 0,028 | -0,087 | 0,109 | -0,134 | 0,68 | 0,311 | 3d, 3s | |
| L79P | -1,699 | -0,587 | 0,733 | -1,487 | -3,14 | -1,954 | 6d, 0s | |
| S80N | 0,027 | 0,047 | -0,059 | -0,386 | 0,42 | 0,068 | 2d, 4s | |
| S81P | -0,307 | -0,106 | 0,132 | -0,308 | -0,78 | -0,217 | 6d, 0s | |
| L82P | 0,327 | 0,061 | -0,077 | -0,268 | -0,83 | -0,207 | 4d, 2s | |
| G83A | -1,615 | -0,131 | 0,163 | -0,182 | -1,96 | -0,425 | 6d, 0s | |
| P86S | 0,7 | 0,133 | -0,166 | -1,495 | 0,04 | -1,314 | 3d, 3s | |
| Q87P | -0,449 | -0,144 | 0,181 | 0,153 | -0,1 | 0,294 | 4d, 2s | 2s CSM, DUET |
| R88G | -0,768 | -0,303 | 0,379 | -0,891 | 0,49 | -0,699 | 5d, 1s | 1s SDM |
| R88L | -0,472 | -0,265 | 0,331 | 0,276 | 0,62 | 0,397 | 4d, 2s | |
| R88P | -0,207 | 0,032 | -0,04 | 0,015 | 0,03 | -0,003 | 3d, 3s | 3s DDS_vib, CSM, DUET |
| R88W | -0,543 | -0,177 | 0,222 | -0,106 | 0,09 | -0,356 | 5d, 1s | |
| G90R | -0,005 | -0,042 | 0,052 | -0,545 | 0,18 | -0,253 | 5d, 1s | |
| G90V | -0,037 | -0,098 | 0,123 | -0,416 | 0,39 | 0,006 | 4d, 2s | 2s SDM, DUET |
| L91R | -1,332 | -0,068 | 0,085 | -1,095 | -1,76 | -0,999 | 6d, 0s | |
| L91V | -1,172 | -0,161 | 0,201 | -1,529 | -3,1 | -1,952 | 6d, 0s | |
| T94K | 0,011 | -0,152 | 0,19 | -0,66 | 1,13 | 0,014 | 3d, 3s | 3s Dyn, SDM, DUET |
| T94M | 0,252 | -0,102 | 0,127 | 0,117 | 1,33 | 0,624 | 2d, 4s | |
| P95L | 1,108 | 0,349 | -0,436 | -0,539 | 2,12 | 0,319 | 2d, 4s | 2d ENCoM, CSM |
| P95R | 0,781 | 0,452 | -0,565 | -1,105 | 0,55 | -0,684 | 3d, 3s | |
| P95S | -0,633 | 0,241 | -0,301 | -2,086 | -0,59 | -1,875 | 5d, 1s | |
| A98V | -0,073 | 0,139 | -0,173 | -0,601 | -0,97 | -0,458 | 5d, 1s | |
| A99D | -0,542 | 0,362 | -0,452 | -2,27 | -3,15 | -2,616 | 5d, 1s | |
| A99P | -0,045 | 0,278 | -0,347 | -0,495 | -4,45 | -1,171 | 5d, 1s | |

| | | | | | | | | |
|---|---|---|---|---|---|---|---|---|
| S100L | 0,916 | 0,337 | -0,421 | -0,125 | 2,06 | 0,691 | 2d, 4s | |
| M102K | -0,039 | 0,004 | -0,005 | -0,86 | -0,19 | -0,318 | 5d, 1s | 1s DDS_vib |
| M102L | -0,334 | -0,1 | 0,124 | -1,127 | 0,83 | -0,425 | 5d, 1s | |
| M102R | 0,335 | 0,066 | -0,083 | -0,66 | 0,24 | -0,188 | 3d, 3s | 3s Dyn, DDS_vib, SDM |
| Q103H | -0,688 | -0,343 | 0,428 | -0,948 | 0,78 | -0,702 | 5d, 1s | 1s SDM |
| Q103P | -0,101 | -0,484 | 0,605 | 0,13 | -0,5 | 0,163 | 4d, 2s | 2s CSM, DUET |
| F104L | -0,346 | -0,333 | 0,416 | -0,902 | 0,05 | -0,844 | 5d, 1s | |
| T106I | 0,118 | 0,016 | -0,021 | 0,043 | 0,89 | 0,429 | 1d, 5s | |
| G108D | -1,245 | 0,4 | -0,5 | -2,563 | -1,91 | -2,654 | 5d, 1s | 1s DDS_vib |
| G108S | -0,928 | 0,503 | -0,628 | -1,894 | -2,02 | -1,946 | 4d, 2s | 2s ENCoMs |
| G108V | -1,501 | 0,534 | -0,667 | -0,683 | -0,14 | -0,339 | 4d, 2s | 2s ENCoMs |
| Q110E | 0,18 | 0,084 | -0,105 | -0,081 | 0,4 | 0,435 | 2d, 4s | |
| T112A | 0,597 | 0,273 | -0,341 | -0,6 | 0,17 | -0,482 | 3d, 3s | |
| D115N | 0,192 | 0,05 | -0,063 | 0,444 | -0,07 | 0,537 | 2d, 4s | |
| L117R | 0,038 | -0,368 | 0,46 | -0,276 | -0,81 | -0,137 | 5d, 1s | |
| A120S | -0,587 | 0,072 | -0,09 | -1,137 | -1,79 | -1,161 | 5d, 1s | |
| D134G | -0,373 | -0,243 | 0,304 | -0,815 | 2,51 | 0,065 | 4d, 2s | |
| D134N | -0,159 | -0,07 | 0,087 | -0,723 | 1,18 | -0,146 | 5d, 1s | 1s SDM |
| D134V | -0,675 | -0,077 | 0,096 | -0,019 | 0,71 | 0,468 | 4d, 2s | |
| I135K | -0,799 | -0,022 | 0,028 | -1,948 | -1,66 | -1,893 | 6d, 0s | |
| I135T | -1,673 | -0,174 | 0,217 | -2,256 | -1,94 | -2,312 | 6d, 0s | |
| M137R | 0,174 | -0,018 | 0,023 | -1,04 | -2,05 | -0,792 | 5d, 1s | |
| M137V | 0,522 | 0,046 | -0,058 | -0,904 | 0,73 | -0,196 | 3d, 3s | |
| C141R | 0,505 | 0,522 | -0,652 | -2,091 | -1,69 | -1,897 | 3d, 3s | 3s Dyn, ENCoMs |
| C141W | 0,443 | 0,902 | -1,128 | -1,667 | -2 | -1,778 | 3d, 3s | 3s Dyn, ENCoMs |
| H144P | -0,004 | -0,333 | 0,416 | 1,157 | -1,63 | 0,741 | 4d, 2s | 2s ENCoM |
| L145F | -0,527 | 0,01 | -0,013 | -0,977 | -0,33 | -1,081 | 5d, 1s | |
| P147Q | 0,483 | 0,102 | -0,127 | -1,27 | 0,99 | -0,612 | 3d, 3s | |
| V152D | -3,756 | -0,401 | 0,501 | -3,269 | -3,81 | -4,008 | 6d, 0s | |
| H153P | -0,807 | -0,838 | 1,047 | -1,427 | -2,54 | -1,906 | 6d, 0s | |
| H153R | -0,953 | 0,013 | -0,016 | -2,109 | -2,21 | -2,306 | 5d, 1s | |
| I154S | -3,769 | -0,484 | 0,605 | -3,314 | -4,65 | -3,664 | 6d, 0s | |
| I154V | -0,442 | -0,14 | 0,174 | -1,24 | -2,38 | -1,515 | 6d, 0s | |
| G155S | -0,481 | 0,35 | -0,438 | -1,391 | -2,42 | -1,479 | 5d, 1s | |
| L157P | -1,4 | -0,416 | 0,52 | -1,408 | -2,37 | -1,758 | 6d, 0s | |
| Q161P | 0,031 | -0,471 | 0,588 | -0,118 | -0,7 | -0,078 | 5d, 1s | 1s Dyn |
| L163R | -0,19 | 0,063 | -0,079 | -1,244 | -2,89 | -1,342 | 5d, 1s | 1s DDS_vib |
| R170S | -1,38 | -0,367 | 0,459 | -1,539 | -1,85 | -1,781 | 6d, 0s | |

| | | | | | | | | |
|---|---|---|---|---|---|---|---|---|
| Y175C | -0,563 | -0,369 | 0,461 | -1,916 | -0,56 | -1,786 | 6d, 0s | |
| S176T | 0,111 | 0,134 | -0,168 | -1,412 | -0,35 | -1,184 | 4d, 2s | 2s Dyn, DDS_vib |
| R178G | 0,083 | -0,494 | 0,618 | -2,503 | 0,38 | -2,134 | 4d, 2s | 2s Dyn, SDM |
| R178S | -0,558 | -0,406 | 0,507 | -2,44 | -1,36 | -2,6 | 6d, 0s | |
| Q180P | 0,121 | -0,615 | 0,768 | -0,738 | -1,56 | -0,949 | 5d, 1s | |
| Q180R | -0,596 | -0,157 | 0,196 | -0,983 | -0,93 | -0,79 | 6d, 0s | |
| V181I | 0,579 | 0,204 | -0,255 | -0,993 | 0,13 | -0,756 | 4d, 3s | |
| Q182E | -0,923 | -0,254 | 0,317 | -1,868 | 1,29 | -1,162 | 5d, 1s | |
| E183A | 0,125 | -0,123 | 0,153 | -0,483 | -0,41 | -0,3 | 5d, 1s | |
| E183K | 0,965 | 0,216 | -0,271 | 0,133 | -0,98 | 0,345 | 2d, 4s | |
| R184C | -0,48 | -0,609 | 0,762 | -0,865 | -0,72 | -0,888 | 6d, 0s | |
| R184H | -1,073 | -0,247 | 0,309 | -1,386 | 0,03 | -1,234 | 5d, 1s | |
| L185R | -2,145 | -0,254 | 0,317 | -2,01 | -2,88 | -2,07 | 6d, 0s | |
| T186I | -0,167 | 0,223 | -0,278 | -0,452 | 1,35 | 0,216 | 3d, 3s | 3s DDS_vib, SDM, DUET |
| T186K | -0,042 | 0,094 | -0,118 | -0,882 | -1,56 | -0,882 | 5d, 1s | |
| V191I | -0,234 | -0,083 | 0,103 | -0,555 | 0,27 | -0,097 | 5d, 1s | |
| I193M | 0,211 | 0,067 | -0,084 | -0,642 | -0,6 | -0,569 | 4d, 2s | |
| I193N | -2,057 | -0,777 | 0,971 | -2,695 | -2,6 | -2,84 | 6d, 0s | |
| A196D | 0,482 | 0,178 | -0,222 | -0,993 | -1,85 | -0,991 | 4d, 2s | 2s Dyn, DDS_vib |
| A196S | -0,792 | 0,062 | -0,078 | -1,174 | -2,37 | -1,281 | 5d, 1s | 1s DDS_vib |
| R198Q | -0,399 | -0,221 | 0,276 | 0,006 | -0,4 | 0,01 | 5d, 1s | |
| R198W | -0,72 | -0,267 | 0,334 | -0,015 | 0,44 | -0,19 | 5d, 1s | 1s SDM |
| P199A | -0,463 | -0,179 | 0,224 | -1,946 | 2,41 | -1,079 | 5d, 1s | |
| P199L | 0,541 | 0,177 | -0,222 | -0,794 | 3,2 | 0,152 | 2d, 4s | 2d ENCoM, CSM |
| P199S | -0,465 | -0,005 | 0,006 | -2,338 | 0,01 | -2,043 | 5d, 1s | |
| G201E | 1,027 | 1,461 | -1,827 | -1,048 | -2,35 | -1,213 | 3d, 3s | |
| G201R | 1,167 | 1,784 | -2,23 | -0,81 | -2,5 | -0,882 | 3d, 3s | 3s Dyn, ENCoMs |
| V202I | -0,315 | 0,155 | -0,194 | -1,053 | -0,6 | -0,847 | 5d, 1s | 1s DDS_vib |
| G203E | -0,135 | 0,823 | -1,029 | -1,229 | -2,13 | -1,282 | 4d, 2s | |
| G203R | -0,14 | 0,956 | -1,195 | -0,772 | -2,15 | -0,687 | 4d, 2s | |
| V204G | -2,977 | -0,702 | 0,877 | -2,968 | -3,2 | -3,535 | 6d, 0s | |
| V204I | 0,065 | 0,32 | -0,399 | -0,899 | -0,6 | -0,676 | 4d, 2s | |
| V205E | -1,206 | 0,007 | -0,009 | -2,571 | -1,99 | -2,701 | 5d, 1s | 1s DDS_vib |
| V205G | -2,727 | -0,707 | 0,884 | -2,791 | -2,93 | -3,312 | 6d, 0s | |
| V206A | -2,424 | -0,198 | 0,248 | -2,233 | -2,66 | -2,672 | 6d, 0s | |
| A208E | 0,504 | 0,623 | -0,779 | -2,622 | -2,29 | -2,817 | 3d, 3s | |
| T209I | 0,86 | 0,118 | -0,147 | -0,016 | 1,02 | 0,551 | 2d, 4s | |
| T209P | 0,458 | -0,174 | 0,218 | -0,269 | -1,79 | -0,501 | 5d, 1s | |

| | | | | | | | | |
|---|---|---|---|---|---|---|---|---|
| H210Q | -0,437 | -0,246 | 0,308 | -0,467 | -0,92 | -0,483 | 6d, 0s | |
| H210R | -0,394 | -0,189 | 0,236 | -1,008 | -0,18 | -0,825 | 6d, 0s | |
| M211I | 0,854 | 0,13 | -0,163 | -1,275 | 0,44 | -0,677 | 3d, 3s | |
| M211V | 0,38 | -0,059 | 0,073 | -1,504 | 0,14 | -0,998 | 4d, 2s | |
| M213T | -0,278 | -0,411 | 0,514 | -0,744 | -1,06 | -0,406 | 6d, 0s | |
| M213V | -0,455 | -0,296 | 0,37 | -0,871 | 0,06 | -0,343 | 5d, 1s | 1s SDM |
| G217V | 0,243 | 0,18 | -0,225 | -0,401 | -1,49 | -0,548 | 4d, 2s | |
| K220R | 0,011 | 0,197 | -0,246 | -0,199 | 0,01 | -0,033 | 3d, 3s | 3s Dyn, DDS_vib, SDM |
| M221T | -0,259 | -0,525 | 0,656 | 0,3 | -0,43 | 0,581 | 4d, 2s | |
| S223R | -0,335 | -0,011 | 0,014 | -0,733 | 1,05 | -0,299 | 5d, 1s | 1s SDM |
| K224E | -0,255 | -0,287 | 0,358 | -0,038 | 0,3 | 0,415 | 4d, 2s | |
| K224R | -0,352 | -0,182 | 0,228 | -0,169 | -0,15 | 0,207 | 5d, 1s | |
| V226A | -0,452 | -0,203 | 0,253 | -0,836 | -1,41 | -0,819 | 6d, 0s | |
| T227A | -0,453 | -0,234 | 0,292 | -0,856 | 0,29 | -0,524 | 5d, 1s | |
| M230I | -0,37 | -0,15 | 0,188 | -0,498 | 0,66 | 0,21 | 4d, 2s | |
| L231F | 1,106 | 0,179 | -0,224 | -1,155 | -0,32 | -1,145 | 4d, 2s | |
| G232D | -1,291 | -0,266 | 0,333 | -1,085 | -3 | -1,449 | 6d, 0s | |
| G232V | -0,113 | -0,11 | 0,138 | -0,406 | -1,52 | -0,523 | 6d, 0s | |
| F234S | -3,491 | -0,937 | 1,171 | -3,36 | -3,48 | -3,538 | 6d, 0s | |
| R235G | -0,316 | -0,481 | 0,602 | -0,724 | -1,18 | -0,872 | 6d, 0s | |
| R235W | -0,264 | -0,276 | 0,345 | -0,343 | -0,37 | -0,551 | 6d, 0s | |
| R241Q | -1,021 | -0,383 | 0,478 | -1,167 | -0,8 | -1,091 | 6d, 0s | |
| R241W | 0,416 | 0,077 | -0,097 | -0,739 | -0,2 | -0,774 | 4d, 2s | 2s Dyn, DDS_vib |
| R249S | 0,069 | 0,003 | -0,004 | -0,329 | -0,65 | -0,371 | 4d, 2s | 2s Dyn, DDS_vib |
| Average | -0,378814815 | -0,054762963 | 0,068377778 | -0,982059259 | -0,725481481 | -0,850977778 | | |
| Maximum | 1,167 | 1,784 | 1,188 | 1,157 | 3,2 | 0,741 | | |
| Minimum | -3,769 | -0,95 | -2,23 | -3,36 | -4,65 | -4,008 | | |

**Table S3. ΔΔG and ΔΔS prediction table for published mutations in GCH1 associated with PD and/or DRD.**

Structure-based predictions of change in free energy (**ΔΔ**G) and vibrational entropy (**ΔΔ**S) arising from GCH1 missense mutations associated with human disease. All calculations (six for each mutation) were conducted through the DynaMut web service using the structure of human GCH1 (PDB entry 1FB1) in which the N-terminal 54 residues are absent. Column A: GCH1 missense mutation. Columns B, C, E, F and G are the **ΔΔ**G values (kcal $mol^{-1}$) calculated by DynaMut, ENCoM, mCSM, SDM and DUET methods, respectively. Column C = predicted change in vibrational entropy (kcal $mol^{-1}$ $K^{-1}$) calculated by ENCoM. Column H: of the six calculations, the number of calculations that predict the mutation to be destabilizing (d) and the number that predict the mutation to be stabilizing (s); note that a predicted increase in molecule flexibility (positive predicted change in vibrational entropy) is treated as an indicator of destabilization. The average, maximum and minimum values of each calculation type are listed at the bottom of the respective columns. DDG_Dyna, Dynamut prediction; DDG_ENCoM, NMA-based prediction; DDS_vib_ENCoM, predicted change in vibrational entropy.

**Table S4. Differential gene expression of *Gch1*$^{+}$ positive versus *Gch1*$^{-}$ cells within ventral midbrain DAergic neurons.**

**Supplemental Videos**

**Supplemental Video S1. Hind limb dragging in AAV5-*Th-Cre* treated *Gch1*$^{flox/flox}$ mice.**

**Supplemental Video S2. Hind limb dragging in AAV5-*Th-Cre* treated *Gch1*$^{flox/flox}$ mice.**

**Supplemental Video S3. Hind limb dragging in AAV5-*Th-Cre* treated *Gch1*$^{flox/flox}$ mice.**

**Supplemental Video S4. Repetitive movement in AAV5-*Th-Cre* treated *Gch1*$^{flox/flox}$ mice.**

**Supplemental Video S5. Tremors in AAV5-*Th-Cre* treated *Gch1*$^{flox/flox}$ mice.**
